# Phylogenomics, Classification, and Lifestyle Evolution in Raft- and Nursery Web- Spiders (Araneae: Dolomedidae and Pisauridae)

**DOI:** 10.1101/2024.08.23.609317

**Authors:** Kuang-Ping Yu, Ren-Chung Cheng, Charles R. Haddad, Akio Tanikawa, Brogan L. Pett, Luis N. Piacentini, Peter Jäger, Ho Yin Yip, Yuya Suzuki, Arnaud Henrard, Christina J. Painting, Cor J. Vink, Eileen A. Hebets, Mark S. Harvey, Matjaž Kuntner

**Affiliations:** Department of Organisms and Ecosystems Research, National Institute of Biology, Večna pot 121, 1000 Ljubljana, Slovenia; Department of Biology, Biotechnical Faculty, University of Ljubljana, Večna pot 111, 1000 Ljubljana, Slovenia; Department of Life Sciences, National Chung Hsing University, No. 145, Xingda Rd., South Dist., 40227 Taichung, Taiwan; Research Center for Global Change Biology, National Chung Hsing University, No. 145, Xingda Rd., South Dist., 40227 Taichung, Taiwan; Department of Zoology and Entomology, University of the Free State, P.O. Box 339, 9300 Bloemfontein, South Africa; Laboratory of Biodiversity Science, School of Agriculture and Life Sciences, The University of Tokyo, Japan; SpiDiverse, Biodiversity Inventory for Conservation (BINCO), 3380 Walmersumstraat, Glabbeek, Belgium; Centre for Ecology and Conservation, University of Exeter, Penryn Campus, Penryn, Cornwall, UK; Arachnology Division, Argentine Museum of Natural Science “Bernardino Rivadavia” CONICET, Av. Angel Gallardo 470, C1405DJR, Buenos Aries, Argentina; Senckenberg Research Institute, Arachnology, Mertonstraße 17-21, 60325 Frankfurt am Main, Germany; Discovery & Education Department, Ocean Park Hong Kong, 180 Wong Chuk Hang Road, Aberdeen, Hong Kong; Tokushima Prefectural Museum, Mukoterayama, Tokushima City, Tokushima Prefecture (post code 770-8070), Japan; Royal Museum of Central Africa, Leuvensesteenweg 13, B-3080, Tervuren, Belgium; Te Aka Mātuatua School of Science, University of Waikato, New Zealand; Te Pūnaha Matatini, Centre of Research Excellence, New Zealand; Department of Pest-management and Conservation, Lincoln University, Lincoln, New Zealand; School of Biological Sciences, University of Nebraska-Lincoln, Lincoln, NE 68588, USA; Collections & Research, Western Australian Museum, 49 Kew Street, Welshpool, Western Australia 6106, Australia; School of Biological Sciences, University of Western Australia, Crawley, Western Australia 6009, Australia; Jovan Hadži Institute of Biology, ZRC SAZU, Novi trg 2, 1000 Ljubljana, Slovenia; Department of Entomology, National Museum of Natural History, Smithsonian Institution, 10th and Constitution, NW, Washington, DC 20560-0105, USA; State Key Laboratory of Biocatalysis and Enzyme Engineering, and Centre for Behavioural Ecology and Evolution, School of Life Sciences, Hubei University, Hubei, China

**Keywords:** Fishing spiders, semi-aquatic spiders, ultraconserved elements, ancestral state reconstruction, comparative analysis, taxon sampling bias

## Abstract

Pisauridae Simon, 1890 or “nursery web spiders” are a global and heterogenous assemblage of spider genera with diverse lifestyles, containing web builders and webless species, as well as terrestrial and semi-aquatic species, notably “fishing spiders”, genus *Dolomedes* Latreille, 1804. Incomplete, unresolved, or conflicting phylogenies have so far hampered testing for *Dolomedes* and pisaurid monophyly and evolution. Here, we broadly address these questions within a phylogenomic and comparative framework. Our goals are i) reconstruction of a robust phylogeny to test the monophyly of *Dolomedes* and Pisauridae and to amend *Dolomedes* classification; ii) estimation of evolutionary shifts and trends in lifestyles and capture webs; and iii) evaluation of hypotheses of morphological trait association with a semi-aquatic lifestyle. To this end we generate subgenomic data (ultraconserved elements or UCE) for 53 *Dolomedes* species and 28 pisaurid genera. We analyze these data using maximum likelihood, Bayesian, and multi-species coalescence approaches, as well as using two different phylogenetic time calibration methods, RelTime and MCMCtree. Consistent across analytical approaches, our phylogenies reject the monophyly of both Pisauridae and *Dolomedes*. “Pisaurid” genera fall into three clades: 1) Focal Clade I groups the majority, including *Pisaura* Simon, 1886, hence representing true pisaurids; 2) Focal Clade II = *Blandinia* Tonini et al., 2016 is sister to Trechaleidae Simon, 1890 and Lycosidae Sundevall, 1833; 3) Focal Clade III with fishing and raft spiders groups *Dolomedes*, *Megadolomedes* Davies and Raven, 1980, and *Ornodolomedes* Raven and Hebron, 2018 and is sister to Focal Clade II, Trechaleidae, and Lycosidae. Our taxonomy, based on complementary taxa and morphological evidence, resurrects Dolomedidae Simon, 1876 to include *Dolomedes* and the Oceanic genera *Bradystichus* Simon, 1884, *Megadolomedes*, *Caledomedes* Raven and Hebron, 2018, *Mangromedes* Raven and Hebron, 2018, *Ornodolomedes*, and *Tasmomedes* Raven and Hebron, 2018. Both RelTime and MCMCtree analyses yield comparable divergence estimations: Pisauridae origin is estimated between 29 and 40 Ma; *Blandinia* between 21 and 34 Ma; Dolomedidae between 10 and 17 Ma; and *Dolomedes* between 9 and 16 Ma. In order to avoid misleading significant correlations and/or over-resolved ancestral states, we performed taxon sampling bias correction in all evolutionary analyses. Evolutionary analyses reconstruct semi-aquatic lifestyle as ancestral to a large clade containing pisaurids, lycosids, trechaleids, *Blandinia*, and dolomedids, with several reversals to terrestrial lifestyle. Capture webs evolved at least three times, with reversals. Counter to expectation, the evolution of lifestyles and capture webs are independent. Although leg and tarsus lengths do not indicate lifestyles, semi-aquatic taxa are significantly larger than terrestrial ones. We explain this pattern with a biomechanical threshold over which surface tension can be broken while spiders forage under water. Our time-calibrated analyses indicate that the evolution of terrestrial and web-building lifestyles from semi-aquatic ancestors in Pisauridae coincided with cooling and drying climates in the mid-Miocene. We therefore hypothesize that climatic changes have acted as strong selection pressures toward lifestyle diversification.

## Introduction

Known as raft or fishing spiders, *Dolomedes* Latreille, 1804 (Figs. 1a–b) is a globally distributed genus with 105 species of large wandering spiders, currently under the family Pisauridae Simon, 1890 (World Spider Catalog 2024). *Dolomedes* species are well known for their semi-aquatic lifestyle as most of them inhabit a variety of waterbodies. Their ecology is accompanied by remarkable behaviors such as rafting on the water surface, diving under the surface (McAlister 1960; Deshefy 1981; Suter 1999), and, quite unusual for spiders, preying on aquatic vertebrates (Williams 1979; McCormick and Polis 1982; Bleckmann and Lotz 1987; Zimmermann and Spence 1989; Nyffeler and Pusey 2014; Baba et al. 2019; Nyffeler and Gibbons 2022). *Dolomedes* species are therefore popular models for studying semi-aquatic adaptations in arthropods, focusing on neurosensing of water ripples (Roland and Rovner 1983; Bleckmann and Barth 1984; Bleckmann et al. 1994; Suter 2003), locomotion on water (Suter and Wildman 1999; Suter and Gruenwald 2000a, 2000b; Stratton et al. 2004; Suter 2013), as well as the associated morphologies (e.g., appendage proportions, setae morphologies and densities, cuticle structures, and respiratory organs; see Suter et al. 2004; Stratton and Suter 2009; Stratton et al. 2004; see also Lapinski et al. 2015).

**FIGURE 1.**
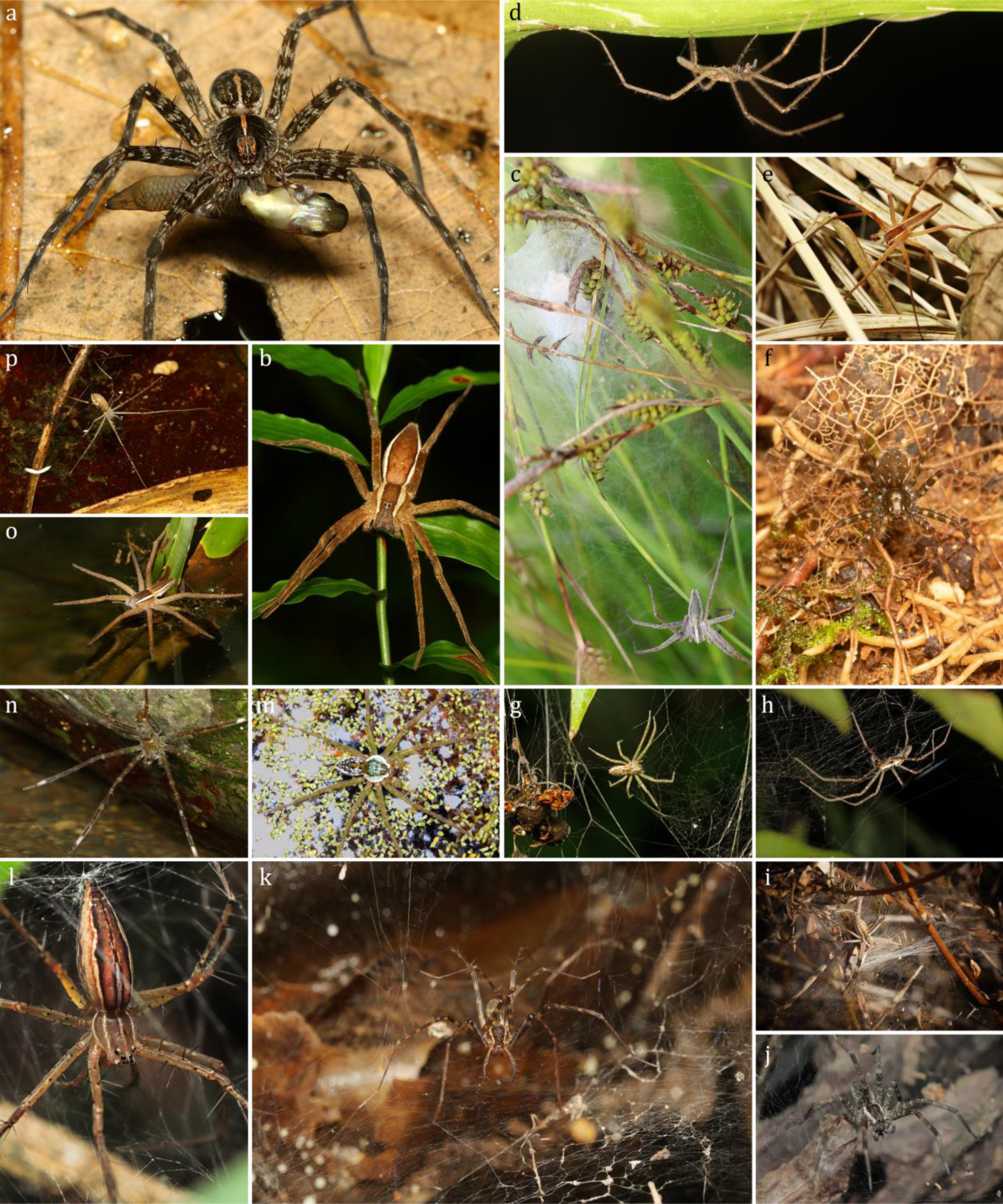
A glimpse into lifestyle diversity of the genera currently in Pisauridae Simon, 1890: (a) Female of semi-aquatic *Dolomedes mizhoanus* Kishida, 1936 preying on a fish, (b) Female of terrestrial *Dolomedes sulfureus* L. Koch, 1878 standing on vegetation; c–f, terrestrial lifestyle without capture web: (c) A female *Pisaura mirabilis* (Clerck, 1757) guarding her egg sac and nursery web, (d) Male of an unknown Pisauridae genus from Madagascar hanging under a leaf, (e) Male of *Perenethis fascigera* Bösenberg and Strand, 1906 roving on grassland litter layer (credit: Han-Po Chang), (f) Female of *Hala* sp. standing on the forest understory; g–l, terrestrial lifestyle with capture web: (g) Female of *Sphedanus quadrimaculatus* Thorell, 1897 hanging under her 3-dimensional web with retreat made of debris, (h) Female of *Caripetella madagascariensis* Lenz, 1886 hanging under her sheet web, (i) Juvenile of *Euprosthenopsis* sp. on its sheet web (credit: Matjaž Bedjanič), (j) Female of *Dendrolycosa* cf. *songi* (Zhang, 2000) standing on her sheet web (credit: Lok Ming Tang), (k) Female of *Blandinia mahasoana* (Blandin, 1979) standing on her sheet web, (l) Female of *Polyboea zonaformis* (Wang, 1993) standing in the center of her 3-dimentional web (credit: Lok Ming Tang); m–p, semi-aquatic lifestyle, (m) Subadult male of *Thaumasia velox* Simon, 1898 floating on water, (n) Female of *Qianlingula* cf. *turbinata* Zhang et al., 2004 in hunting posture, (o) Subadult female of *Nilus phipsoni* (F. O. Pickard-Cambridge, 1897) in hunting posture, (p) Female of *Hygropoda higenaga* (Kishida, 1936) in hunting posture.

Although most *Dolomedes* species strongly depend on freshwater-associated habitats (Carico 1973a; Zhang et al. 2004; Tanikawa and Miyashita 2008; Vink and Dupérré 2010; Raven and Hebron 2018; Yu and Kuntner 2024), not all species are semi-aquatic. For example, *D. sulfureus* L. Koch, 1878 (Fig. 1b), *D. silvicola* Tanikawa and Miyashita, 2008, *D. tenebrosus* Hentz, 1844, *D. albineus* Hentz, 1845, *D. minor* L. Koch, 1876, and *D. schauinslandi* Simon, 1899, are reported to be terrestrial, and largely independent of water bodies (Carico 1973a; Tanikawa and Miyashita 2008; Ono 2009; Vink and Dupérré 2010). Their distinct morphologies and disparate distributions in Asia, North America, and New Zealand suggest that these terrestrial species are not each other’s closest relatives. The currently understood lifestyle diversity within the genus raises questions of how, when, where, and why terrestrial versus semi-aquatic lifestyles evolved in *Dolomedes*. The lack of a comprehensive *Dolomedes* species phylogeny currently precludes answering these questions.

Similar questions related to the evolution of lifestyles can be asked more broadly about phylogenetic proximity of *Dolomedes*, i.e. to the family Pisauridae, whose constituent taxa display an even more complex array of lifestyle and trait diversity (Sierwald 1997; Zhang et al. 2004; Santos 2007a; Ono 2009; Jäger 2011; Raven and Hebron 2018; Dippenaar-Schoeman et al. 2020). With 52 genera and 365 species (World Spider Catalog 2024), this diverse global family is thought to be defined by the construction of nursery webs to protect their spiderlings (Fig. 1c). As adults, most pisaurid genera are terrestrial wandering spiders (Figs. 1b–f), others construct sheet or three-dimensional capture webs of numerous shapes and sizes in terrestrial habitats (Figs. 1g–l), and some are semi-aquatic (Figs. 1a, m–p) (Table 1). This phenotypic and ecological diversity makes pisaurids an ideal model group for studying the evolution of lifestyles and capture webs, yet their evolution has not been adequately analyzed due to unknown or conflicting phylogenies (Griswold 1993; Santos 2007b; Bayer and Schönhofer 2013; Moradmand et al. 2014; Polotow et al. 2015; Albo et al. 2017; Wheeler et al. 2017; Cheng and Piel 2018; Fernández et al. 2018; Piacentini and Ramírez 2019; Kallal et al. 2021; Hazzi and Hormiga 2023; Kulkarni et al. 2023) (Table 2; see also Taxonomy). Here, we address these issues within an original phylogenomic-evolutionary framework.

**TABLE 1.** The 52 genera currently listed in Pisauridae Simon 1890 and their known distribution ranges and lifestyles.

| Genus and author | Biogeographic realm | Lifestyle | Capture web | Data source | Include in this study |
| --- | --- | --- | --- | --- | --- |
| *Afropisaura* Blandin, 1979 | Afrotropic | terrestrial | absent | Dippenaar-Schoeman et al. 2020 | yes |
| *Archipirata* Simon, 1898 | Indomalayan | unknown | unknown |  | no |
| *Architis* Simon, 1898 | Neotropic | terrestrial | present | Santos 2007 | no |
| *Blandinia* Tonini et al., 2016 | Afrotropic | terrestrial | present | Yu original observation | yes |
| *Bradystichus* Simon, 1884 | Australasian | terrestrial | absent | Platnick and Forster 1993 | no |
| *Caledomedes* Raven and Hebron, 2018 | Australasian | terrestrial | absent | Raven and Hebron 2018 | no |
| *Caripetella* Strand, 1928 | Afrotropic | terrestrial | present | Yu original observation | yes |
| *Charminus* Thorell, 1899 | Afrotropic | terrestrial | absent | Dippenaar-Schoeman et al. 2020 | yes |
| *Chiasmopes* Pavesi, 1883 | Afrotropic | terrestrial | absent | Dippenaar-Schoeman et al. 2020 | no |
| *Cispinilus* Roewer, 1955 | Afrotropic | unknown | unknown |  | no |
| *Cispius* Simon, 1898 | Afrotropic | terrestrial | absent | Dippenaar-Schoeman et al. 2020 | yes |
| *Cladycnis* Simon, 1898 | Palaearctic | terrestrial | absent | Wunderlich 1987 | yes |
| *Conakrya* Schmidt, 1956 | Afrotropic | unknown | unknown |  | no |
| *Dendrolycosa* Doleschall, 1859 | Afrotropic & Australasian & Indomalayan | terrestrial | present | Jäger 2011; Raven and Hebron 2018 | yes |
| *Dolomedes* Latreille, 1804 | Global | semi-aquatic | absent | see main text | yes |
| *Eucamptopus* Pocock, 1900 | Indomalayan | unknown | unknown |  | no |
| *Euprostenops* Pocock, 1897 | Afrotropic | terrestrial | present | Dippenaar-Schoeman et al. 2020 | yes |
| *Euprostenopsis* Blandin, 1974 | Afrotropic | terrestrial | present | Dippenaar-Schoeman et al. 2020 | yes |
| *Hala* Jocqué, 1994 | Afrotropic | terrestrial | absent | Jocqué 1994; Yu original observation | yes |
| *Hygropoda* Thorell, 1895 | Afrotropic & Indomalayan | semi-aquatic | absent | Zhang et al. 2004; Dankittipakul et al. 2008 | yes |
| *Ilipula* Simon, 1903 | Indomalayan | unknown | unknown |  | no |
| *Inola* Davies, 1982 | Indomalayan | terrestrial | present | Davies 1982 | no |
| *Mangromedes* Raven and Hebron, 2018 | Australasian | semi-aquatic | absent | Raven and Hebron 2018 | no |
| *Maypaci* Simon, 1898 | Afrotropic | terrestrial | absent | Dippenaar-Schoeman et al. 2020 | yes |
| *Megadolomedes* Davies and Raven, 1980 | Australasian | semi-aquatic | absent | Raven and Hebron 2018 | yes |
| *Nilus* O. Pickard-Cambridge, 1876 | Afrotropic & Indomalayan | semi-aquatic | absent | Dippenaar-Schoeman et al. 2020 | yes |
| *Ornodolomedes* Raven and Hebron, 2018 | Australasian | terrestrial | absent | Raven and Hebron 2018 | yes |
| *Papakula* Strand, 1911 | Indomalayan | unknown | unknown |  | no |
| *Paracladynis* Blandin, 1979 | Afrotropic | terrestrial | absent | Yu original observation | yes |
| *Perenethis* L. Koch, 1878 | Afrotropic & Australasian & Indomalayan | terrestrial | absent | Sierwald 1997; Dippenaar-Schoeman et al. 2020 | yes |
| *Phalaeops* Roewer, 1955 | Afrotropic | unknown | unknown |  | no |
| *Pisaura* Simon, 1886 | Palaearctic | terrestrial | absent | Zhang et al. 2004 | yes |
| *Pisaurina* Simon, 1898 | Nearctic | terrestrial | absent | Carico 1973b | yes |
| *Polyboea* Thorell, 1895 | Indomalayan | terrestrial | present | Sierwald 1997 | yes |
| *Qianlingula* Zhang et al., 2004 | Indomalayan | semi-aquatic | absent | Yu original observation | yes |
| *Rothus* Simon, 1898 | Afrotropic | terrestrial | absent | Dippenaar-Schoeman et al. 2020 | yes |
| *Sphegulus* Thorell, 1877 | Indomalayan | terrestrial | present | Jäger 2011 | yes |
| *Stoliczka* O. Pickard-Cambridge, 1885 | Palaearctic | unknown | unknown |  | no |
| *Tallonia* Simon, 1889 | Afrotropic | unknown | unknown |  | yes |
| *Tapinothele* Simon, 1898 | Afrotropic | unknown | unknown |  | no |
| *Tapinothelella* Strand, 1909 | Afrotropic | unknown | unknown |  | no |
| *Tapinothelops* Roewer, 1955 | Afrotropic | unknown | unknown |  | no |
| *Tetragonophthalma* Karsch, 1878 | Afrotropic | terrestrial | present | Pett B. L. original observation; but see Sierwald 1997 | yes |
| *Thalassipsis* Roewer, 1955 | Afrotropic | unknown | unknown |  | no |
| *Thaumasia* Perty, 1833 | Neotropic | semi-aquatic | absent | Silva and Carico 2012 | yes |
| *Timus* F. O. Pickard-Cambridge, 1901 | Neotropic | semi-aquatic | absent | Carico 1976 | no |
| *Tolma* Jocqué, 1994 | Afrotropic | terrestrial | absent | Jocqué 1994; Yu<br>original<br>observation | yes |
| *Voraptipus* Roewer, 1955 | Afrotropic | unknown | unknown |  | no |
| *Voraptus* Simon, 1898 | Afrotropic &<br>Indomalayan | unknown | unknown |  | no |
| *Vuattouxia* Blandin, 1979 | Afrotropic | unknown | unknown |  | no |
| *Walrencea* Blandin, 1979 | Afrotropic | terrestrial | absent | Dippenaar-Schoeman et al.<br>2020 | no |

**TABLE 2.** Monophyly of Pisauridae Simon, 1890 and the affiliation of *Dolomedes* Latreille, 1804 discussed and analyzed in the previous literature.

| Literature | Evidence | Monophyly<br>of Pisauridae | Affiliation of<br>*Dolomedes* | Number of<br>genera<br>discussed |
| --- | --- | --- | --- | --- |
| Simon 1898b | eye arrangement | Yes | Dolomedae | 33 |
| Petrunkévitch 1928 | eye arrangement | Yes | Thaumasiinae | after Simon<br>1898b |
| Lehtinen 1967 | morphological<br>comparisons | No | Dolomedidae | 16 |
| Griswold 1993 | morphological<br>cladogram | Yes | sister to *Pisaura* | 2 |
| Sierwald 1990 | genital structure<br>homologous | Yes | closely related to<br>*Thalassius* (= *Nilus*) | 55 |
| Zhang et al. 2004 | morphological<br>cladogram | Yes | sister to *Thalassius*<br>(= *Nilus*) | 9 |
| Santos 2007b | morphological<br>cladogram | Yes | sister to *Pisaurina* | 8 |
| Bayer and Schönhofer<br>2013 | multi-genes phylogeny | No | sister to *Thalassius*<br>(= *Nilus*) | 3 |
| Moradmand et al. 2014 | multi-genes phylogeny | No | sister to Thalassinae | 2 |
| Polotow et al. 2015 | mutli-evidence<br>phylogeny | No | NA | 3 |
| Albo et al. 2017 | multi-genes phylogeny | No | separate clade, basal to<br>Lycosioidea | 5 |
| Wheeler et al. 2017 | multi-genes phylogeny | Yes | sister to *Bradystichus*,<br>basal pisaurid | 9 |
| Fernández et al. 2018 | transcriptomic<br>phylogeny | Yes | sister to *Pisaurina* | 2 |
| Cheng and Piel 2018 | transcriptomic<br>phylogeny | Yes | separate clade, basal<br>pisaurid | 4 |
| Piacentini and Ramírez<br>2019 | multi-genes phylogeny | Yes | sister to *Bradystichus*,<br>basal pisaurid | 9 |
| Kallal et al. 2021 | transcriptomic<br>phylogeny | Yes | separate clade, basal<br>pisaurid | 5 |
| Kulkarni et al. 2023 | UCE + multi-genes<br>phylogeny | No | separate clade, sister to<br>Lycosidae +<br>Trechaleidae | 13 |
| Hazzi and Hormiga 2023 | multi-genes phylogeny | No | separate clade, sister to<br>Lycosidae +<br>Trechaleidae | 7 |

Our first goal is to test the monophyly of *Dolomedes* and Pisauridae family, as well as the phylogenetic position of *Dolomedes*, using phylogenomic data, and to use these results to update the classification of *Dolomedes*. Although the monophyly of Pisauridae and the affiliation of *Dolomedes* have been tested multiple times under different frameworks (see citations above), none of the recent studies covered more than a quarter of the 52 pisaurid genera (Table 2). Limited taxon coverage has also led to an unclear topology within *Dolomedes,* the most speciose “pisaurid” genus (World Spider Catalog 2024). So far, *Dolomedes* species phylogenies have been regional and based on a few Sanger loci (Vink and Dupérré 2010; Tanikawa 2012; Yu and Kuntner, 2024). For example, a species level phylogeny of *Dolomedes* used a single gene fragment for 13 Japanese species (Tanikawa 2012) and failed to resolve all the nodes. Considering that Raven and Hebron’s (2018) taxonomic review of Australian pisaurids increased the number of *Dolomedes*’ close relatives from two to six genera with different degrees of intra- and intergeneric morphological variation, a better sampled *Dolomedes* phylogeny is also needed to test the boundaries of this diverse genus. We therefore set about to reconstruct a robust and well-sampled global phylogeny based on ultraconserved elements (UCE; Faircloth et al. 2012, 2013, 2015; Faircloth 2017; Miles Zhang et al. 2019) data and including more than half of the *Dolomedes* species as well as other genera currently in Pisauridae. To test the robustness of the phylogenetic results, we utilize maximum likelihood, Bayesian, and multi-species coalescence approaches. In order to add a historic time component to the evolution of the target clades, we also subject the phylogenomic data to two different time calibration methods, *a posteriori* (RelTime; Tamura et al. 2012; 2018) and *a priori* (MCMCtree; Yang 2007) (see Beavan et al. 2020).

Our second goal is to utilize the newly obtained phylogeny to reconstruct the evolution of lifestyles in *Dolomedes* and Pisauridae. To this end, we first populate the data for the terminals in the phylogeny on semi-aquatic vs. terrestrial lifestyles as well as the presence/absence of capture webs. Next, we test: i) whether a semi-aquatic lifestyle is ancestral in *Dolomedes* but not in Pisauridae, as suggested by a prior phylogenetic study of Lycosoidea (Piacentini and Ramírez 2019); ii) whether the absence of a capture web is ancestral in *Dolomedes* and Pisauridae (Piacentini and Ramírez 2019); and iii) if lifestyles and capture webs are correlated traits. This third, original hypothesis, predicts that evolution of a semi-aquatic lifestyle excludes the presence of a capture web.

Our third goal is to test the hypotheses that body size, leg length, and tarsus length all relate to a semi-aquatic lifestyle. Although researchers have analyzed how spider morphologies differ across foraging guilds on the spider tree of life (Wolff et al. 2022), semi-aquatic lifestyle was not considered as a separate guild and was therefore not discussed. To fill this knowledge gap, we test the correlation between a semi-aquatic lifestyle and morphological traits on the phylogeny. Although there are no rigorous empirical comparisons, the literature suggests that several semi-aquatic spiders exhibit iv) larger body sizes (*Ancylometes* Bertkau 1880, see Lapinski et al. 2015); v) relatively shorter legs (Tanikawa and Miyashita 2008); and vi) relatively longer tarsi (Lapinski et al. 2015) compared to terrestrial species (non-pisaurid species). We test these hypotheses on the phylogeny of *Dolomedes* and Pisauridae.

## Materials & Methods

### Phylogenetic taxon sampling

Phylogenetic taxon sampling comprised 96 terminals, including 53 *Dolomedes* morphospecies and 31 pisaurid species of 28 genera (of which one is without a certain genus affiliation, Fig. 1d) as ingroups (Table 3). Each pisaurid genus was represented by only one species, except for the diverse genera occupying more than one biogeographic realm: *Dendrolycosa* Doleschall, 1859, *Nilus* O. Pickard-Cambridge, 1876, and *Hygropoda* Thorell, 1895 were thus each represented by two species, one per biogeographic realm (Table 3). As outgroups to Pisauridae and *Dolomedes*, we used representatives of Trechaleidae Simon, 1890, Lycosidae Sundevall, 1833, Zoropsidae Bertkau, 1882, Psechridae Simon, 1890, Oxyopidae Thorell, 1869, Thomisidae Sundevall, 1833, and Viridasiidae Lehtinen, 1967 (Table 3).

**TABLE 3.**
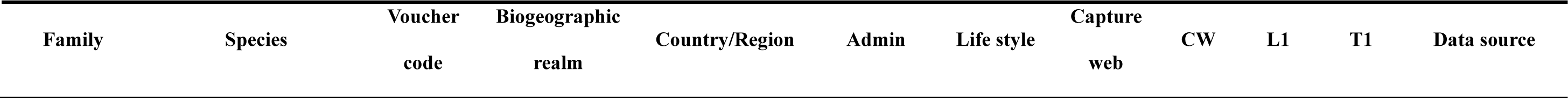
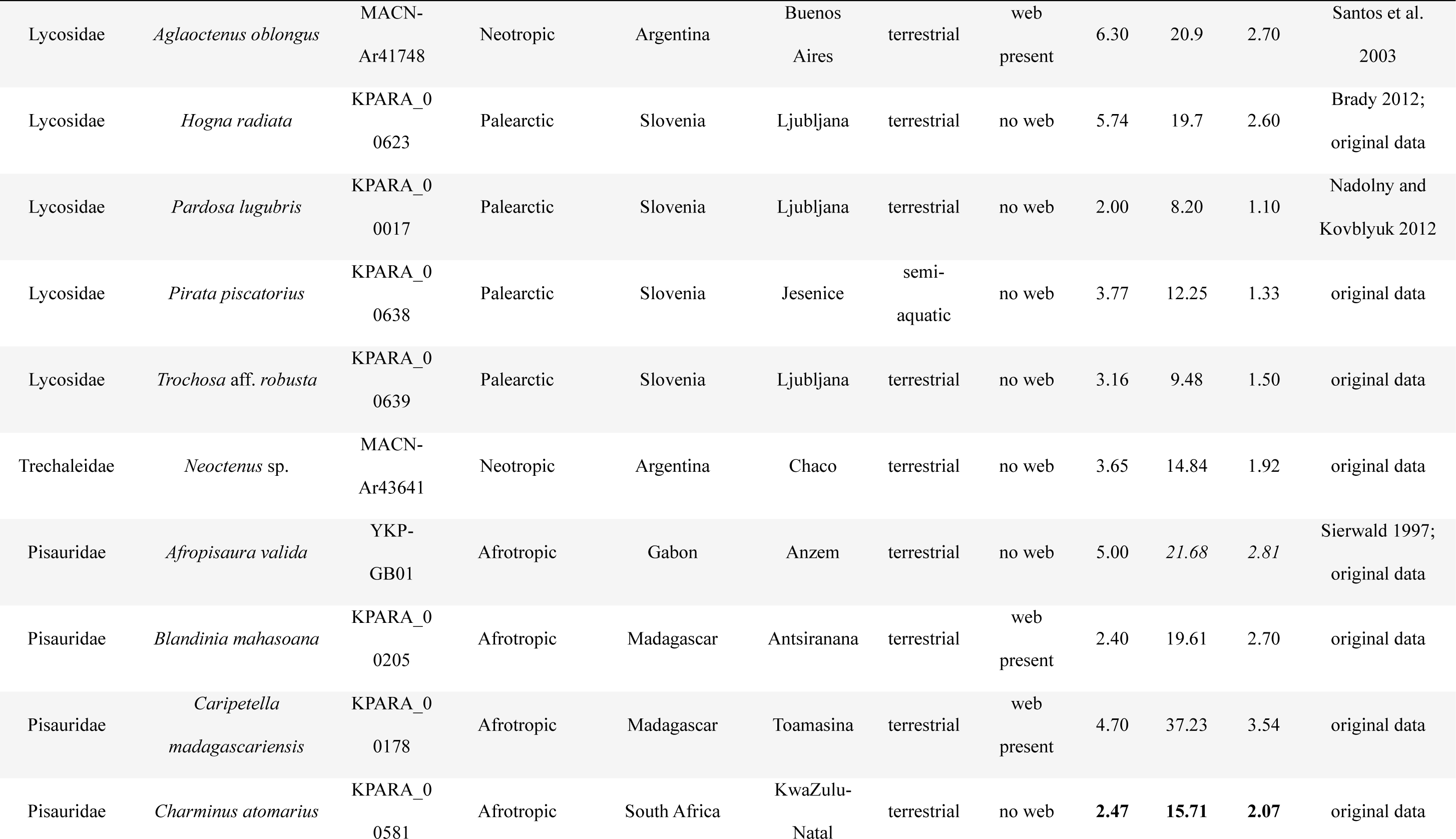
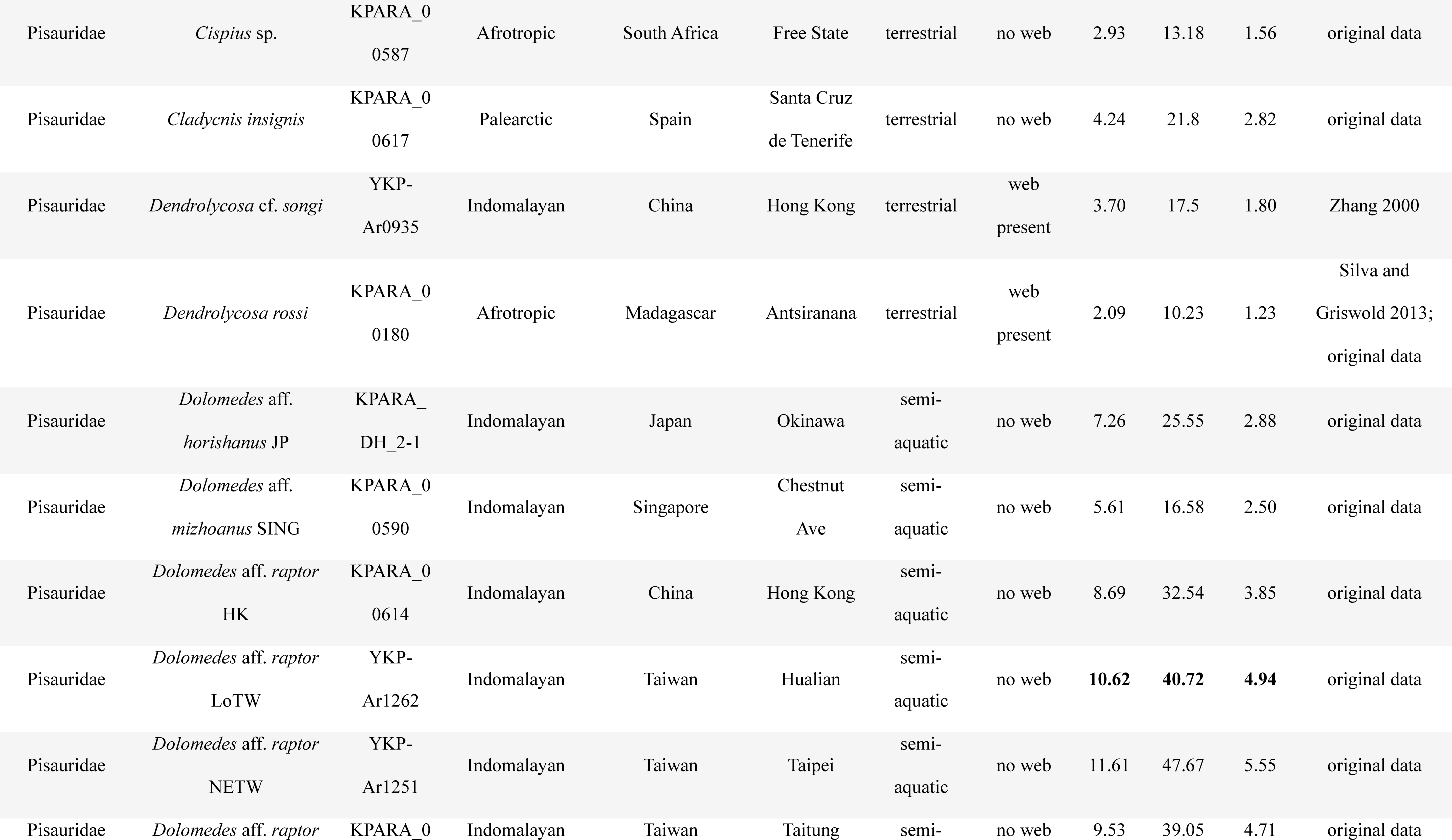
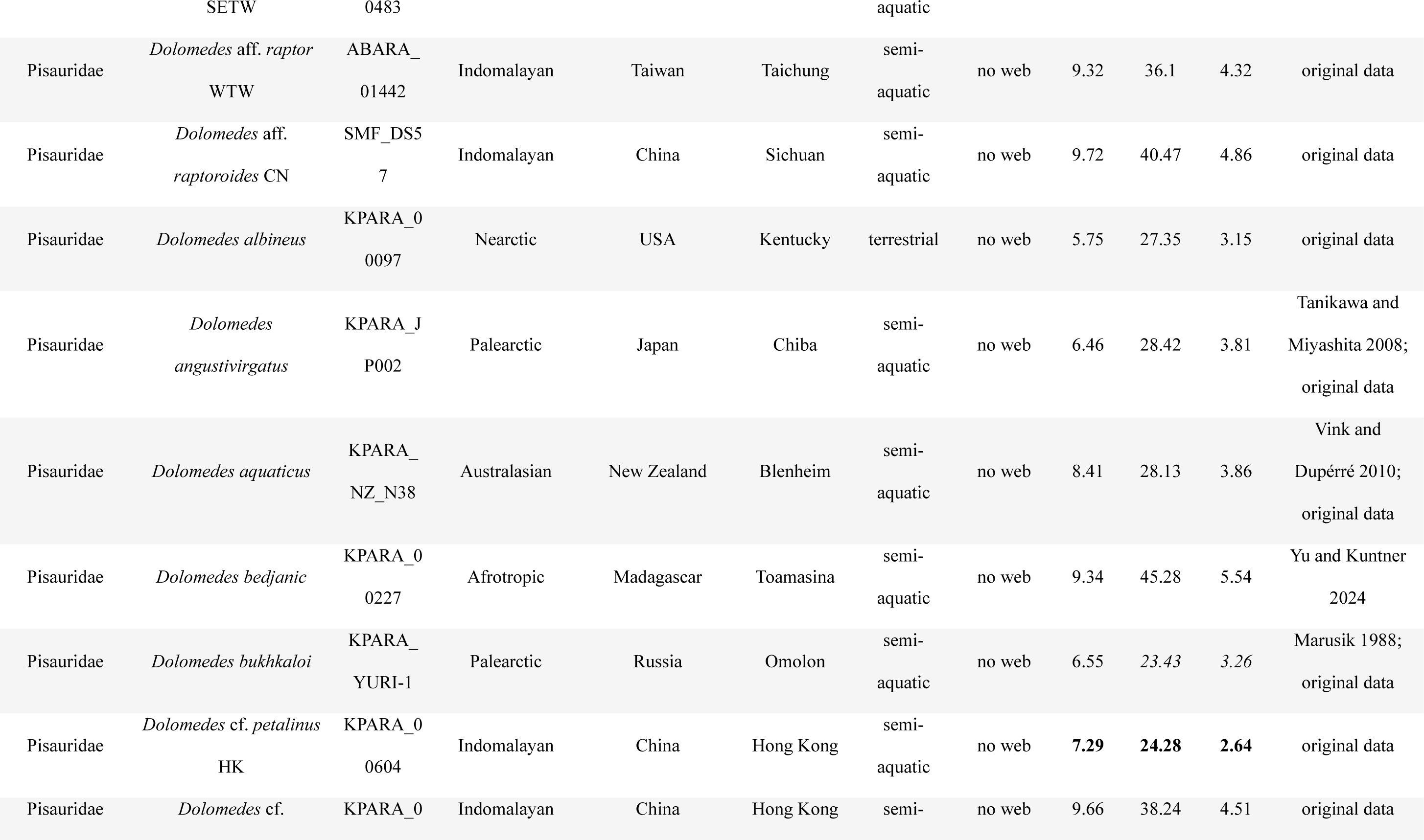
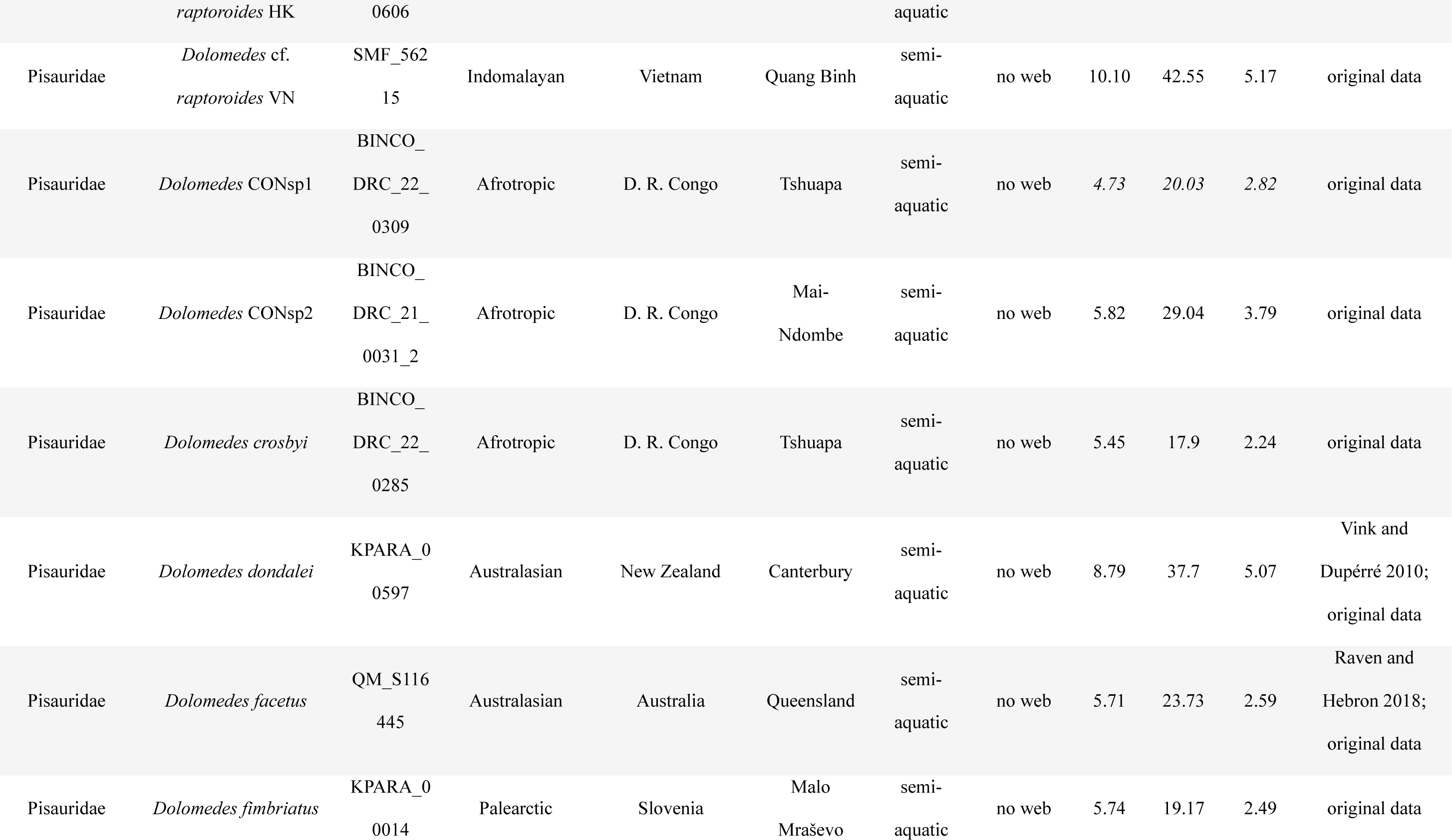
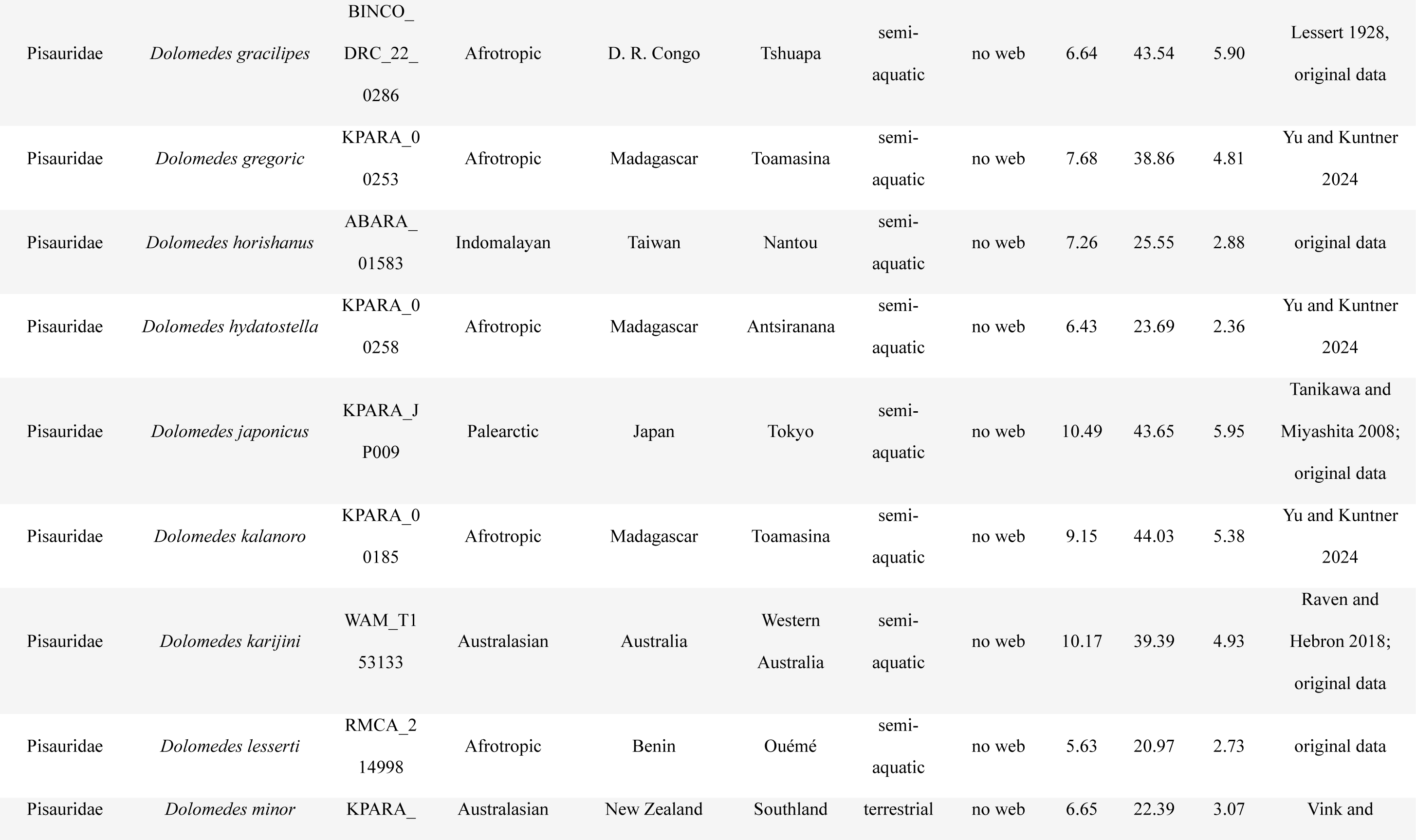
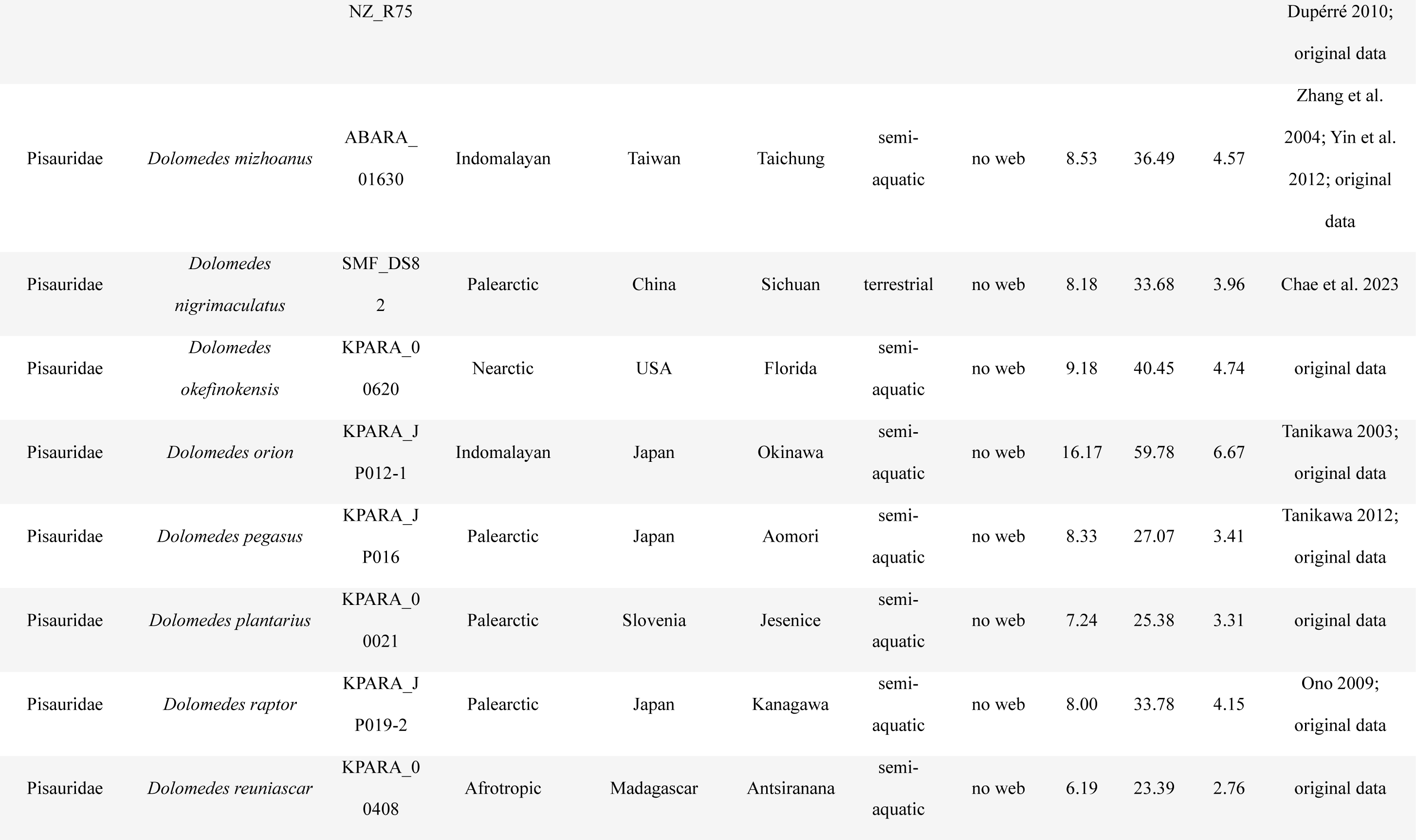
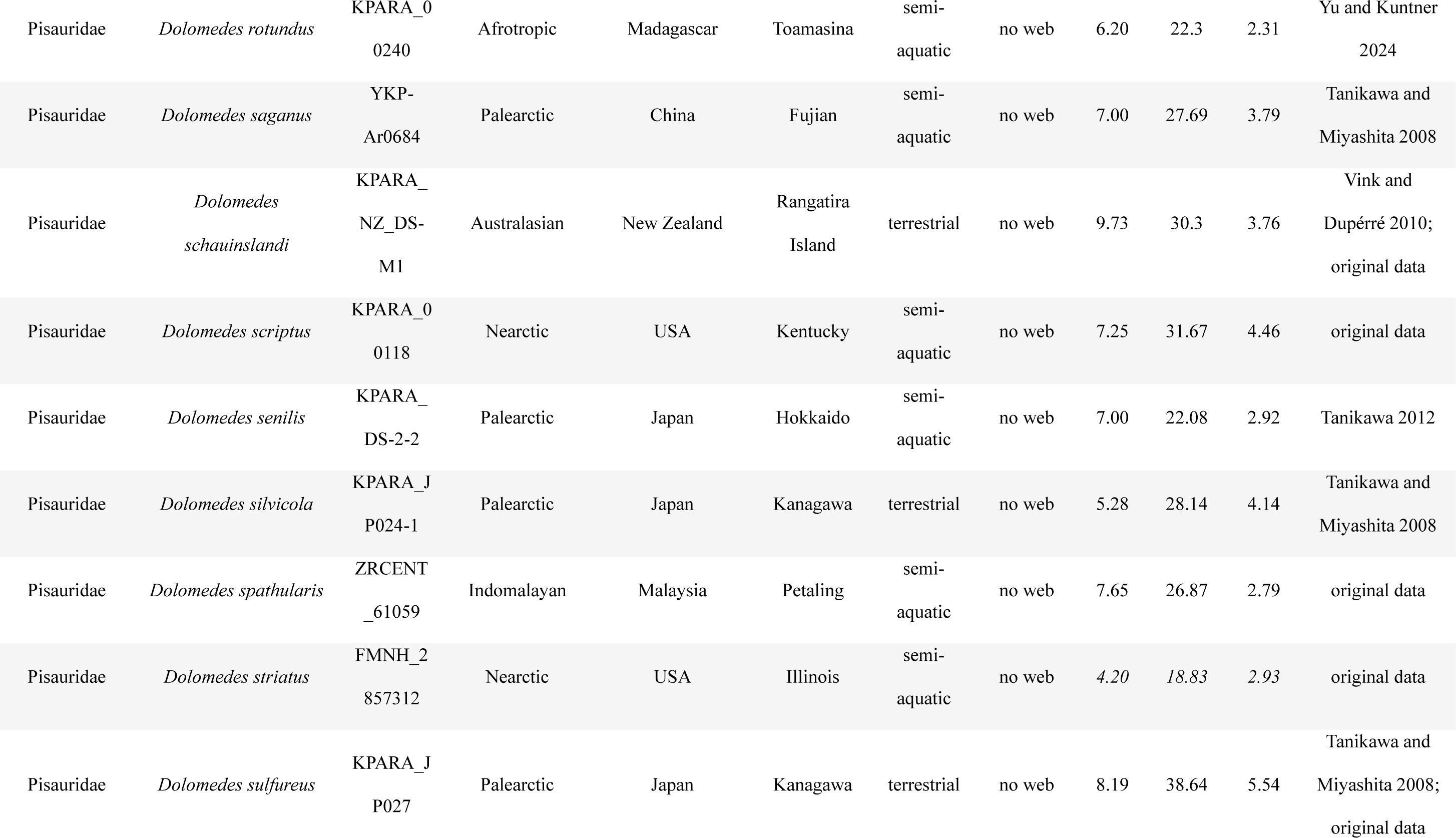
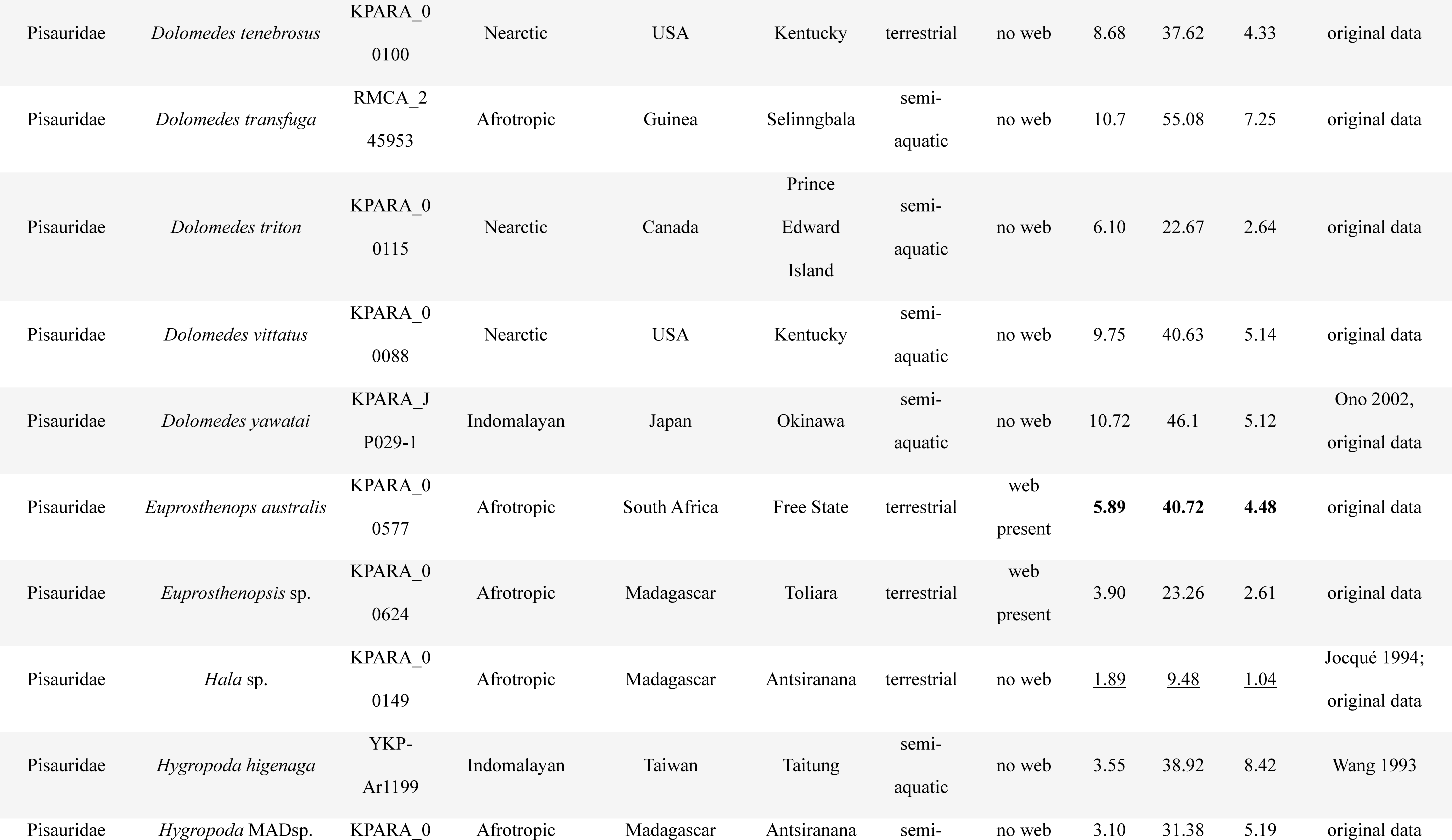
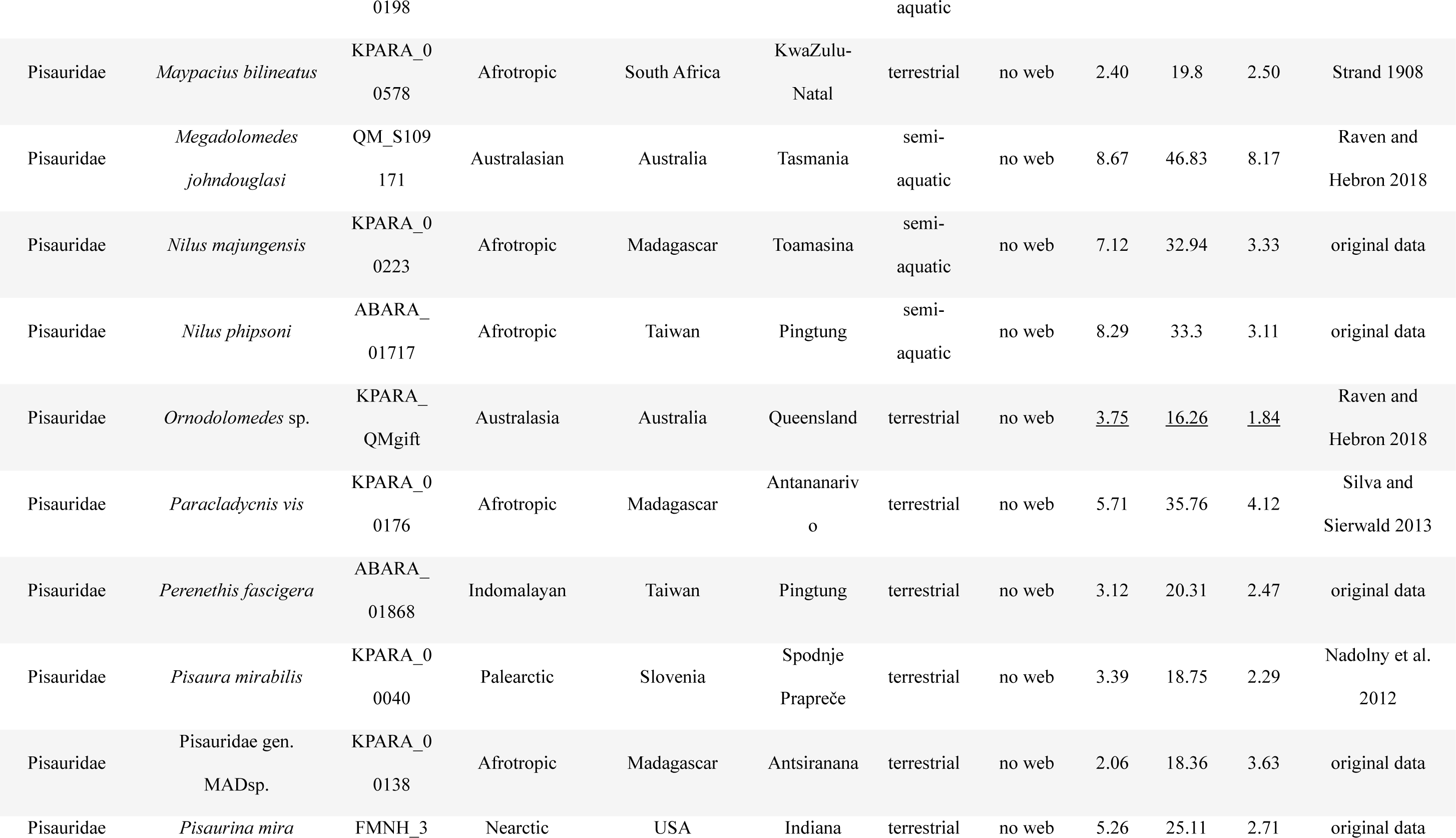
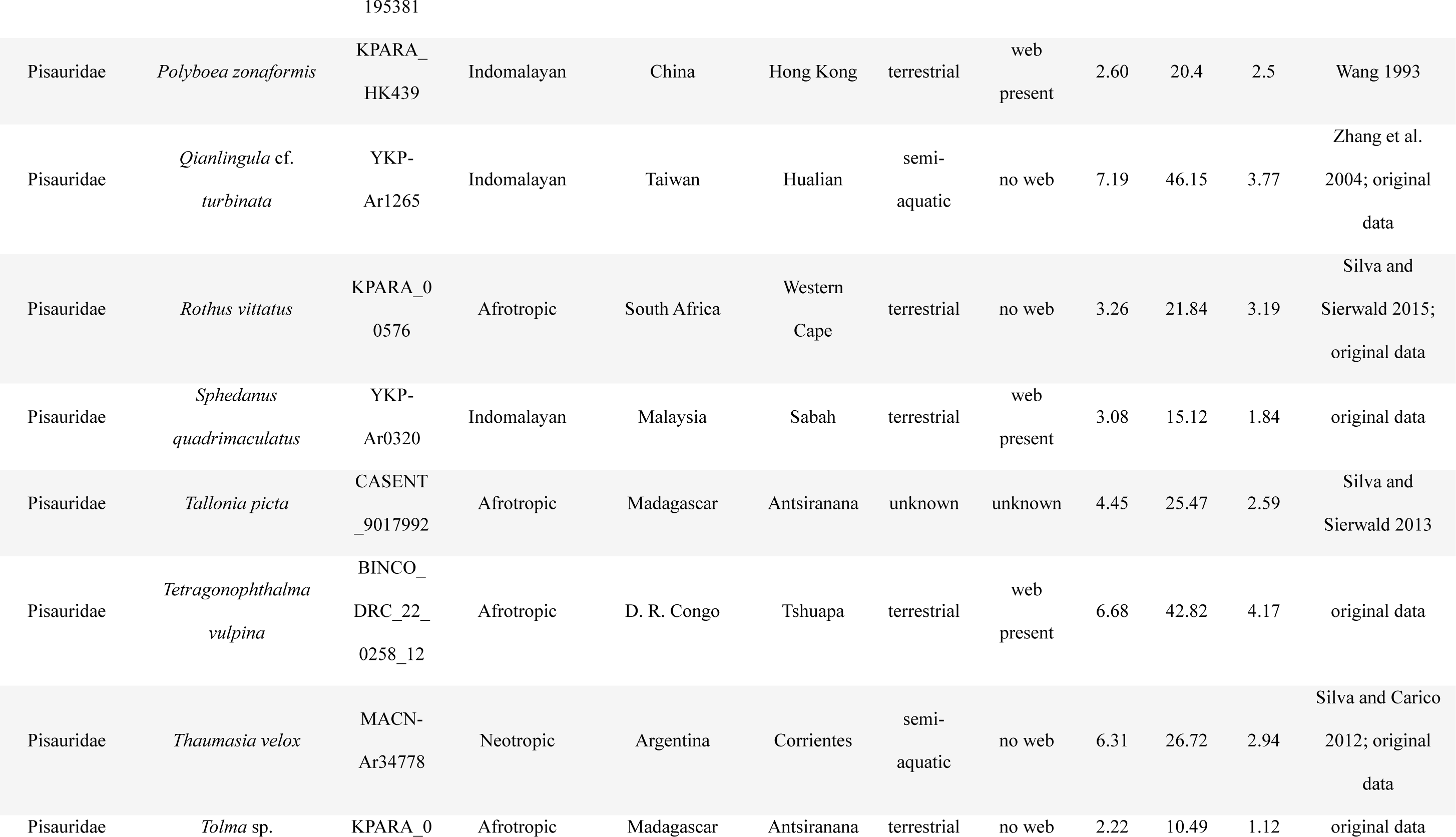
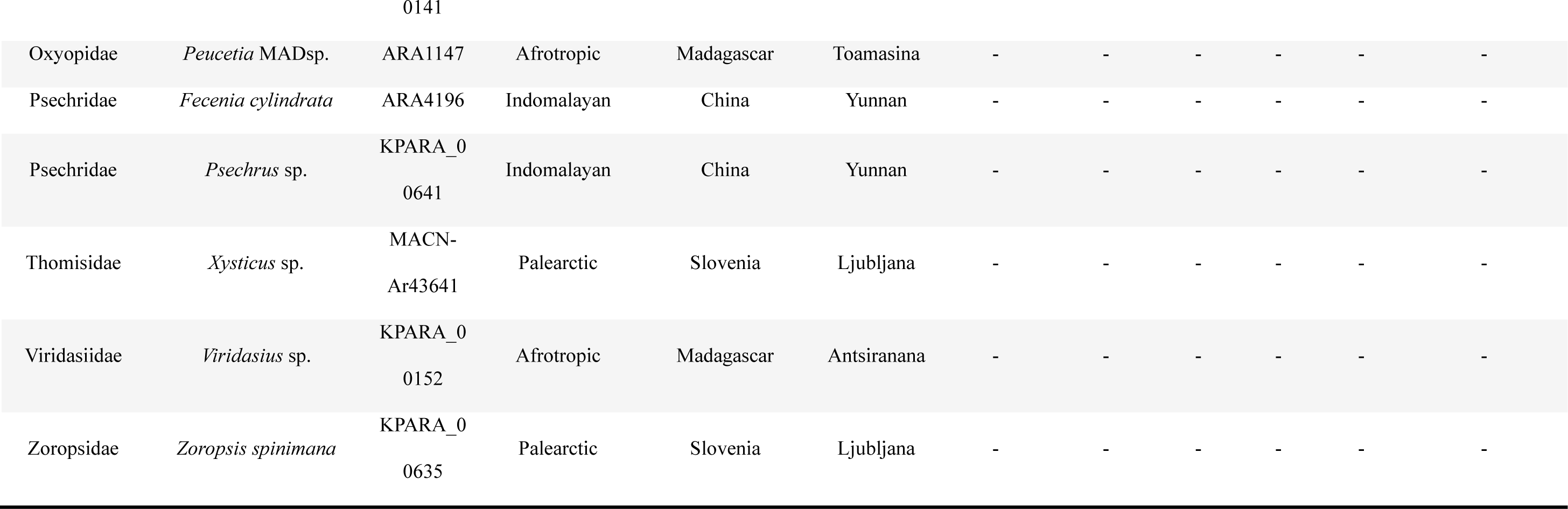
Raw data matrix of all the specimens used in the study, showing the region where they were collected and their traits used in the analyses. All measurements are averages from female specimens. If no female data or specimen was available, we then used data from males (italic numbers). If the terminals were represented by juveniles lacking species-level identification, the averages of the female of their genus were used (numbers underlined). Measurements or data from a single specimen are in bold. All measurements are in millimeters and the biogeographic realms were modified from Udvardy (1975). CW: carapace width; L1: leg I length; T1: tarsus I length; “-”: outgroups that are not in question. Voucher code and deposition: ABARA, YKP: National Chung Hsing University, Taiwan; ARA, KPARA: National Institute of Biology, Slovenia; BINCO: Biodiversity Inventory for Conservation, Belgium; CASENT: California Academy of Sciences, USA; FMNH: Field Museum, USA; MACN: Bernardino Rivadavia Natural Sciences Argentine Museum, Argentina; QM: Queensland Museum, Australia; RMCA: Royal Museum of Central Africa, Belgium; SMF: Senckenberg Natural History Museum, Germany; WAM: Western Australian Museum, Australia; ZRCENT: Lee Kong Chian Natural History Museum, Singapore.

### Lifestyle, capture web, and morphological data

We scored lifestyle and capture web as binary characters: semi-aquatic versus terrestrial lifestyle, and capture web presence versus absence (Table 3). A semi-aquatic lifestyle in our study refers to the adult spiders being documented to inhabit any water bodies in direct contact with the water. In the absence of such documentation, species are taken to be terrestrial. The presence of a capture web was only scored such in those species for which literature documents a prey capturing web (not merely webbing, nursery web, or nest; see Eberhard 2020) in adults. In the absence of such information for adults, species were scored as webless. We obtained data on carapace width (CW), leg I length (L1), and tarsus I length (T1), either by microscopic measuring or from the literature. We used CW to represent the size of a spider while relative leg and tarsus lengths were calculated by dividing L1 by CW and T1 by L1, respectively. We emphasized the measurement of adult females, but in the four cases where females were not available, we measured males instead (Table 3). This introduces little bias, as most pisaurids are not extremely sexually size dimorphic (Kuntner and Coddington 2020). Measurements were taken using a Leica M205C stereomicroscope or a Keyence VHX7000 digital microscope. For the two genera whose phylogenetic terminals were represented by juveniles and therefore lacked precise species identification, we averaged female measurements from the literature (Table 3).

### DNA extraction and phylogenomic pipeline

We used one to four legs from each specimen for DNA extraction. For small spiders, we also used the whole prosoma. We extracted genomic DNA using QIAamp DNA Micro Kits (Qiagen, Hilden, Germany), with the DNA quality checked using electrophoresis and a Qubit™ 4 Fluorometer (Invitrogen, Carlsbad, CA, USA) with the dsDNA HS Assay Kit. DNA samples containing more than 320 ng of DNA were then diluted to equal volumes and sent to Rapid Genomics LLC (Gainesville, FL, USA) for library preparation to capture and sequence UCE data. We used the RTA spider probe set (Zhang et al. 2023) for capturing UCE loci.

To assemble and filter the UCE matrix we employed the Phyluce v1.7.3 pipeline (Faircloth 2016) following the online tutorial (https://phyluce.readthedocs.io/en/latest/tutorials/index.html). After trimming the adaptors, we assembled the data using the package’s command lines under the SPAdes de novo assembly algorithm (Prjibelski et al. 2020). UCE loci were then found and extracted by matching the assembled contigs to the RTA spider probe set (Zhang et al. 2023). We ran both edges and internal trimming on each captured locus before aligning them. Finally, we aligned the captured loci with 75% and 95% taxon completeness into a sparse and a dense matrix, respectively, for subsequent phylogeny reconstruction. Both data matrices were uploaded to DRYAD.

### Phylogeny reconstruction

We reconstructed the phylogenies on both sparse and dense matrices using maximum likelihood (ML), Bayesian inference (BI), and multi-species coalescence (MSC) approaches. We chose the topology that received overall the highest nodal supports as the best tree and mapped the statistics from all these analyses onto it.

To partition both the sparse and the dense UCE matrices, we first split each locus into three fragments, a core region, a right, and a left flank, using entropy-based Sliding-Window Site Characteristics (SWSC-EN; Tagliacollo and Lanfear 2018). We then used PartionFinder 2 (Lanfear et al. 2017) to reclassify all fragments into the best-fitted partition subsets using AICc algorithm with the substitution models, suggested by SWSC-EN, set as GTR+G.

We preformed ML analyses in IQ-TREE v2.3.2 (Minh et al. 2020) using the partitioned matrices. Nodal supports were derived from ultrafast bootstrap approximation (UFBoot; Minh et al. 2013; Hoang et al. 2017) and SH-like approximate likelihood ratio test (SH-aLRT; Guindon et al. 2010), both with 1,000 replicates.

We carried out the BI analyses using the unpartitioned matrices in Exabayes 1.5.1 (Aberer et al. 2014) with default settings. On each matrix, we ran the Markov chain Monte Carlo (MCMC) searches in two independent and two coupled chains. Each chain had one million generations sampled every 50 generations with a 25% burn-in. We then generated the best consensus trees after evaluating the results by their effective sample sizes (ESS). The posterior probabilities of the MCMC searches were used as nodal supports.

For MSC, we first generated a gene tree per partition using IQ-TREE v2.3.2 with 1,000 UFBoot replicates. We then imported all gene trees into Accurate Species Tree Algorithm (ASTRAL-IV; Zhang et al. 2018; Zhang and Mirarab 2022) with default settings. Each analysis had 16 rounds of search and 16 rounds of subsampling and the posterior probabilities were used as nodal supports.

### Divergence time estimation

Due to the lack of synapomorphies, the classification of all fossils previously assigned to Pisauridae has recently been questioned (Wunderlich 2008; Magalhaes et al. 2020), so these fossils could not be used for time calibration. We calibrated the phylogeny using three relatively reliable fossils with a relevant family affiliation, following Magalhaes et al. (2020), as follows. First, we treated an unidentified juvenile lycosid (Penney 2001) from Dominican amber (15–20 million years ago (Ma); Iturralde-Vinent and MacPhee 1996) as a crown lycosid. Secondly, we treated *Oxyopes succini* Petrunkevitch, 1958 from Baltic amber (43– 48.5 Ma; Ritzkowski 1997) as a stem oxyopid. Finally, we treated *Syphax* cf. *megacephalus* L. Koch and Berendt, 1854 (see Wunderlich 2004) from Baltic amber as a stem thomisid (see also Piacentini and Ramírez 2019; Magalhaes et al. 2020).

*A posteriori* calibration was performed in RelTime (Tamura et al. 2012; 2018) using a relaxed local clock model under MEGA 11 (Tamura et al. 2021) with *Viridasius* sp. set as the outgroup to all Lycosoidea. We used the unpartitioned sparse UCE matrix and its best ML topology, generated from IQ-TREE v2.3.2, for time calibration analyses. The probability distributions of time at the calibration points were set as log-normal with their 95% confidence intervals specified close to their proposed fossil ages. This reconstructed ultrametric tree was then used in subsequent evolutionary analyses.

To test whether RelTime yielded reliable absolute divergence times (see Lozano-Fernandez et al. 2017; Beavan et al. 2020), we also ran *a priori* time calibration under MCMCtree (Yang 2007) on a matrix of 300 loci with 99% taxon completeness, randomly selected using Phyluce v1.7.3, in PAML 4.10 (Yang 2007). We selected the ML topology generated from the dense matrix as the starting tree after removing the branch lengths. Calibration points were selected and set as in the RelTime analysis using MCMCtreeR (Puttick 2019) in R 4.3.3 (R Core Team 2024). Following the program tutorial (dos Reis et al. 2017), we first estimated the average substitution rate per time unit as well as the length, gradient, and Hessian of each branch. With the above estimated parameters, we ran divergence time dating in two independent runs. Each run had 500,000 generations sampled every 50 generations and with a 10% burn-in. We compared the results of the two runs in MCMCtreeR, using the script from Gutiérrez-Trejo et al. (2024).

### Evolutionary analyses

We performed all evolutionary analyses in R 4.3.3 using the packages “ape” (Paradis et al. 2004), “phytools” (Revell 2024), and “MCMCglmm” (Hadfield 2010). We first pruned the distantly related outgroups (Zoropsidae, Psechridae, Oxyopidae, Thomisidae, Viridasiidae) from the ultrametric tree (using the command code “drop.tip” and “extract.clade” in “ape”) as they were irrelevant for the evolutionary questions posed, but retained Lycosidae and Trechaleidae because they nested with our ingroups (see Results). We also removed the pisaurid *Tallonia picta* Simon, 1889 due to insufficient trait data. Given the strong emphasis on *Dolomedes* species in our study, our taxon sample, if left unaltered, could lead to biased evolutionary outcomes (Ackerly 2000; Li et al. 2008; Sidlauskas 2008; Jorgensen et al. 2023). For all evolutionary analyses, we therefore used both the full taxon tree (putatively biased), as well as the average of randomized pruned taxon trees (putatively less biased), as follows. First, we assigned *Dolomedes* terminals to their biogeographic realms modified from Udvardy (1975), then randomly selected one terminal per realm to generate a reduced subset of *Dolomedes* terminals and pruned the other *Dolomedes* terminals from the tree. We repeated this randomization 30 times to generate 30 different reduced taxon trees, each presumably with a relatively balanced taxon sampling.

To reconstruct the evolution of semi-aquatic vs. terrestrial lifestyles and the presence/absence of capture webs, we first sought the best evolutionary model by fitting, on i) the full taxon tree and ii) the reduced taxon trees, four different assumptions of trait transition rate matrices: equal rates (ER), all-rates-different (ARD), state A to B only and B to A only, into *k*-state Markov models (M*k* model; Pagel 1994; Lewis 2001). For the evolutionary models estimated from the full taxon tree, we selected the one with the lowest AIC as the best model (Fig. 2a). For the evolutionary models estimated from the reduced taxon trees, we first averaged the AIC and the estimated trait transition probability generated from each model with the same assumptions on all reduced taxon trees (Fig. 2b). Then, we selected the evolutionary model with the lowest average AIC as the most suitable across all reduced taxon trees. To reconstruct the ancestral states of lifestyles and capture webs in *Dolomedes* and pisaurids, we simulated stochastic character maps 100 times on the full taxon tree, once using the best model from the full taxon tree, and again using the best-averaged model generated from the reduced taxon trees (Fig. 2b).

**FIGURE 2.**
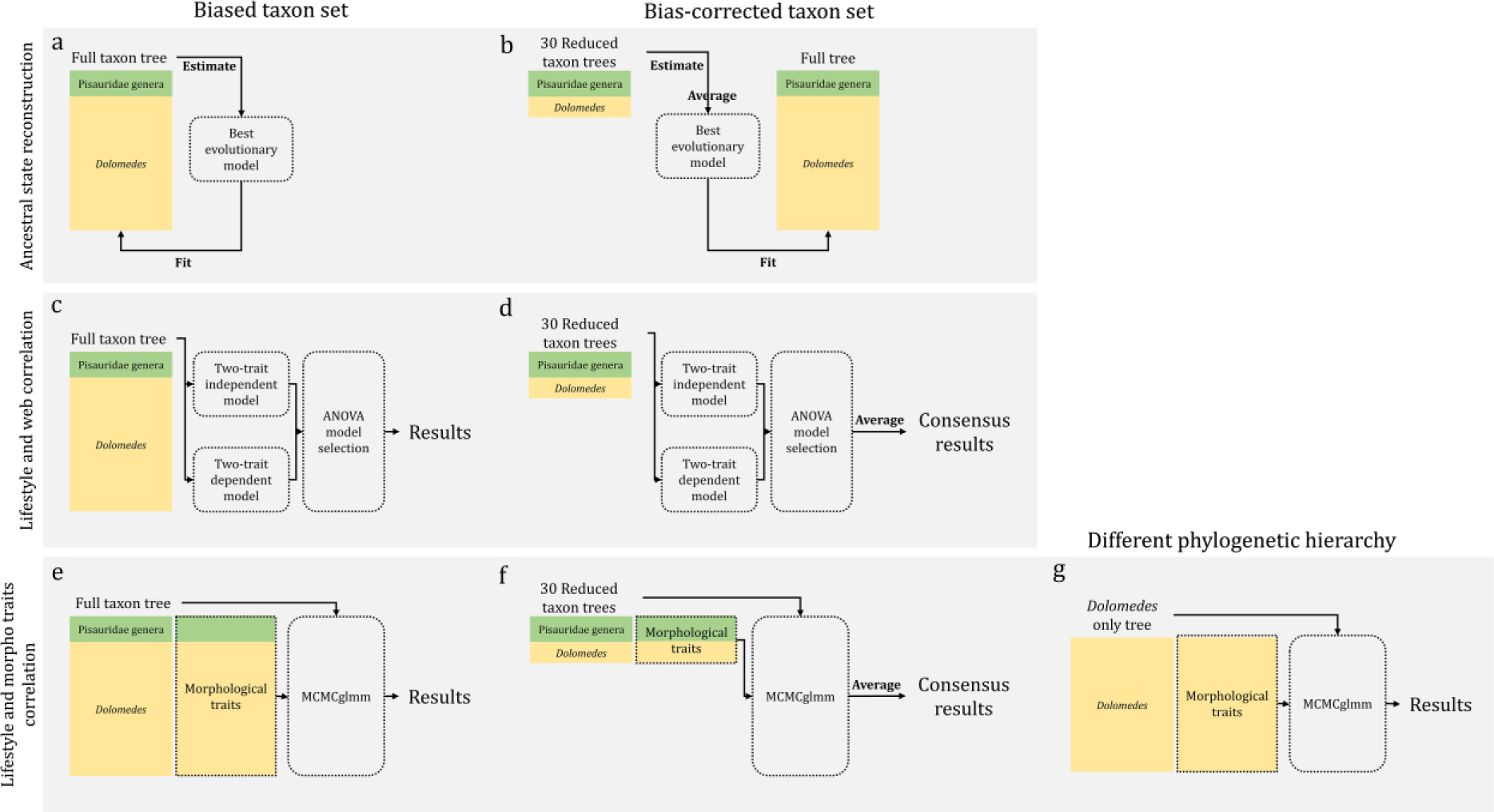
Analytical design for investigating how biased taxon sampling can influence the three evolutionary analyses; and how different phylogenetic hierarchies change the correlation between lifestyles and morphological traits. a–b, analytical design for ancestral state reconstruction using (a) the full taxon tree, and (b) 30 reduced taxon trees; c–d, analytical design for comparative analyses between lifestyles and capture webs using (c) the full taxon tree, and (d) 30 reduced taxon trees; e–g, analytical design for comparative analyses between lifestyles and morphological traits using (e) the full taxon tree, (f) 30 reduced taxon trees, and (g) the *Dolomedes*-only tree.

We used Pagel’s (1994) binary character correlation test to examine the evolutionary correlation between lifestyles and capture web on i) the full taxon tree and ii) reduced taxon trees. We first fit the two traits into a two-trait independent, and a dependent ARD-M*k* model. For the models estimated from the full taxon tree, we applied model selection by analysis of variance (ANOVA) to test whether the dependent model significantly better fit our data (Fig. 2c). For the models estimated from the reduced taxon trees, we further averaged the values in the statistic tables generated from the ANOVA model selections and the estimated trait transition probabilities of all the reduced taxon trees. We then used the averaged p-value and the number of reduced taxon trees that showed significance in each ANOVA model selection to seek the consensus on whether semi-aquatic versus terrestrial lifestyles and the presence or absence of capture webs were correlated across all reduced taxon trees (Fig. 2d).

We tested the correlation between lifestyles and three morphological traits (CW; L1/CW; T1/L1) using Markov chain Monte Carlo Sampler for Multivariate Generalized Linear Mixed Models (MCMCglmm; Hadfield 2010) on i) the full taxon tree (Fig. 2e) and ii) the reduced taxon trees (Fig. 2f) with 10 million MCMC iterations. The thinning interval and burn-in were set to 5,000 and 2.5 million, respectively. For those morphological trait(s) that were significantly (pMCMC < 0.05) correlated with lifestyles on the full taxon tree, we further examined their relationship with lifestyles by generating density curves of estimated moment-matching values (Hadfield 2010). To test for correlations between lifestyles and the three morphological traits on the reduced taxon trees, we averaged all values in the MCMCglmm tables from all the reduced taxon trees, and also counted the number of the reduced taxon trees that showed significance (Fig. 2f). Then, as above, we further examined the relationship between morphological traits with lifestyles by generating density curves of estimated moment-matching values (Hadfield 2010). As phylogenetic correlations among traits can be detected at some, but not all, hierarchical levels of a given phylogeny (see Wolff et al. 2022), we performed an additional set of MCMCglmm analyses on a full set of *Dolomedes* terminals only following the above algorithm for the full taxon tree (Fig. 2g).

## Results

The RTA spider probe set, on average, captured 2,790 UCE loci consisting of 2,815,984 base pairs (Table S1) with *Peucetia* sp. (Oxyopidae) yielding only 1,125 loci. After trimming and aligning, the sparse matrix with 75% taxon completeness contained 2,676 UCE loci with 1,221,957 base pairs, and the dense matrix with 95% taxon completeness contained 1,175 UCE loci with 875,429 base pairs. Only 94 loci were shared by all taxa.

Phylogenies reconstructed from the sparse and the dense matrix show identical topologies in all analyses except within *Dolomedes* (Fig. S1), where MSC recovered slightly different topologies (Figs. S1e–f), interpreted in the following paragraphs. With very few exceptions, all nodes receive full or high statistical support from UFboot, SH-aLRT, and BI posterior probabilities. Despite yielding a few additional poorly supported nodes within *Dolomedes*, most nodes are still strongly supported by MSC. Given the best overall supports, we choose the ML topology reconstructed from the sparse matrix as the preferred tree (Figs. 3 and 4; see also Supplementary Material).

**FIGURE 3.**
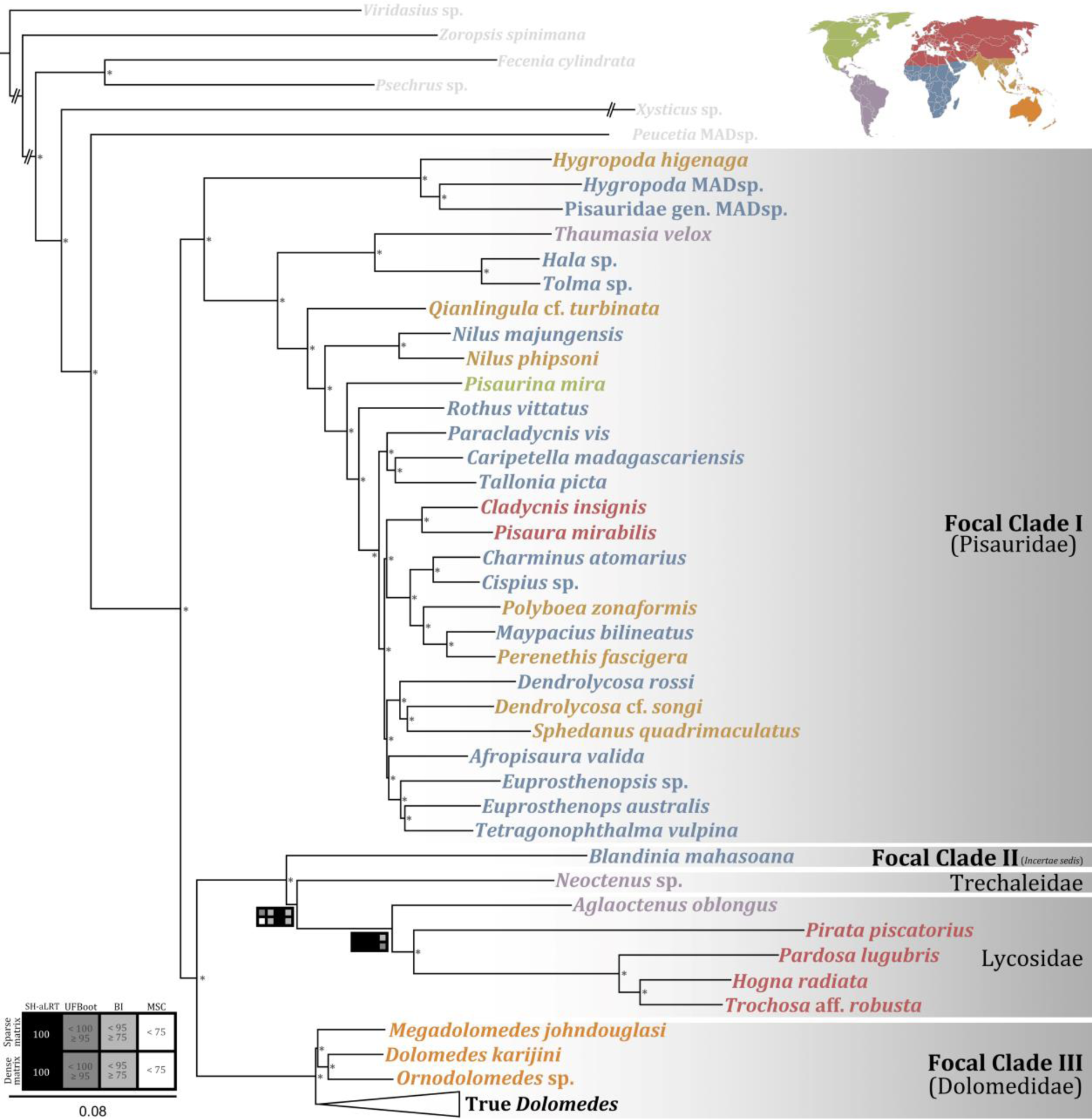
Maximum likelihood phylogeny of the three focal clades reconstructed from the sparse UCE matrix; nodal supports are labeled according to the legend at the bottom left, nodes with full supports are represented by “*”; each terminal is color-coded by its biogeographic region according to the map (modified from Udvardy 1975) at the top right with the outgroups in light gray. UFboot: ultrafast bootstrap (Minh et al. 2013; Hoang et al. 2017), SH-aLRT: SH-like approximate likelihood ratio test (Guindon et al. 2010), BI: posterior probability (%) of Bayesian inference, MSC: posterior probability (%) of multi-species coalescence. See supplementary materials for the full branch lengths of the outgroups.

**FIGURE 4.**
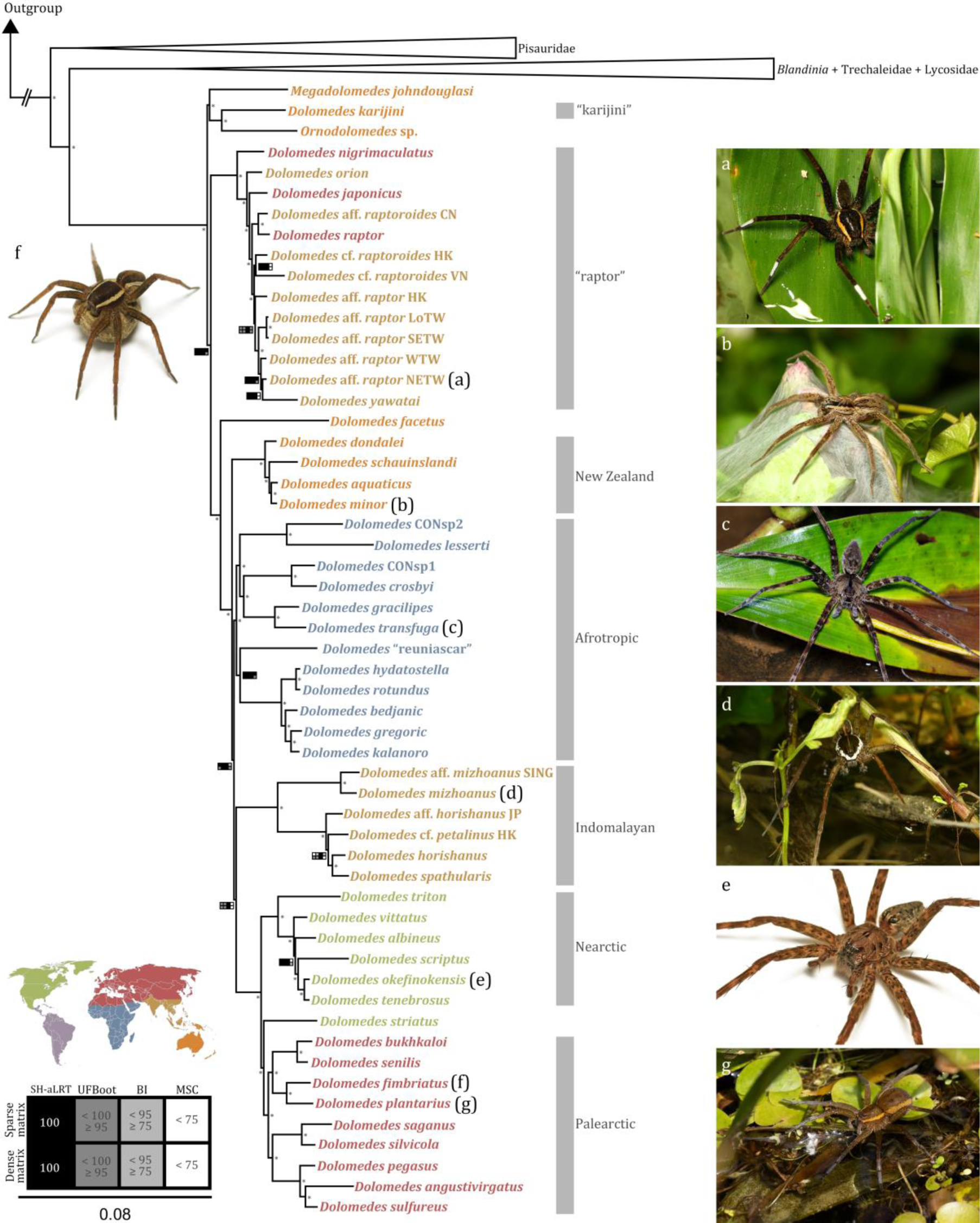
Maximum likelihood phylogeny of *Dolomedes* Latreille, 1804 and its relatives reconstructed from the sparse UCE matrix; nodal supports are labeled according to the legend at the bottom left, nodes with full supports are represented by “*”; each terminal is color-coded by its biogeographic region according to the map at the bottom left; images in panels a–g show habitus morphologies of the species labeled correspondingly in the phylogeny: (a) an unknown morphospecies of the “raptor” clade hiding between leaves, (b) a female *D. minor* L. Koch, 1876 guarding her nursery web, (c) a female *D. transfuga* Pocock, 1900 in hunting posture, (d) a male *D. mizhoanus* Kishida, 1936 in hunting posture, (e) a female *D. okefinokensis* Bishop, 1924 placed on a white background, (f) a *D. fimbriatus* (Clerck, 1757) carrying her egg sac, and (g) a female *D. plantarius* (Clerck, 1757) standing on macrophytes. UFboot: ultrafast bootstrap (Minh et al. 2013; Hoang et al. 2017), SH-aLRT: SH-like approximate likelihood ratio test (Guindon et al. 2010), BI: posterior probability (%) of Bayesian inference, MSC: posterior probability (%) of multi-species coalescence.

### Systematics of raft- and nursery web- spiders

All analyses reject the monophyly of Pisauridae as currently classified. This family-level name refers to a paraphyletic assemblage of genera that fall into three clades, labeled as focal clades in this paper (Fig. 3; see Taxonomy): 1) **Focal Clade I** groups the majority of the genera classified in Pisauridae, and because it contains *Pisaura* Simon, 1886, this clade represents the “true pisaurids”. Focal Clade I (Pisauridae) is sister to a taxon-rich clade containing all the taxa listed below; 2) **Focal Clade II** contains one genus that is currently still cataloged as a pisaurid, i.e. the monotypic *Blandinia* Tonini et al., 2016 from Madagascar (Fig. 1K). Because it is recovered as sister to Trechaleidae and Lycosidae, *Blandinia* cannot be considered a pisaurid, and is instead *incertae sedis*, pending better taxonomic resolution; 3) **Focal Clade III** contains *Dolomedes*, *Megadolomedes* Davies and Raven, 1980, and *Ornodolomedes* Raven and Hebron, 2018. This clade is sister to Focal Clade II, Trechaleidae, and Lycosidae (Fig. 3). Considering the phylogenetic results, the topological stability, the strong clade supports, and our understanding of the morphology of *Dolomedes* and its relatives, detailed in the Taxonomy section below, we will from here on formally refer to the Focal Clade III as Dolomedidae Simon, 1876, colloquially named *raft spiders*. We suggest the use of *fishing spiders* for *Dolomedes*.

Within Dolomedidae, we tested for the monophyly of *Dolomedes*. Strictly speaking, *Dolomedes* is not monophyletic. The majority of *Dolomedes* species, including the type species *D. fimbriatus* (Clerck, 1757), are in a clade that represents true *Dolomedes*, but one species, *D. karijini* Raven and Hebron, 2018, is recovered in the sister clade that also contains *Ornodolomedes* and *Megadolomedes* (Figs. 3 and 4). While the Afrotropical *Dolomedes* species are monophyletic within their biogeographic realm, other biogeographic realms do not show strict species monophyly (Fig. 4). For example, in addition to the distal Palearctic *Dolomedes* clade, other Palearctic species are found in the “raptor” clade, which is sister to all other true *Dolomedes* (Fig. 4). The Australasian species are not monophyletic, but the New Zealand species are (Fig. 4). An Indomalayan clade groups some, but not all, species from this region, as many Indomalayan terminals also form the “raptor” clade (Fig. 4). Most Nearctic species are monophyletic, but *D. striatus* Giebel, 1869 is sister to the Palearctic clade (Fig. 4).

Multi-species coalescence recovers three topological differences within *Dolomedes* (Figs. S1e–f) from the preferred tree (Fig. 4), two among the distal “raptor” clade and the other among the Indomalayan, Afrotropic, and Nearctic plus Palearctic (Holarctic) clades. The MSC phylogenies reconstructed from both matrices suggest the two morphospecies, *D*. cf. *raptoroides* HK and *D*. cf. *raptoroides* VN, are not sister as they are in the preferred tree (Fig. 4). Additionally, MSC has *D. yawatai* Ono, 2002 as sister to a clade with four *Dolomedes* morphospecies from Taiwan (Figs. S1e–f) instead of nesting within that clade (Fig. 4). MSC on the dense matrix suggests the Indomalayan clade as sister to a large clade including the Afrotropic and Holarctic *Dolomedes* (Figs. S1e–f). However, such topologies are poorly supported with their posterior probabilities (PP) between 0.6 and 0.7. In the preferred tree, Afrotropic *Dolomedes* are sister to a clade uniting the Indomalayan and the Holarctic clades, with better nodal supports (> 0.9; see Fig. 4).

### Divergence times of raft- and nursery web- spiders

RelTime and MCMCtree (results in parentheses) yield comparable divergence times of the three focal clades. Both analyses reconstruct relatively recent origins for the major clades in question. The crown that gave rise to all three focal clades plus Trechaleidae and Lycosidae is estimated to have originated between 32 and 37 Ma (34–43 Ma), in the late Eocene to early Oligocene (Fig. 5). The origin of Pisauridae (Focal Clade I), estimated between 29 and 35 Ma (31–40 Ma) (Fig. 5), is the earliest of the three focal clades. The Oligocene-Miocene split of *Blandinia* (Focal Clade II) from Trechaleidae plus Lycosidae is estimated between 21 and 31 Ma (24–34 Ma) (Fig. 5). Compared to the above two focal clades, the origin of Dolomedidae (Focal Clade III) is estimated to be more recent, between 10 and 15 Ma (12–17 Ma) in the mid-Miocene (Fig. 5). The origin of *Dolomedes* is estimated to be in the Miocene, between 9 to15 Ma (11–16 Ma) (Fig. 5), and since then, the clade has undergone a rapid diversification, with the 95% confidence intervals and highest posterior densities for final splits between species doublets lying between 0.2 Ma (0.3 Ma) (*D. hydatostella* Yu and Kuntner, 2024 and *D. rotundus* Yu and Kuntner, 2024 from Madagascar) and 9.2 Ma (8.9 Ma) (*D. lesserti* Roewer, 1955 from Benin and an unnamed *Dolomedes* species from D.R. Congo) (Fig. 5). On the other hand, some species that group with larger clades in our phylogeny could plausibly be much older, e.g. between 7.1 and 14.7 Ma (10.2–15 Ma) (*D. facetus* Koch, 1876 from Australia).

**FIGURE 5.**
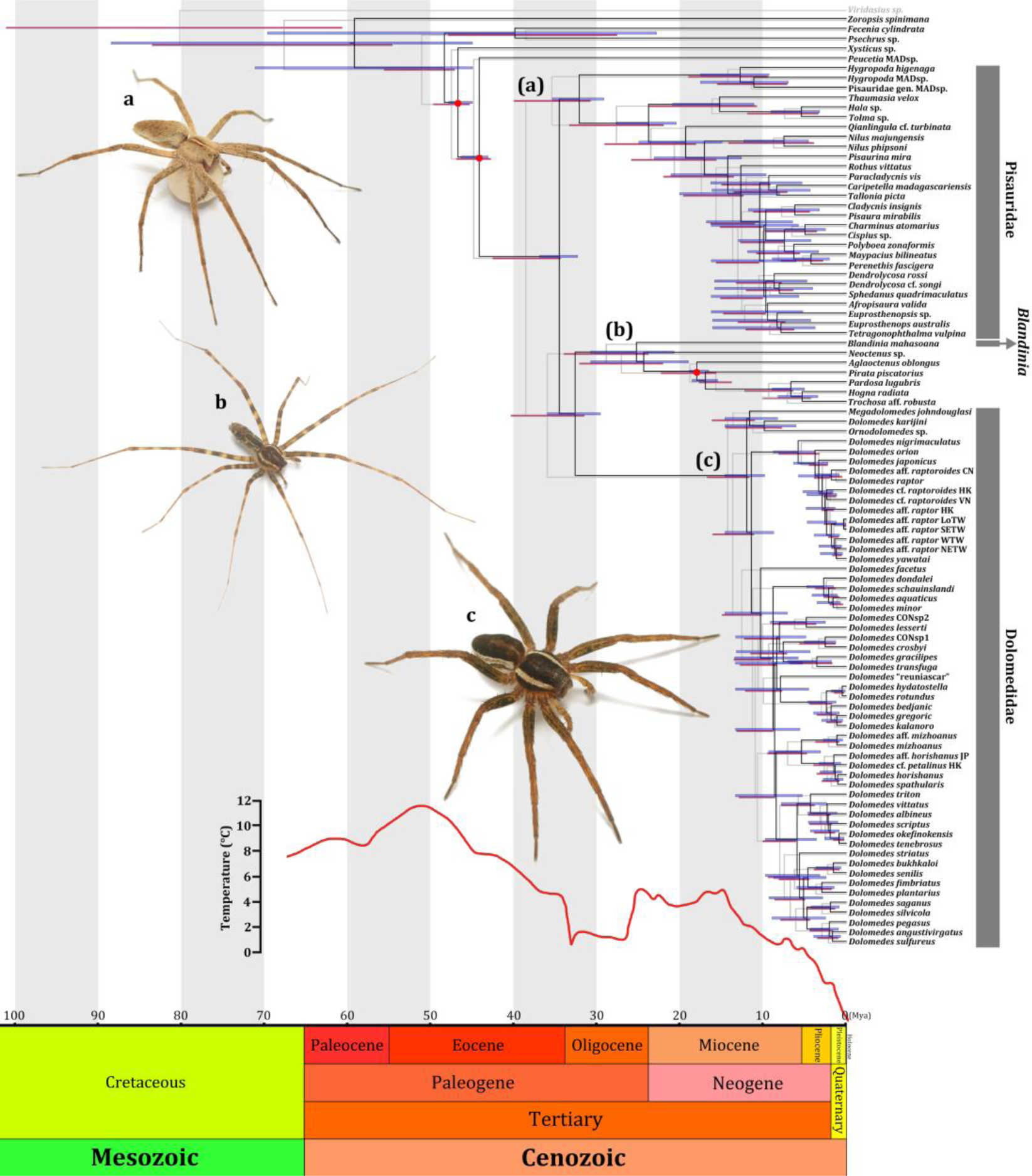
Phylogenies of all three focal clades (a–c) calibrated by RelTime (black lines and blue bars; Tamura et al. 2012; 2018) and MCMCtree (gray lines and red bars; Yang 2007) with 95% confidence intervals and highest posterior densities of diversification time; red dots represent the fossil calibration points. A red curve of paleotemperature, modified from Zachos et al. (2001), shows the overlap between diversification and paleoclimate changes. Ma: million years ago.

### Evolution of lifestyles in raft- and nursery web- spiders

With the lowest average AIC, the model allowing only transitions from a semi-aquatic to a terrestrial lifestyle (“semi-aquatic to terrestrial” model) is the best evolutionary model (Table 4) on the reduced taxon trees (Fig. S2). Partially rejecting our hypothesis, the semi-aquatic lifestyle is the ancestral state not only of Dolomedidae (Focal Clade III), but also of the entire clade that unites Pisauridae (Focal Clade I), Focal Clade II, Lycosidae, and Trechaleidae (Fig. 6A, see also Fig. S3). On the full taxon tree, the model detects no fewer than 15 reversals to a terrestrial lifestyle, including three in Pisauridae and eight in Dolomedidae. The terrestrial lifestyle in the focal clades of our tree, thus evolved from semi-aquatic ancestors. In Focal Clade I, all semi-aquatic terminals or genera, *Hygropoda*, *Thaumasia* Perty, 1833, *Qianlingula* Zhang et al., 2004, and *Nilus*, have retained the ancestral, semi-aquatic lifestyle (Fig. 6a). The seven terrestrial *Dolomedes* species are not closely related to each other (Fig. 6a), suggesting independent terrestrialization in each species. This result is not strongly affected by taxon sampling bias. Model selection on the full taxon tree namely also suggests the “semi-aquatic to terrestrial” model as the best (Table 4), and thus the reconstructed ancestral lifestyle is identical to that from the reduced taxon trees, albeit with different rates of trait transition (Fig. 6b, see also Fig. S4).

**FIGURE 6.**
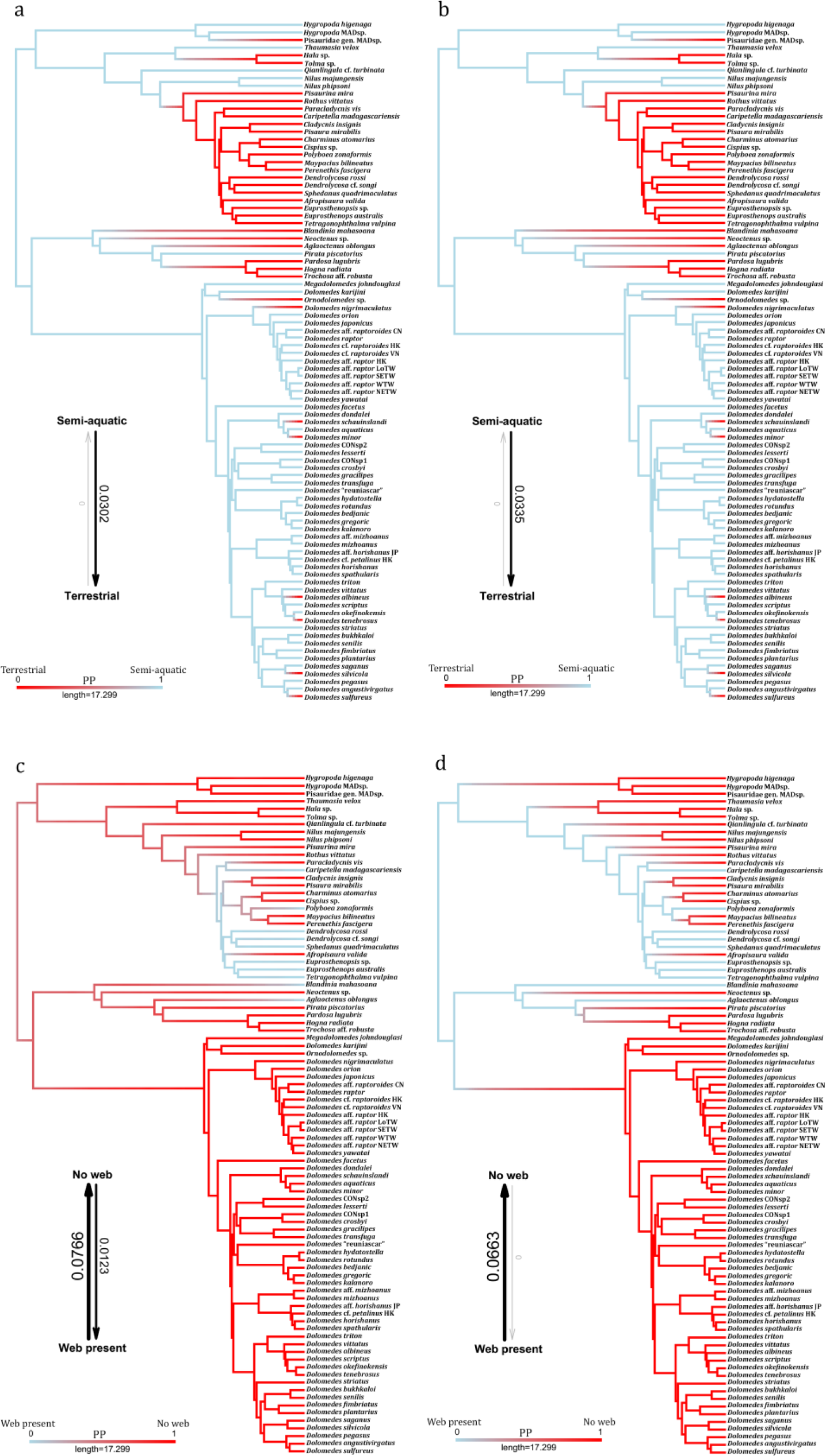
Ancestral state reconstructions rendered from 100 stochastic character mapping on the full taxon tree and trait transition probabilities of: a–b, semi-aquatic vs terrestrial lifestyle with their best *k*-state Markov models (M*k* model, Pagel 1994; Lewis 2001) estimated from the (a) 30 reduced taxon trees, and (b) full taxon tree; c–d, capture web absence vs presence with their best M*k* model estimated from the (c) 30 reduced taxon trees, and (d) full taxon tree. PP: posterior probability.

**TABLE 4.**
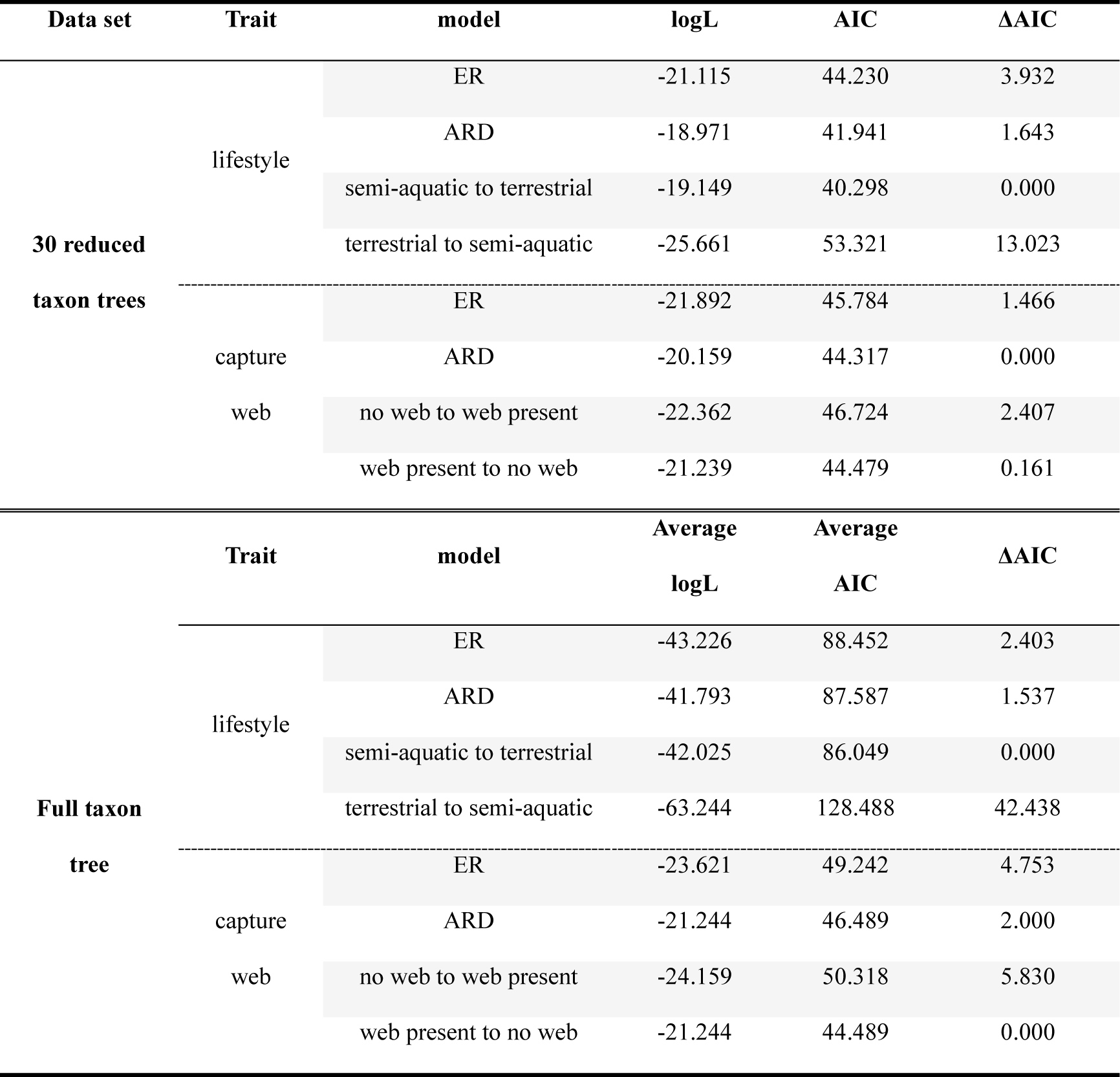
AIC comparisons among the *k*-state Markov models (Mk model, Pagel 1994; Lewis 2001) with different trait transition rates assumptions conducted on the 30 reduced taxon trees and the full taxon tree. ER: equal rates, ARD: all-rates-difference.

After averaging on the reduced taxon trees (Fig. S2), the ARD is the best model to explain the evolution of the capture web (Table 4); the reconstruction does not recover a clear ancestral state (Fig. 6c, see also Fig. S5). Due to this ambiguity, the hypothesis of an ancestral absence of a capture web for the clade containing Pisauridae, *Blandinia*, and Dolomedidae is not supported. Reconstructions on the full taxon tree suggest at least three independent origins of the capture web: one each in Lycosidae, *Blandinia*, and distally within the Focal Clade I, the latter containing numerous reversals to no web (Fig. 6c). Within *Dolomedes*, however, the absence of a capture web is ancestral, as partially predicted by the hypothesis (Fig. 6c). Furthermore, the absence of a web is ancestral for the entire Dolomedidae, with no reversals (Fig. 6c). This analysis is strongly affected by the taxon sampling bias and its correction. Model selection on the full taxon tree namely suggests that the model allowing only transitions from web presence to absence, rather than the ARD, is the best model. This model reconstructs the presence of a capture web as the ancestral state of all three focal clades, Trechaleidae, and Lycosidae with numerous reversals to no web (Fig. 6d, see also Fig. S7). Such results would outright reject our hypothesis of an ancestral absence of a web in Pisauridae.

Our analyses also reject the hypothesis of an evolutionary correlation between capture webs and lifestyles. Instead, the evolution of the web presence or absence and semi-aquatic versus terrestrial lifestyles are independent (Fig. 7a). None of the reduced taxon trees suggest that the two-trait dependent model fits our data significantly better than the independent model (average p-value of the ANOVA model selection = 0.1, see also Supplementary Material). However, this result is strongly affected by the likely bias in the taxon sample on the tree. On the full taxon tree, the two-trait dependent ARD model fits the data significantly better than the two-trait independent ARD model (see also Supplementary Material), suggesting that a semi-aquatic lifestyle excludes the presence of a capture web (Fig. 7b). This result would corroborate the predicted evolutionary correlation between these traits.

**FIGURE 7.**
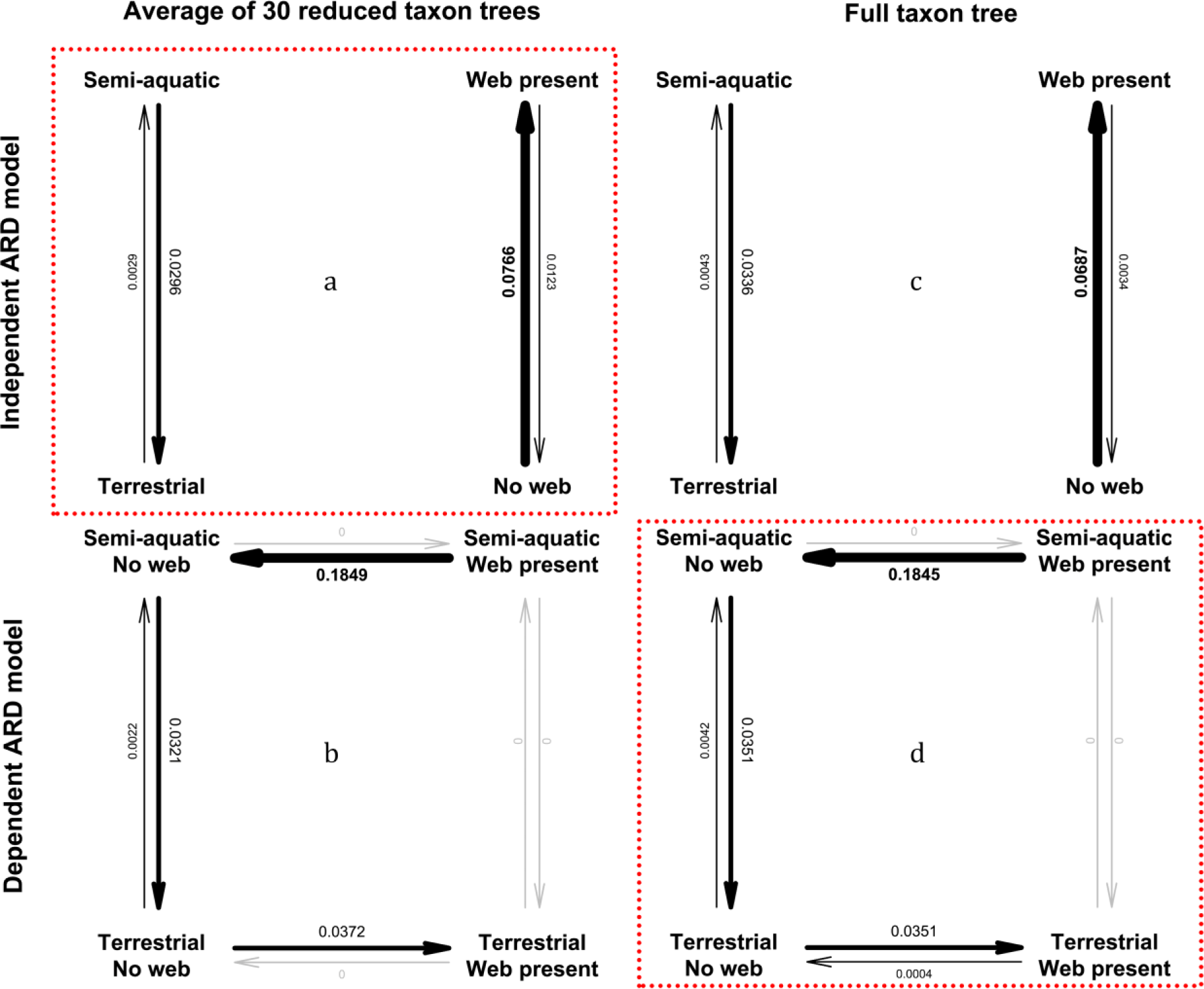
Pagel’s (1994) binary character correlation test between semi-aquatic vs terrestrial lifestyle and capture web absence vs presence based on the: a–b, 30 reduced taxon trees with their estimated and averaged trait transition probabilities in two-trait (a) independent all-rates-different (ARD) models, and (b) dependent ARD models; c–d, full taxon tree and their estimated trait transition probabilities in two-trait (c) independent ARD models, and (d) two-trait dependent ARD models. The model selected by model selection highlighted by red dotted line.

Our final set of hypotheses predicted that semi-aquatic taxa would exhibit wider carapaces, relatively shorter legs I, and relatively longer tarsi I compared to the terrestrial taxa in the phylogeny. The results support only the first hypothesis (Table 5). The significance of a wider carapace in semi-aquatic taxa is recovered in 29 out of 30 reduced taxon trees (see Supplementary Material). On average, the carapace of semi-aquatic spiders is 2.4 mm wider than that of the terrestrial spiders in the phylogeny (Fig. 8a). On the other hand, no significance is detected in the relative length of leg I and tarsus I between semi-aquatic and terrestrial taxa (Table 5). These results appear to be less susceptible to taxon bias, as analyses using the full taxon tree yield identical results (Table 5, Fig. 8b). However, when only *Dolomedes* terminals are considered, all three of the above hypotheses are rejected (Table 5).

**FIGURE 8.**
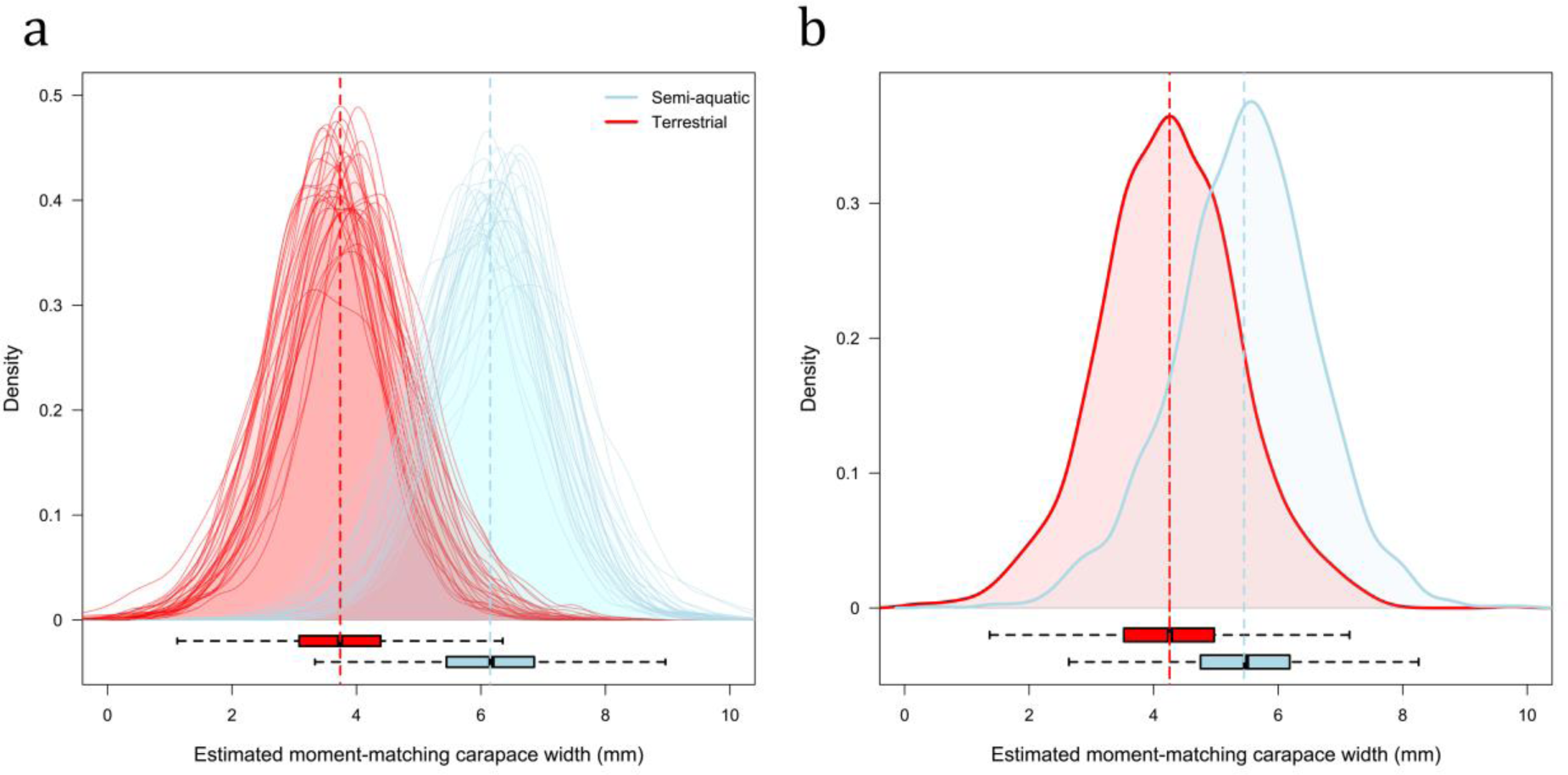
Posterior density curves of moment-matching carapace width estimated via Monte Carlo Sampler for Multivariate Generalized Linear Mixed Models (MCMCglmm; Hadfield 2010) from the (a) 30 reduced taxon trees and (b) full taxon tree; estimated mean carapace width of each lifestyle pinpointed by the vertical dotted lines. The box plots below show the medians (bold lines), first quartiles (Q1, left margins of the boxes), third quartiles (Q3, right margins of the boxes), Q1 − 1.5 × interquartile range (IQR) (left whisker), and Q3 + 1.5 × IQR (right whisker) of all estimated values in both lifestyles.

**TABLE 5.**
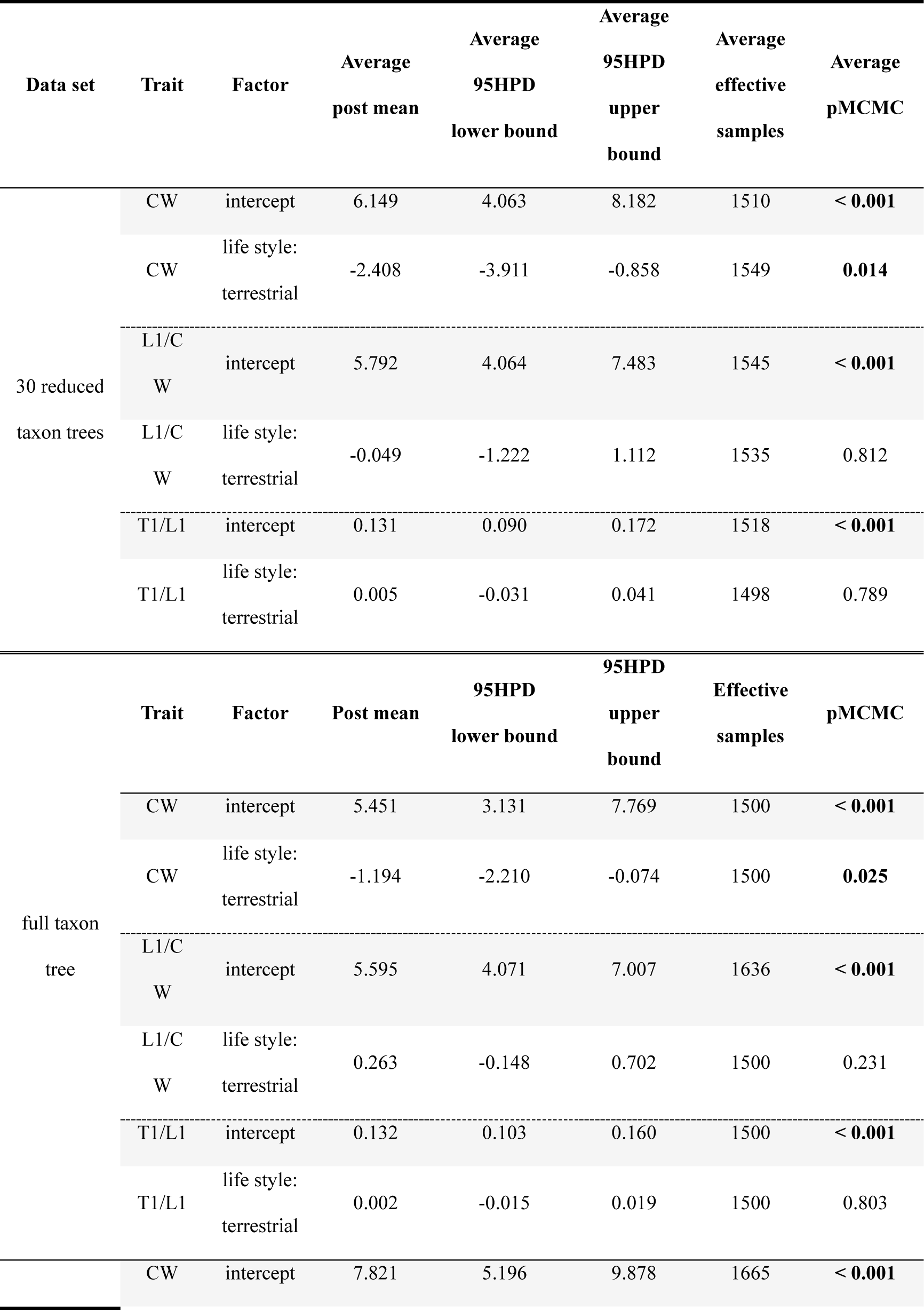
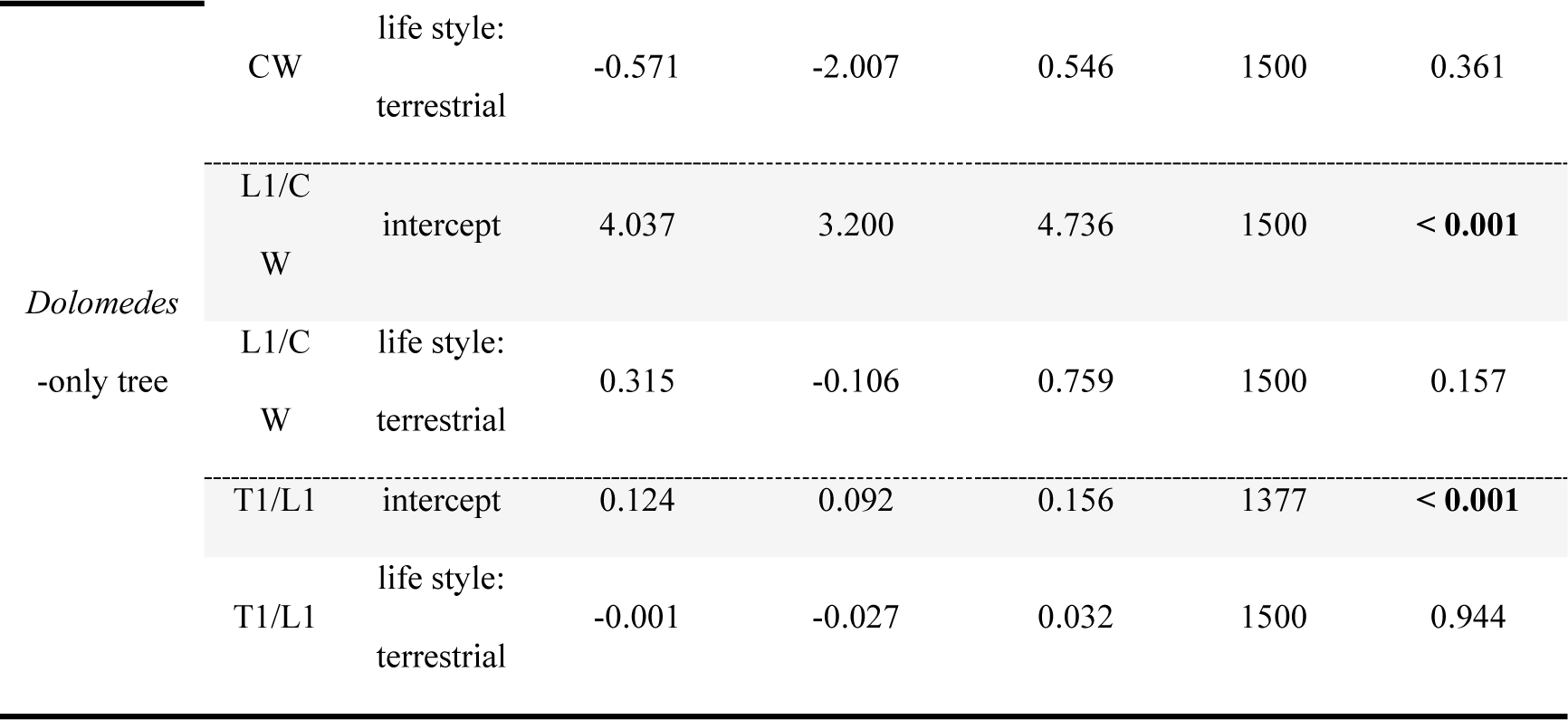
Statistic tables of Monte Carlo Sampler for Multivariate Generalized Linear Mixed Models (MCMCglmm; Hadfield 2010) between lifestyles and carapace width, relative length of teg I, and relative length of tarsus I, showing the correlation between morphological traits and semi-aquatic versus terrestrial lifestyles. CW: carapace width, L1: leg I length, T1: tarsus I length, 95HPD: 95% highest posterior density, pMCMC, twice the probabilities of the posterior values that are above or below zero.

## Discussion

The lack of comprehensive phylogenies has so far precluded a thorough systematic account of *Dolomedes* and Pisauridae *sensu lato*, as well as generalizations of the lifestyle evolution of this global assemblage of spider genera. We pursue these broad goals in the phylogenomic and comparative study reported here. Our phylogenetic results reject the monophyly of both Pisauridae and *Dolomedes*. The genera previously placed within Pisauridae fall into three focal clades: the true Pisauridae, *Blandinia*, and Dolomedidae, whose family rank we formally resurrect (see Taxonomy). The origin of each of the focal clades is estimated to be relatively recent, i.e. 40 Ma or less. *Dolomedes* is paraphyletic, as *D. karijini* groups with *Ornodolomedes* and *Megadolomedes*. Our reconstructions partially reject our first hypothesis that the semi-aquatic lifestyle is ancestral to *Dolomedes* but not to Pisauridae. A semi-aquatic lifestyle is reconstructed as ancestral for all three focal clades, followed by independent reversals to terrestrial lifestyle. Our second hypothesis of the ancestral absence of a capture web in *Dolomedes* and Pisauridae is also partially rejected. The evolution of a capture web is ancestrally ambiguous, with a web origin in both Pisauridae and *Blandinia*. In Dolomedidae, on the other hand, the unequivocal ancestral absence of a capture web is retained through their evolutionary history. Our third hypothesis, which predicted that a semi-aquatic lifestyle would exclude the presence of a capture web, is also rejected, because our evolutionary tests reveal an independent evolution of lifestyle and capture web. Of the hypotheses that semi-aquatic pisaurids having larger body sizes, relatively shorter legs, and relatively longer tarsi, our comparative analyses support only the larger body size hypothesis, but only at higher hierarchical levels and not within *Dolomedes* species. Contrary to our prediction, the relative lengths of legs and tarsi do not differ between lifestyles. Our study is the first step towards stability in systematics of raft and nursery web spiders and at the same time provides a phylogenetic basis for further comparative research.

### Dolomedidae: A step towards the redefinition of Pisauridae

We provide a well-sampled and robust phylogenomic backbone to support the resurrection of Dolomedidae, Lehtinen’s (1967) taxonomic hypothesis that has regained recent attention (Albo et al. 2017; Hazzi and Hormiga 2023; Kulkarni et al. 2023). Our phylogeny is consistent with the topologies found by Hazzi and Hormiga (2023) and Kulkarni et al. (2023) that *Dolomedes* is sister to the clade including Lycosidae, Trechaleidae, and additionally *Blandinia*. Although several putatively closely related genera of *Dolomedes*, including *Bradystichus* Simon, 1884, *Caledomedes* Raven and Hebron, 2018, *Mangromedes* Raven and Hebron, 2018, and *Tasmomedes* Raven and Hebron, 2018, were not available for inclusion with UCE data, additional evidence from morphological (Platnick and Forster 1993; Raven and Hebron 2018) and molecular (Piacentini and Ramírez 2019; Kulkarni et al. 2023) studies both suggest that these genera fit our definition of Dolomedidae rather than Pisauridae. This updated classification of Dolomedidae is formalized in the Taxonomy.

The *Dolomedes* paraphyly hinges on one species, *D. karijini*, and could be resolved through an establishment of a new genus for this species group. Although we envision this solution being valid in the future, we refrain from doing so here due to our limited access to Australian species. Based on details of the genitalia, Raven and Hebron (2018) treated *D. karijini* in the *D*. *albicomus* group with *Dolomedes* species from Australia and New Caledonia. Considering the incomplete taxon sampling and the current ambiguity of species boundaries within *Dolomedes* (Tanikawa and Miyashita 2008; Vink and Dupérré 2010; Yu and Kuntner 2024), it is unclear whether the entire species group, or only *D. karijini* should be elevated to a new genus. This decision will await future integrative taxonomic work with more Australasian species.

Our taxonomy is only the first step towards a better definition of Pisauridae and its subfamilies. Although our phylogenies reject the classification of *Blandinia* as a pisaurid, it is unclear whether this monotypic genus can be assigned to another existing family. Its placement in Trechaleidae would be unfounded, given conflicting phylogenies (see Piacentini and Ramírez 2019; Hazzi and Hormiga 2023; Kulkarni et al. 2023). In the Taxonomy below, we therefore avoid creating a new family and hence treat *Blandinia* as *incertae sedis* until further evidence is available. Our phylogeny also partially rejects the currently used pisaurid subfamilial classifications, Thaumasinae, Thalassinae, and Pisaurinae (Simon 1890; Sierwald 1990), a problem that awaits more focused systematics efforts.

### Independent evolution of lifestyles and capture web

Most spiders are terrestrial, so the evolution of aquatic or semi-aquatic lifestyles in this group is an evolutionary oddity worthy of in-depth research. Species and genera within at least 22 spider families are known to have aquatic or semi-aquatic lifestyles (Crews et al. 2020). A comparative study across 42 spider families by Stratton et al. (2004) recovered aquatic adaptive locomotion and exoskeleton morphologies as ancestral to Pisauridae, Lycosidae, and Trechaleidae. These results agree with ours that a semi-aquatic lifestyle is ancestral to those three families as well as to Dolomedidae. However, research focusing on Dictynidae O. Pickard-Cambridge 1871 (Crews et al. 2020) and Lycosoidea (Piacentini and Ramírez 2019), which includes Pisauridae and *Dolomedes*, both recovered a terrestrial lifestyle as ancestral to these clades, followed by multiple origins of aquatic or semi-aquatic lifestyles. Our results, which suggest that a semi-aquatic lifestyle is ancestral to both Pisauridae and Dolomedidae, do not align with those above (Piacentini and Ramírez 2019). This discrepancy may stem from the less taxon sampling focus on terrestrial lycosids and a more complete taxon sample of pisaurids and dolomedids in this present study compared to that of Piacentini and Ramírez (2019). Character states of long and basal branches are more likely to be reconstructed as ancestral (see Li et al. 2008), and our study indeed detected several newly sequenced semi-aquatic pisaurids at the base of the family, and, additionally, improved the phylogenetic resolution of *Dolomedes*.

Aside from the widely documented freshwater bound, semi-aquatic lifestyle, Dolomedidae also comprises the marine intertidal *Mangromedes* (Raven and Hebron 2018) and the mud-dwelling *Bradystichus* (Platnick and Forster 1993; Raven and Hebron 2018; Harvey M.S., personal observation). The marine intertidal lifestyle is very rare in the spider tree of life (Raven 1988; Crews et al. 2020; Yu et al. 2023). Mud-dwelling, likewise, is rare and may represent an intermediate state between semi-aquatic and terrestrial lifestyles. These extraordinary cases could imply a more complex and diverse evolutionary history of water-associated lifestyles in Dolomedidae compared to Pisauridae. However, detailed evolutionary analyses of these additional lifestyles will require a phylogeny including all dolomedid genera.

The evolution of capture webs in spiders has been the focus of numerous recent studies (Blackledge et al. 2009; Bond et al. 2014; Garrison et al. 2016; Cheng and Piel 2018; Fernández et al. 2018; Kallal et al. 2021). By focusing on all spiders, these studies were unable to explore the lifestyle diversity within large and speciose clades, such as Pisauridae. The above studies have not reached a consensus on the ancestral presence or absence of a capture web in Pisauridae, Lycosoidea, or the RTA clade by either reconstructing no web as ancestral (Garrison et al. 2016; Fernández et al. 2018; Piacentini and Ramírez 2019), the presence of a ground sheet web as ancestral (Blackledge et al. 2009), or producing an ambiguous/unresolved result (Bond et al. 2014; Cheng and Piel 2018; Kallal et al. 2021). These differing results are not surprising, considering that only three of the above studies (Cheng and Piel 2018; Piacentini and Ramírez 2019; Kallal et al. 2021) included any web-building pisaurids, in addition to non-web-building representatives. Our study covers most of the pisaurids known to build capture webs, but does not unambiguously resolve the ancestral state. This strongly suggests that the results of the above works, which were based on very few selected pisaurid taxa were over-resolved. Among the known web-building pisaurids unavailable in our study is the South American *Architis* Simon, 1898, which, based on published morphological (Santos 2007a) and phylogenetic studies (Wheeler et al. 2017; Piacentini and Ramírez 2019; Hazzi and Hormiga 2023), might belong to the clade containing *Thaumasia*, *Hala* Jocqué, 1994, and *Tolma* Jocqué, 1994—all webless genera. If this is the case, we expect this clade to represent another case of independent origin of a capture web in pisaurids.

As no pisaurid or dolomedid taxa are known to be both semi-aquatic *and* build a capture web (Table 1), we hypothesized that these traits are evolutionarily exclusive, at least in wandering spiders. Although some studies may suggest that in Araneoidea, these two traits may co-occur in Tetragnathidae Menge, 1866, Linyphiidae Blackwall, 1859, Theridiidae Sundevall, 1833, and Theridiosomatidae Simon, 1881 (see Crews et al. 2020), the most convincing cases are rare (Eberhard 2020) and seemingly aberrant. One such case is *Conculus* Kishida, 1940 (Anapidae Simon, 1895) that is known for its diving behaviors and for building a part of its capture web underwater (Komatsu 1940). The other case is *Wendilgarda* Keyserling, 1886 (Theridiosomatidae) that attaches its web anchors to the water surface (Coddington and Valerio 1980). Curiously, this is the first study to test for an evolutionary correlation between these two traits. Even more curious is that, contrary to our expectations, the result suggests their independence. The above cases of semi-aquatic *and* web-building araneoids may not be so aberrant after all and the evolution of semi-aquatic lifestyle and the presence of a capture web may not be mutually exclusive across Araneae.

Aside from capture and nursery webs, literature also documents other webbing types constructed by pisaurids and dolomedids, both terrestrial and semi-aquatic, throughout their ontogeny. Such examples are juvenile webbing in *Thaumasia* (Eberhard 2020), *Qianlingula* (Yu K.-P.; personal observations), and *Dolomedes fimbriatus* (Jäger P.; personal observations). Although these types of non-capture webs were not considered in our comparative analyses, which only considered adult phenotypes, the abilities of semi-aquatic spiders to build large-scale webbing may also agree with the independent evolution of lifestyles and capture webs. Future research might profitably look into similarities and differences in web structure, composition, and web building behaviors among these different types of webs at different ontogenetic stages.

### Morphological adaptations to a semi-aquatic lifestyle

Our comparative analysis at a higher phylogenetic hierarchy revealed that semi-aquatic representatives of all three focal clades have wider carapaces compared to terrestrial ones. This finding is consistent with the general understanding that semi-aquatic and aquatic animals are larger or heavier than terrestrial ones, whether in mammals (Gearty et al. 2018), reptiles (Meiri 2008), birds (Gaston and Blackburn 1973), or invertebrates (Sabo et al. 2002; Braddy et al. 2008). Biological explanations for this phenomenon usually invoke buoyancy and gravity (Reynolds 1977; Reynolds and Karlotski 1977), trophic structure (Shurin et al. 2006; Tucker and Rogers 2014), thermoregulation (Downhower and Blumer 1988; Meiri 2008), aquatic locomotion (Schmidt-Nielsen 1971; Anderson et al. 1979), energy trade-off (Gearty et al. 2018), and predator avoidance (Wolff and Guthrie 1985). However, these explanations are more relevant to vertebrates, whereas gravity and thermoregulation may be less applicable to much smaller invertebrates, such as spiders.

We hypothesize that larger sizes of semi-aquatic pisaurids and dolomedids may be related to a biomechanical threshold above which surface tension can be broken when spiders forage under water. For generalist predators, aquatic habitats offer additional food resources compared to the terrestrial habitats, e.g. crustaceans, fishes, and amphibians, but only if the predator can reach underwater and subdue this relatively large prey. Semi-aquatic spiders that are large enough to break through the surface tension to access these additional food sources, while avoiding being eaten, may therefore have a competitive advantage over terrestrial species. Our field observations on *Dolomedes plantarius* (Clerck, 1757) that smaller juveniles usually do not dive when disturbed but larger adults and juveniles usually do (Yu K.-P., Klemen Č, and Kuntner M., personal observation) may add credibility to this hypothesis. Additionally, larger size in animals can facilitate aquatic locomotion through increased area in touch with the water surface and consequentially increased buoyancy (Soncini and Klein 2023), and by holding larger and denser hydrophobic structures. However, semi-aquatic species do not simply evolve to become increasingly larger as their maximum size cannot break the physical equilibrium of rafting on water surfaces (Soncini and Klein 2023).

We repeated the test of correlation between body size and lifestyle on *Dolomedes* terminals only and found that semi-aquatic species of *Dolomedes* were not significantly larger than the exclusively terrestrial ones. It is not unusual for comparative work to find conflicting results at different phylogenetic hierarchies, including examples in a study about correlation between morphological traits and foraging guilds in spiders (Wolff et al. 2022). In many such cases, explanations are not readily available, but our case does not seem complicated. Why is that? The vast majority of known *Dolomedes* are semi-aquatic, and this lifestyle is ancestral and nearly universal in this clade, with only a few derived species having recently evolved terrestrial lifestyles. In these cases, the time on land, and presumably a relaxed selection for larger sizes, has simply not been long enough for these species to evolve smaller sizes that are otherwise more typical of terrestrial inhabitants. This analysis of *Dolomedes* terminals only may therefore be a less powerful test of terrestrial traits compared than an analysis of all focal clades at the level of genera.

### Reducing taxon sampling bias in trait evolution analyses

Our research provides a case study of how taxon sampling bias can affect the outcome of commonly used evolutionary analyses and how to avoid this bias. Trait transition probability matrices (Q), estimated by fitting the extended M*k* model to the phylogeny, are sensitive to the number of terminals of each character state (Lewis 2001; see also Jorgensen et al. 2023). Therefore, if the trees or clades do not contain all extant taxa, a biased taxon coverage in each character state could result in misleading or less robust results (Ackerly 2000; Sidlauskas 2008; Jorgensen et al. 2023). Two solutions have been proposed to reduce such bias: i) calibration of Q by additional parameters derived from the taxon sampling completeness (Jorgensen et al. 2023); and ii) data selection (Ackerly 2000; Sidlauskas 2008). In our study, taxon sampling was heavily weighted towards *Dolomedes*, a genus, in which most species are semi-aquatic and none have a capture web. The relative over-sampling of a homogenous clade could lead to inflation or reduction of certain transition probabilities. Due to the unknown lifestyles in many pisaurid genera, calibration parameters by taxon sampling completeness were not feasible. We therefore approached the reduction of taxonomic bias in our analyses through data selection by randomly pruning *Dolomedes* terminals from the full taxon tree, corrected for their representation of biogeographic realms. We subjected all our evolutionary and comparative tests on trees with and without bias correction, i.e. the full taxon tree, and 30 reduced taxon trees.

Comparing the outcomes of the analyses with and without bias correction reveals that the ancestral state reconstruction of lifestyles and the correlation test between lifestyles and morphological traits are relatively robust to taxon sampling bias. However, the ancestral state reconstruction of a capture web and the correlation between lifestyle and capture web are sensitive to taxon sampling bias. Without reducing the taxon sampling bias, both analyses yielded over-resolved ancestral states, as well as misleading correlation outcomes between pairs of traits.

### RelTime and MCMCtree yield congruent results

We subjected the data to two approaches to divergence time estimation, the *a posteriori* and the *a priori* methods implemented in RelTime (Tamura et al. 2012; 2018) and MCMCtree (Yang 2007), respectively. Both types of analyses yielded largely overlapping results with interpretable confidence intervals and posterior densities, which may give considerable credibility to the estimated absolute and relative ages of the focal clades in our study. Credibility of the estimated absolute ages in our study is also suggested by comparing the hypothesized ages of the two island endemic species *Dolomedes schauinslandi* (0.9–4.2 Ma) and *D*. *orion* Tanikawa, 2003 (2.3–6.4 Ma) with the proposed time of the major geological events having shaped the Chatham Islands (2–3 & 4–6 Ma; Heenan et al. 2010) and Okinawa Islands (6–10 Ma; Wang et al. 2014).

The estimated origin of Pisauridae (29–40 Ma) is consistent with the literature (25–75 Ma; see Piacentini and Ramírez 2019; Magalhaes et al. 2020). However, our estimation yields a markedly younger origin of the crown Dolomedidae (compare 10–17 Ma in our study with 25–50 Ma in Piacentini and Ramírez 2019; Magalhaes et al. 2020). These differences could be due to variation in taxon sampling, amount of molecular data, and different topologies. Despite the relatively recent origin of the crown Dolomedidae, it sits on a long stem, as the split between Dolomedidae and its sister clade is estimated to be between 30 and 40 Ma. While the shape of this chronogram could plausibly be an artifact of missing some critical (extinct) taxa, we find the interpretation more likely that this shape reflects a recent diversification of Dolomedidae with relatively rapid species radiations, similar to the pattern in the extant segmented spiders (Xu et al. 2015). The incomplete reproductive isolation implied by the introgression between the sister species *Dolomedes minor* and *D. aquaticus* Goyen, 1888 (Lattimore et al. 2011) may reinforce the interpretation that *Dolomedes* undergoes rapid species radiations.

### Did Miocene climate change drive lifestyle diversification?

Cenozoic climate oscillations may have shaped the evolution of the terrestrial lifestyles and the presence of capture webs in Pisauridae. Combining the results of time calibration and ancestral state reconstructions, the three reversals from semi-aquatic to terrestrial lifestyles and the origin of capture webs in Pisauridae are all estimated between 10 and 20 Ma. This time interval matches the era when the Earth’s temperature and humidity reached the optima and started dropping (Zachos et al. 2001; Sun et al. 2020). Considering i) the extent to which semi-aquatic pisaurids are all distributed in the tropics, and ii) the distant phylogenetic proximity among Palearctic and Nearctic pisaurids, one can hypothesize that the cooling and drying climate in the mid-Miocene may have driven the evolution from semi-aquatic to terrestrial lifestyles (see also Ye et al. 2018), which may have spurred the emergence of web-based foraging behaviors.

The terrestrial lifestyle in *Dolomedes* may have undergone a different evolutionary scheme compared to pisaurids, judging from the fact that the reversals to terrestrial lifestyle in *Dolomedes* are phylogenetically derived and independent. Not unlike the semi-aquatic pisaurids, extant *Dolomedes* species of early branching lineages are distributed in subtropical to tropical Asia and Australasia, corresponding to regions that harbored humid refugia during the Miocene (Milne 2006; Steinthorsdottir et al. 2021). However, the more northern distribution ranges of some terminals from the same lineages strongly suggest that *Dolomedes* species can cope more easily with cold climates than the semi-aquatic pisaurids. During Miocene climate oscillations, *Dolomedes* may therefore have taken over niches and habitats that became less hospitable to other large semi-aquatic spiders. This could perhaps explain their colonization of most continents, and the current biogeographic patterns of *Dolomedes* as the only large semi-aquatic spiders inhabiting temperate or even boreal regions of the Northern Hemisphere. This scenario, on the one hand, and the observed cohabitation of certain terrestrial and semi-aquatic sister species of *Dolomedes*, –on the other (Ono 2009; Vink and Dupérré 2010), suggest that the evolution of the terrestrial lifestyle in *Dolomedes* stems from episodes of rapid radiation and niche partitioning.

## Taxonomy

### Taxonomic and phylogenetic history of Dolomedidae and Pisauridae

Among the first organisms to be named using Linnaeus’ binomial nomenclature, *Araneus fimbriatus* Clerck, 1757 and *Araneus plantarius* Clerck, 1757 from Sweden, represent the first descriptions of the species now known as *Dolomedes*. Latreille (1804) introduced the genus name *Dolomedes* and classified it under “wolf spiders” to include these two species, along with *Araneus mirabilis* Clerck, 1757 [today *Pisaura mirabilis* (Clerck, 1757), the type species of *Pisaura* and thus Pisauridae]. Simon (1864) placed *Dolomedes* in Lycosidae and later in the lycosid subfamily “Dolomedinae” (Simon 1876). This was the original designation of Dolomedinae as a family-level taxon, and thus we attribute Dolomedidae to this author, eventhough Simon (1898b) himself later transferred *Dolomedes* to Pisauridae. In 1898, Simon then classified Pisauridae into “Pisaurieae”, “Thalassieae”, and “Dolomedeae” based on the arrangement of their eyes. This classification was followed by Petrunkevitch (1928), but renamed as Pisaurinae, Thalassiinae (=Thalassinae), and Thaumasiinae (=Thaumasinae), respectively. Lehtinen (1967) was the first to propose the family Dolomedidae which included the elevation of *Dolomedes* and Thaumasiinae. This proposed taxonomic act was subsequently considered unjustified (Sierwald 1990; Griswold 1993). Although the subfamily placement of *Dolomedes* in Simon (1898b) and Petrunkevitch (1928) has been followed by some authors (Roewer 1955; Raven and Hebron 2018), the proximity of *Dolomedes* and *Thaumasia* has never been confirmed phylogenetically. Detailed studies and analyses of pisaurid genitalia seemed to support the monophyly of Pisauridae (Sierwald 1990; Griswold 1993; Zhang et al. 2004; Santos 2007b), with *Dolomedes* deemed a close relative of *Thalassius* Simon, 1885 (=*Nilus*) (Sierwald 1990; Zhang et al. 2004) or *Pisaurina* Simon, 1898 (Santos 2007b). Sierwald (1989, 1990) outlined some unique genital features in *Dolomedes*.

While some phylogenetic studies recovered results similar to the above morphological hypotheses (Bayer and Schönhofer 2013; Moradmand et al. 2014; Fernández et al. 2018), the majority suggested otherwise. Some studies still recovered a monophyletic Pisauridae, albeit with *Dolomedes* and the New Caledonian endemic *Bradystichus* Simon, 1884, if included, forming a clade sister to all other pisaurids (Wheeler et al. 2017; Cheng and Piel 2018; Piacentini and Ramírez 2019; Kallal et al. 2021). Other studies suggest that Pisauridae is paraphyletic (Polotow et al. 2015; Albo et al. 2017; Hazzi and Hormiga 2023; Kulkarni et al. 2023). They recover *Dolomedes* either as sister to a large clade with Pisauridae, Trechaleidae, and Lycosidae (Albo et al. 2017) or to Trechaleidae and Lycosidae (Hazzi and Hormiga 2023; Kulkarni et al. 2023). Kulkarni and co-workers (2023) recovered *Dolomedes* and *Bradystichus* as sister to Trechaleidae and Lycosidae.

Family Dolomedidae Simon, 1876, family rank resurrected

Bradystichidae Simon 1884, new synonymy

(*Raft spiders*)

### Composition

*Dolomedes* Latreille, 1804 (*Fishing spiders*), *Bradystichus* Simon, 1884, *Megadolomedes* Davies and Raven 1980, *Caledomedes* Raven and Hebron, 2018, *Mangromedes* Raven and Hebron, 2018, *Ornodolomedes* Raven and Hebron, 2018, and *Tasmomedes* Raven and Hebron, 2018.

### Diagnosis

Dolomedidae resemble Pisauridae, Trechaleidae, and Lycosidae. According to Simon (1876), these four families share the combination of the following habitus characteristics: i) carapace pear-shaped and longer than wide, dorsally elevated without tubercles or spines; ii) eight eyes in two rows: the posterior row strongly recurved with the lateral eyes distantly separated from the anterior eyes, all posterior eyes larger than the anterior eyes; iii) legs prograde with three claws, the fourth legs are the longest while the third legs are the shortest; iv) abdomen longer than wide; and v) ecribellate, spinnerets regular without elongated sections.

Dolomedidae genera can be distinguished from Trechaleidae, Lycosidae, and Pisauridae by the following characters: i) a tiny stem and head of spermathecae, which together form an accessory bulb, connected to the base of the spermathecae (Figs. S7–S9, after Sierwald 1989); ii) an elongated, coiled, or spirally arranged base of the spermatheca and spermathecal lumen (Figs. S7–S9; see also Remarks); iii) the presence of all three, the retrolateral tibial apophysis, the ventral tibial apophysis, and the basal cymbium apophysis on the male palp (Figs. S10–S11, see also Raven and Hebron 2018); v) the presence of a round sclerotized “saddle” instead of a distal tegular apophysis (Figs. S10–S11, see Sierwald 1990; Raven and Hebron 2018); and vi) the presence of a lateral subterminal apophysis, which is connected to the distal sclerotized tube of the apical division together with the embolus and fulcrum (Figs. S12).

Dolomedidae genera differ from Lycosidae and Trechaleidae by i) having both the copulatory duct and fertilization duct connected to the base of the spermatheca (Figs. S7–S9, see also Sierwald 1989); ii) the presence of a distal tegular projection, which forms a U-shaped tegular ring with the tegulum and conductor (Figs. S10–S11); and iii) the stereotypical behavior whereby females carry their egg sac on the chelicerae (as opposed to attached to the spinnerets in Lycosidae and in Trechaleidae).

Dolomedidae genera can also be diagnosed from the subfamily Pisaurinae by i) a nearly straight or strongly recurved, but never strongly procurved, anterior eye row (see Raven and Hebron 2018); ii) the carapace not protruding anteriorly at the anterior eye row in the lateral view (see Blandin 1979; Raven and Hebron 2018); iii) the absence of a “carina” in the female epigyne (Figs. S7–S8; see Sierwald 1997); iv) the copulatory and fertilization ducts both connected to the base of spermatheca (Figs. S7–S9, see also Sierwald 1989); v) a U-shaped tegular ring with a triangular conductor (Figs. S10–S11).

### Remarks

Female vulvae in Dolomedidae are usually elongated tubes whose structures are difficult to differentiate. Sierwald (1989) considered female *Dolomedes* to have a rather small base of spermatheca connected to a thick tubular fertilization duct (see Fig. 4 in Sierwald 1989); such nomenclature was followed subsequently (e.g., Yu and Kuntner 2024). However, further dissection into those tubular “fertilization ducts” detects matter that can be recognized as a sperm mass (Fig. S13). This newly emerged detail suggests that those tubular structures are responsible for sperm storage and should therefore be classified as the elongation of a base of spermatheca rather than a tubular fertilization duct (see also Jäger 2008). As in Sierwald (1989), we did not find any glandular pores on those tubular structures. In *Dolomedes* and in the investigated pisaurids, those pores are found exclusively on the tip of accessory bulbs or the head of spermathecae but not on its base (Sierwald 1989). The actual function of different parts of the female vulva in Dolomedidae awaits further detailed studies.

### Genus Blandinia Tonini et al., 2016, *incertae sedis*

#### Remarks

Blandin (1979) first established the genus as *Ransonia* Blandin, 1979, based on female specimens of a single species from Madagascar. The genus was renamed to *Blandinia* by Tonini et al. (2016) because *Ransonia* had been preoccupied by a jellyfish genus described by Kramp (1947). Blandin (1979) classified this genus in Pisauridae and Pisaurinae, owing to its strongly procurved anterior eye row. Blandin also pointed out that the genus had a distinct number of cheliceral teeth and a unique epigyne among Pisaurinae genera (Blandin 1979).

Our phylogenomic analyses suggest that *Blandinia* is not a pisaurid but is sister to Trechaleidae and Lycosidae. Although such a topology might suggest the establishment of a new family, we refrain from doing so due to unknown and conflicting topologies in Trechaleidae (Piacentini and Ramírez 2019; Hazzi and Hormiga 2023; Kulkarni et al. 2023), as well as the unknown male morphology of *Blandinia*. We therefore suggest to treat *Blandinia* as *incertae sedis* for the time being.

### Family Pisauridae Simon, 1890

#### Remarks

Based on our phylogeny (Fig. 3) Pisauridae should be delimited to *exclude* the genera in Dolomedidae (above) as well as *Blandinia*. This is the first step towards pisaurid delimitation. Although we refrain from providing a diagnosis for this family here, the current phylogenetic evidence (Fig. 3) suggests these 24 genera could be classified as pisaurids: *Afropisaura* Blandin, 1979, *Caripetella* Strand, 1928, *Charminus* Thorell, 1899, *Cispius* Simon, 1898, *Cladycnis* Simon, 1898, *Dendrolycosa* Doleschall, 1859, *Euprosthenops* Pocock, 1897, *Euprosthenopsis* Blandin, 1974, *Hala* Jocqué, 1994, *Hygropoda* Thorell, 1895, *Maypacius* Simon, 1898, *Nilus* O. Pickard-Cambridge, 1876, *Paracladycnis* Blandin, 1979, *Perenethis* L. Koch, 1878, *Pisaura* Simon, 1886, *Pisaurina* Simon, 1898, *Polyboea* Thorell, 1895, *Qianlingula* Zhang et al., 2004, *Rothus* Simon, 1898, *Sphedanus* Thorell, 1877, *Tallonia* Simon, 1889, *Tetragonophthalma* Karsch, 1878, *Thaumasia* Perty, 1833, and *Tolma* Jocqué, 1994.

The following 20 genera were not included in our phylogenetic analyses but may or may not be true pisaurids: *Archipirata* Simon, 1898, *Architis* Simon, 1898, *Chiasmopes* Pavesi, 1883, *Cispinilus* Roewer, 1955, *Conakrya* Schmidt, 1956, *Eucamptopus* Pocock, 1900, *Ilipula* Simon, 1903, *Inola* Davies, 1982, *Papakula* Strand, 1911, *Phalaeops* Roewer, 1955, *Stoliczka* O. Pickard-Cambridge, 1885, *Tapinothele* Simon, 1898, *Tapinothelella* Strand, 1909, *Tapinothelops* Roewer, 1955, *Thalassiopsis* Roewer, 1955, *Tinus* F. O. Pickard-Cambridge, 1901, *Voraptipus* Roewer, 1955, *Voraptus* Simon, 1898, *Vuattouxia* Blandin, 1979, and *Walrencea* Blandin, 1979.

## Supplementary Material

All supplementary material including the UCE matrices is available at *doi* and DRYAD, https://doi.org/10.5061/dryad.12jm63z6x.

## Acknowledgments

We thank Robert J. Raven for useful suggestions and kind help with materials and images; as well as Peter Michalik for the help in spider genial nomenclature. We thank Collin Chu, Volker Framenau, Nuria Macías Hernández, Kyle Knysh, Joseph Koh, Yuri Marusik, Sean McCann, Daniel Schoenberg, Michael Skvarla, and Lok Ming Tang for kindly providing materials, as well as Martín Ramírez, Lauren Esposito, Denise Montelongo, Petra Sierwald, Maureen Turcatel, Jessica Wadleigh, Dmitri Logunov, Michael Rix, Joseph Schubert, Owen Seeman, Julianne Waldock, Christophe Allard, and Julia Altmann for facilitating museum loans. We also thank Matjaž Bedjanič, Han-Po Chang, Matjaž Gregorič, Daiqin Li, Mu-Ming Lin, Ying-Yuan Lo, Annie Rasoanoeliarimanana, Rhina Harin’ Hala Rasolondalao, Jeremia Ravelojaona, Tiana Vololona, and Yi-Hua Yo for their support in the field. We would also like to express our thanks to Klemen Čandek, Yi-Yan Li, David Stanković, Eva Turk and an anonymous peer reader for bioinformatics suggestions and supports.

