## Supplementary table and figure for "Phylogenomics, Classification, and Lifestyle Evolution in Raft- and Nursery Web- Spiders (Araneae: Dolomedidae and Pisauridae)"

TABLE S1. Ultraconserved elements (UCE) loci generated in this study, captured by using the RTA spider probe set (Zhang et al. 2023). 95-CI: 95% confidence interval; bp: base pairs; kb: kilo base pairs (1,000 bp).

| File name | Contigs (loci) | Total bp | Mean length | 95 CI length | Min length | Max length | Median length | Contigs >1kb |
|---|---|---|---|---|---|---|---|---|
| Afropisaura-valida.unaligned.fasta | 2713 | 2722169 | 1003.379654 | 6.235615713 | 208 | 2524 | 1038 | 1467 |
| Aglaoctenus-oblongus.unaligned.fasta | 2850 | 2648750 | 929.3859649 | 5.053964879 | 208 | 1946 | 960 | 1234 |
| Blandinia-mahasoana.unaligned.fasta | 2795 | 3979563 | 1423.815027 | 8.034657238 | 207 | 3420 | 1476 | 2314 |
| Caripetella-madagascariensis.unaligned.fasta | 2871 | 3409747 | 1187.651341 | 7.319231711 | 207 | 3910 | 1234 | 2019 |
| Charminus-atomarius.unaligned.fasta | 2784 | 2795069 | 1003.975934 | 6.065345403 | 208 | 1957 | 1035 | 1500 |
| Cispius-sp.unaligned.fasta | 2791 | 2925867 | 1048.322107 | 6.21518479 | 211 | 2980 | 1069 | 1619 |
| Cladycnis-insignis.unaligned.fasta | 2870 | 3392519 | 1182.062369 | 6.899267472 | 209 | 3905 | 1216.5 | 2032 |
| Dendrolycosa-cf-songi.unaligned.fasta | 2767 | 3278901 | 1185.002168 | 7.294662127 | 209 | 2479 | 1232 | 1931 |

| | | | | | | | | |
|---|---|---|---|---|---|---|---|---|
| Dendrolycosa-rossi.unaligned.fasta | 2818 | 2462053 | 873.6880767 | 5.071219999 | 207 | 2166 | 894 | 1009 |
| Dolomedes-aff-horishanus-JP.unaligned.fasta | 2911 | 3231193 | 1109.99416 | 6.440801117 | 208 | 2624 | 1138 | 1894 |
| Dolomedes-aff-mizhoanus-SING.unaligned.fasta | 2861 | 2819004 | 985.3212164 | 5.912071655 | 208 | 2503 | 999 | 1425 |
| Dolomedes-aff-raptor-HK.unaligned.fasta | 2935 | 3318321 | 1130.603407 | 6.544440859 | 207 | 2805 | 1176 | 1980 |
| Dolomedes-aff-raptor-LoTW.unaligned.fasta | 2831 | 2790554 | 985.7131756 | 5.795255099 | 173 | 5572 | 1015 | 1472 |
| Dolomedes-aff-raptor-NETW.unaligned.fasta | 2911 | 3708580 | 1273.98832 | 7.194659922 | 207 | 6800 | 1326 | 2266 |
| Dolomedes-aff-raptoroides-CN.unaligned.fasta | 2901 | 3831793 | 1320.852465 | 7.930428724 | 210 | 4593 | 1380 | 2280 |
| Dolomedes-aff-raptor-SETW.unaligned.fasta | 2837 | 3130850 | 1103.577723 | 6.500908828 | 208 | 3569 | 1145 | 1845 |
| Dolomedes-aff-raptor-WTW.unaligned.fasta | 2866 | 2877120 | 1003.879972 | 6.089124171 | 207 | 3348 | 1038.5 | 1579 |
| Dolomedes-albineus.unaligned.fasta | 2924 | 3325669 | 1137.369699 | 6.674603905 | 210 | 5873 | 1167.5 | 1973 |
| Dolomedes-angustivirgatus.unaligned.fasta | 2692 | 1595254 | 592.5906389 | 4.304530697 | 101 | 1926 | 582 | 99 |
| Dolomedes-aquaticus.unaligned.fasta | 2912 | 4045698 | 1389.319368 | 7.786049975 | 207 | 7657 | 1443 | 2405 |

| | | | | | | | | |
|---|---|---|---|---|---|---|---|---|
| Dolomedes-bedjanic.unaligned.fasta | 2961 | 3733761 | 1260.979737 | 6.84956644 | 207 | 4081 | 1316 | 2294 |
| Dolomedes-bukhkaloi.unaligned.fasta | 2791 | 1710652 | 612.917234 | 3.518966498 | 173 | 2409 | 618 | 41 |
| Dolomedes-cf-petalinus-HK.unaligned.fasta | 2827 | 2027403 | 717.157057 | 4.614841009 | 208 | 2357 | 711 | 355 |
| Dolomedes-cf-raptoroides-HK.unaligned.fasta | 2869 | 2858313 | 996.2750087 | 6.2394552 | 207 | 3071 | 1018 | 1486 |
| Dolomedes-cf-raptoroides-VN.unaligned.fasta | 2812 | 1411484 | 501.9502134 | 2.915399077 | 209 | 2929 | 502 | 17 |
| Dolomedes-CONsp1.unaligned.fasta | 2789 | 3064654 | 1098.836142 | 6.473338646 | 211 | 3127 | 1129 | 1783 |
| Dolomedes-CONsp2.unaligned.fasta | 2881 | 2294026 | 796.2603263 | 5.160580227 | 207 | 1908 | 805 | 673 |
| Dolomedes-crosbyi.unaligned.fasta | 2859 | 3103711 | 1085.593214 | 6.284162978 | 210 | 2558 | 1115 | 1798 |
| Dolomedes-dondalei.unaligned.fasta | 2914 | 3077780 | 1056.20453 | 5.960259761 | 212 | 2761 | 1084 | 1728 |
| Dolomedes-facetus.unaligned.fasta | 2720 | 1445954 | 531.6007353 | 3.699975829 | 207 | 1313 | 516 | 60 |
| Dolomedes-fimbriatus.unaligned.fasta | 2878 | 3012918 | 1046.879083 | 6.479716476 | 207 | 3916 | 1075 | 1660 |
| Dolomedes-gracilipes.unaligned.fasta | 2852 | 2984177 | 1046.345372 | 6.337799064 | 208 | 2767 | 1074 | 1657 |

| | | | | | | | | |
|---|---|---|---|---|---|---|---|---|
| Dolomedes-gregoric.unaligned.fasta | 2794 | 3143037 | 1124.923765 | 6.635211974 | 210 | 2721 | 1160 | 1835 |
| Dolomedes-horishanus.unaligned.fasta | 2902 | 3713295 | 1279.564094 | 7.516434128 | 207 | 2755 | 1331 | 2213 |
| Dolomedes-hydatostella.unaligned.fasta | 2950 | 3447976 | 1168.805424 | 6.362911729 | 208 | 2739 | 1215.5 | 2106 |
| Dolomedes-japonicus.unaligned.fasta | 2890 | 2945499 | 1019.203806 | 5.827688225 | 207 | 3032 | 1057 | 1644 |
| Dolomedes-kalanoro.unaligned.fasta | 2947 | 3827840 | 1298.89379 | 6.838999951 | 214 | 2808 | 1348 | 2339 |
| Dolomedes-karijini.unaligned.fasta | 2875 | 4136518 | 1438.78887 | 8.233322074 | 212 | 3213 | 1507 | 2400 |
| Dolomedes-lesserti.unaligned.fasta | 2810 | 1318538 | 469.230605 | 2.338602362 | 208 | 1511 | 475 | 4 |
| Dolomedes-minor.unaligned.fasta | 2801 | 3368370 | 1202.5598 | 7.135371206 | 129 | 2571 | 1243 | 2019 |
| Dolomedes-mizhoanus.unaligned.fasta | 2905 | 3270905 | 1125.956971 | 6.33446254 | 207 | 2532 | 1158 | 1946 |
| Dolomedes-nigrimaculatus.unaligned.fasta | 2927 | 3897986 | 1331.734199 | 7.615265621 | 195 | 3049 | 1388 | 2332 |
| Dolomedes-okefinokensis.unaligned.fasta | 2909 | 3276702 | 1126.401513 | 6.349935491 | 209 | 2473 | 1176 | 1939 |
| Dolomedes-orion.unaligned.fasta | 2922 | 3809985 | 1303.896304 | 7.097897656 | 208 | 4591 | 1359 | 2325 |

| | | | | | | | | |
|---|---|---|---|---|---|---|---|---|
| Dolomedes-pegasus.unaligned.fasta | 2917 | 2977785 | 1020.83819 | 6.054830063 | 207 | 4390 | 1049 | 1625 |
| Dolomedes-plantarius.unaligned.fasta | 2885 | 3242359 | 1123.867938 | 6.676638379 | 207 | 3804 | 1149 | 1903 |
| Dolomedes-raptor.unaligned.fasta | 2937 | 3447585 | 1173.845761 | 6.823327769 | 210 | 7336 | 1213 | 2091 |
| Dolomedes-reuniascar.unaligned.fasta | 2952 | 3779960 | 1280.474255 | 7.00203439 | 209 | 3307 | 1336 | 2298 |
| Dolomedes-rotundus.unaligned.fasta | 2890 | 3524960 | 1219.709343 | 7.061408114 | 207 | 3097 | 1274 | 2096 |
| Dolomedes-saganus.unaligned.fasta | 2828 | 1789517 | 632.7853607 | 4.115825771 | 115 | 2408 | 633.5 | 123 |
| Dolomedes-schauinslandi.unaligned.fasta | 2628 | 1436361 | 546.5605023 | 4.064050514 | 204 | 2171 | 530 | 61 |
| Dolomedes-scriptus.unaligned.fasta | 2706 | 1680050 | 620.8610495 | 4.364225566 | 175 | 2158 | 609 | 122 |
| Dolomedes-senilis.unaligned.fasta | 2842 | 2849919 | 1002.786418 | 5.982893564 | 207 | 4274 | 1016.5 | 1488 |
| Dolomedes-silvicola.unaligned.fasta | 2893 | 2802102 | 968.5800207 | 5.664900003 | 207 | 3323 | 989 | 1405 |
| Dolomedes-spathularis.unaligned.fasta | 2937 | 2312319 | 787.3064351 | 4.242519735 | 207 | 2295 | 807 | 474 |
| Dolomedes-striatus.unaligned.fasta | 2713 | 1511397 | 557.0943605 | 3.414177501 | 207 | 1371 | 555 | 21 |

| | | | | | | | | |
|---|---|---|---|---|---|---|---|---|
| Dolomedes-sulfureus.unaligned.fasta | 2866 | 2881311 | 1005.342289 | 5.932763196 | 207 | 2835 | 1031 | 1566 |
| Dolomedes-tenebrosus.unaligned.fasta | 2898 | 3445239 | 1188.833333 | 6.349892066 | 219 | 3693 | 1224 | 2079 |
| Dolomedes-transfuga.unaligned.fasta | 2343 | 1326769 | 566.2693128 | 4.419438737 | 207 | 1708 | 555 | 53 |
| Dolomedes-triton.unaligned.fasta | 2301 | 819802 | 356.2807475 | 2.065246342 | 207 | 750 | 341 | 0 |
| Dolomedes-vittatus.unaligned.fasta | 2949 | 2848154 | 965.8033232 | 5.431321437 | 207 | 2716 | 987 | 1431 |
| Dolomedes-yawatai.unaligned.fasta | 2515 | 879320 | 349.6302187 | 4.103832182 | 207 | 9767 | 342 | 3 |
| Euprosthenops-australis.unaligned.fasta | 2841 | 3563463 | 1254.298838 | 7.617787174 | 209 | 2861 | 1304 | 2105 |
| Euprosthenopsis-sp.unaligned.fasta | 2693 | 2572697 | 955.3275158 | 6.124860559 | 212 | 2561 | 978 | 1273 |
| Fecenia-cylindrata.unaligned.fasta | 2829 | 2669510 | 943.6231884 | 6.113822865 | 207 | 3707 | 954 | 1266 |
| Hala-sp.unaligned.fasta | 2824 | 3292623 | 1165.942989 | 7.191027781 | 209 | 3348 | 1211 | 1950 |
| Hogna-radiata.unaligned.fasta | 2345 | 1380117 | 588.5360341 | 4.574925546 | 207 | 1960 | 577 | 82 |
| Hygropoda-higenaga.unaligned.fasta | 2788 | 3544757 | 1271.433644 | 7.597445197 | 211 | 3533 | 1319 | 2110 |

| | | | | | | | | |
|---|---|---|---|---|---|---|---|---|
| Hygropoda-MADsp.unaligned.fasta | 2881 | 3010518 | 1044.955918 | 6.12867214 | 208 | 2037 | 1060 | 1651 |
| Maypacius-bilineatus.unaligned.fasta | 2889 | 3070174 | 1062.711665 | 6.262637082 | 207 | 2657 | 1085 | 1693 |
| Megadolomedes-johndouglasi.unaligned.fasta | 2842 | 1605899 | 565.0594652 | 3.048022248 | 208 | 1475 | 572.5 | 18 |
| Neoctenus-sp.unaligned.fasta | 2817 | 3763931 | 1336.14874 | 8.143259331 | 210 | 2799 | 1400 | 2207 |
| Nilus-majungensis.unaligned.fasta | 2848 | 2836446 | 995.943118 | 5.694825203 | 183 | 2687 | 1017 | 1496 |
| Nilus-phipsoni.unaligned.fasta | 2722 | 2753362 | 1011.521675 | 6.501279391 | 207 | 2521 | 1045 | 1489 |
| Ornodolomedes-sp.unaligned.fasta | 2931 | 3311077 | 1129.674855 | 5.934808144 | 207 | 2377 | 1168 | 1990 |
| Paracladycnis-vis.unaligned.fasta | 2894 | 3532566 | 1220.651693 | 6.825936823 | 208 | 3098 | 1263.5 | 2150 |
| Pardosa-lugubris.unaligned.fasta | 2738 | 3002938 | 1096.763331 | 6.854210774 | 205 | 5159 | 1110.5 | 1688 |
| Perenethis-fascigera.unaligned.fasta | 2858 | 3877179 | 1356.605668 | 8.059295206 | 210 | 3294 | 1413 | 2290 |
| Peucetia-MADsp.unaligned.fasta | 1125 | 679560 | 604.0533333 | 9.046019806 | 207 | 1588 | 540 | 149 |
| Pirata-piscatorius.unaligned.fasta | 2874 | 3534149 | 1229.696938 | 6.773095041 | 209 | 3682 | 1269 | 2157 |

| | | | | | | | | |
|---|---|---|---|---|---|---|---|---|
| Pisaura-mirabilis.unaligned.fasta | 2846 | 3513184 | 1234.428672 | 7.538780847 | 211 | 3417 | 1276 | 2069 |
| Pisauridae-gen-MADsp.unaligned.fasta | 2810 | 2941071 | 1046.644484 | 6.383071534 | 208 | 2947 | 1068 | 1598 |
| Pisaurina-mira.unaligned.fasta | 2881 | 2648517 | 919.3047553 | 5.113769295 | 207 | 2242 | 950 | 1216 |
| Polyboea-zonaformis.unaligned.fasta | 2815 | 2262771 | 803.8262877 | 5.513698859 | 199 | 2561 | 818 | 753 |
| Psechrus-sp.unaligned.fasta | 2890 | 2811727 | 972.915917 | 5.882095555 | 211 | 2346 | 994.5 | 1424 |
| Qianlingula-cf-turbinata.unaligned.fasta | 2443 | 1333103 | 545.6827671 | 4.286451178 | 178 | 1447 | 523 | 58 |
| Rothus-vittatus.unaligned.fasta | 2857 | 3487299 | 1220.615681 | 6.993847602 | 179 | 3405 | 1266 | 2102 |
| Sphedanus-quadrimaculatus.unaligned.fasta | 2299 | 922400 | 401.2179208 | 3.018808046 | 207 | 1224 | 365 | 6 |
| Tallonia-picta.unaligned.fasta | 2716 | 2358810 | 868.4867452 | 5.072542759 | 207 | 1492 | 903 | 951 |
| Tetragonophthalma-vulpina.unaligned.fasta | 2688 | 2189102 | 814.3980655 | 5.245444047 | 207 | 2461 | 831.5 | 703 |
| Thaumasia-velox.unaligned.fasta | 2798 | 3763496 | 1345.066476 | 8.033331506 | 207 | 3228 | 1419.5 | 2205 |
| Tolma-sp.unaligned.fasta | 2845 | 3625483 | 1274.334974 | 7.935903355 | 208 | 4188 | 1318 | 2137 |

| | | | | | | | | |
|---|---|---|---|---|---|---|---|---|
| Trochosa-aff-robusta.unaligned.fasta | 2853 | 2960507 | 1037.682089 | 6.314586463 | 209 | 2694 | 1057 | 1596 |
| Viridasius-sp.unaligned.fasta | 2682 | 2435223 | 907.9876957 | 5.312677929 | 209 | 2824 | 934 | 1101 |
| Xysticus-sp.unaligned.fasta | 2373 | 1605143 | 676.4193005 | 5.161879506 | 110 | 2324 | 669 | 235 |
| Zoropsis-spinimana.unaligned.fasta | 2675 | 2750603 | 1028.262804 | 6.718675771 | 209 | 4071 | 1054 | 1483 |

55

### Supplementary figure captions

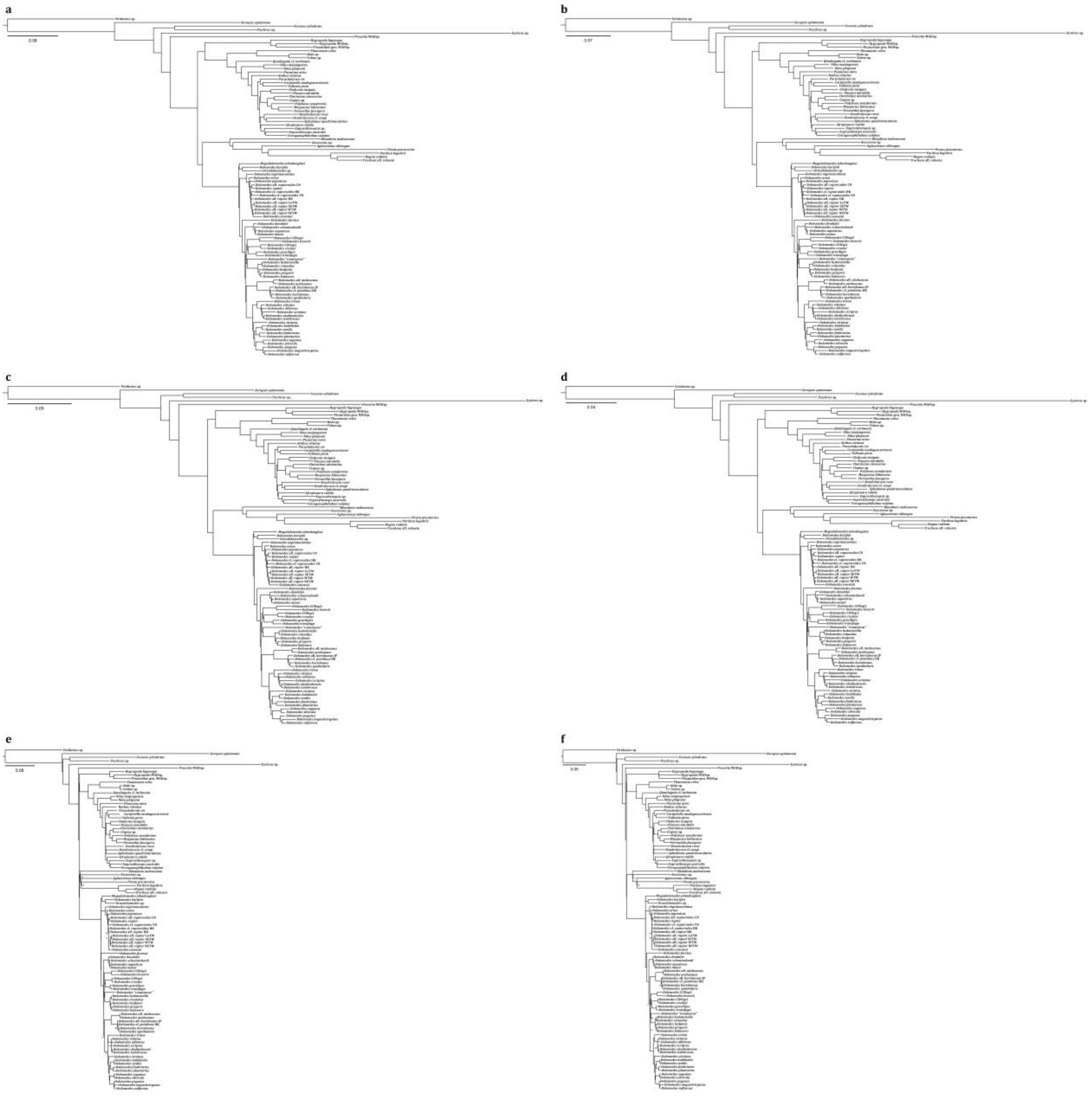


Figure S1. Phylogenies reconstructed using: a–b, maximum likelihood on (a) UCE loci with at least 75% taxon completeness (sparse matrix) and (b) UCE loci with at least 95% taxon completeness (dense matrix); c–d, Bayesian inference on (c) sparse matrix and (d) dense matrix; as well as e–f, multi-species coalescent on (e) sparse matrix and (f) dense matrix. See also supplementary materials for the raw tree files with nodal supports.

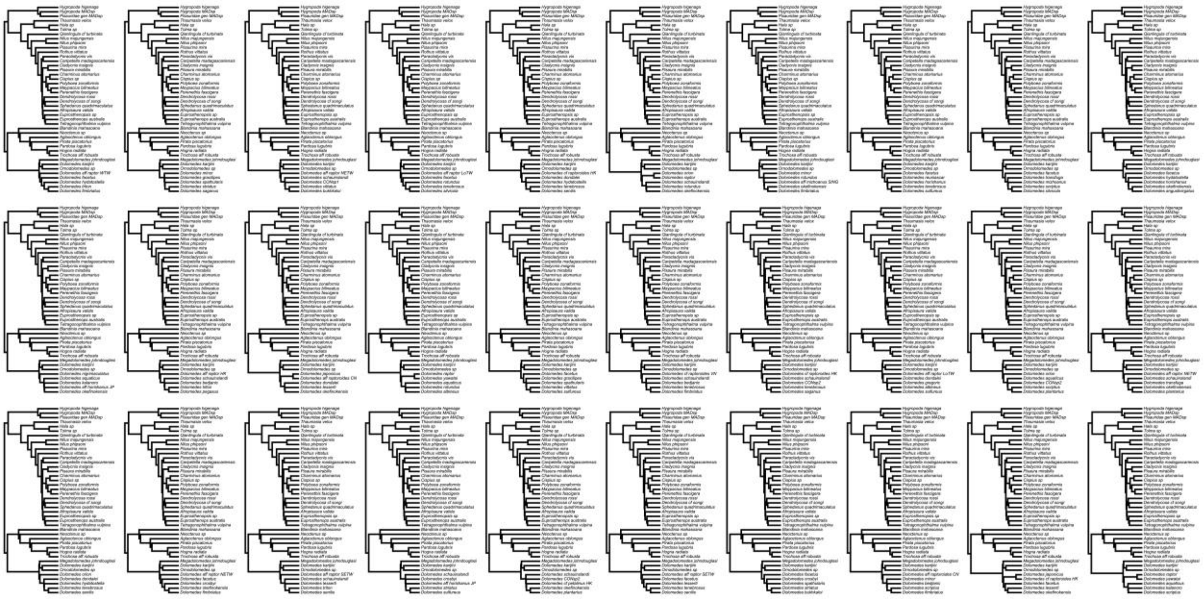

FIGURE S2. Topologies of the 30 reduced taxon trees with *Dolomedes* Latreille, 1804 terminals randomly pruned by biogeographical realms.

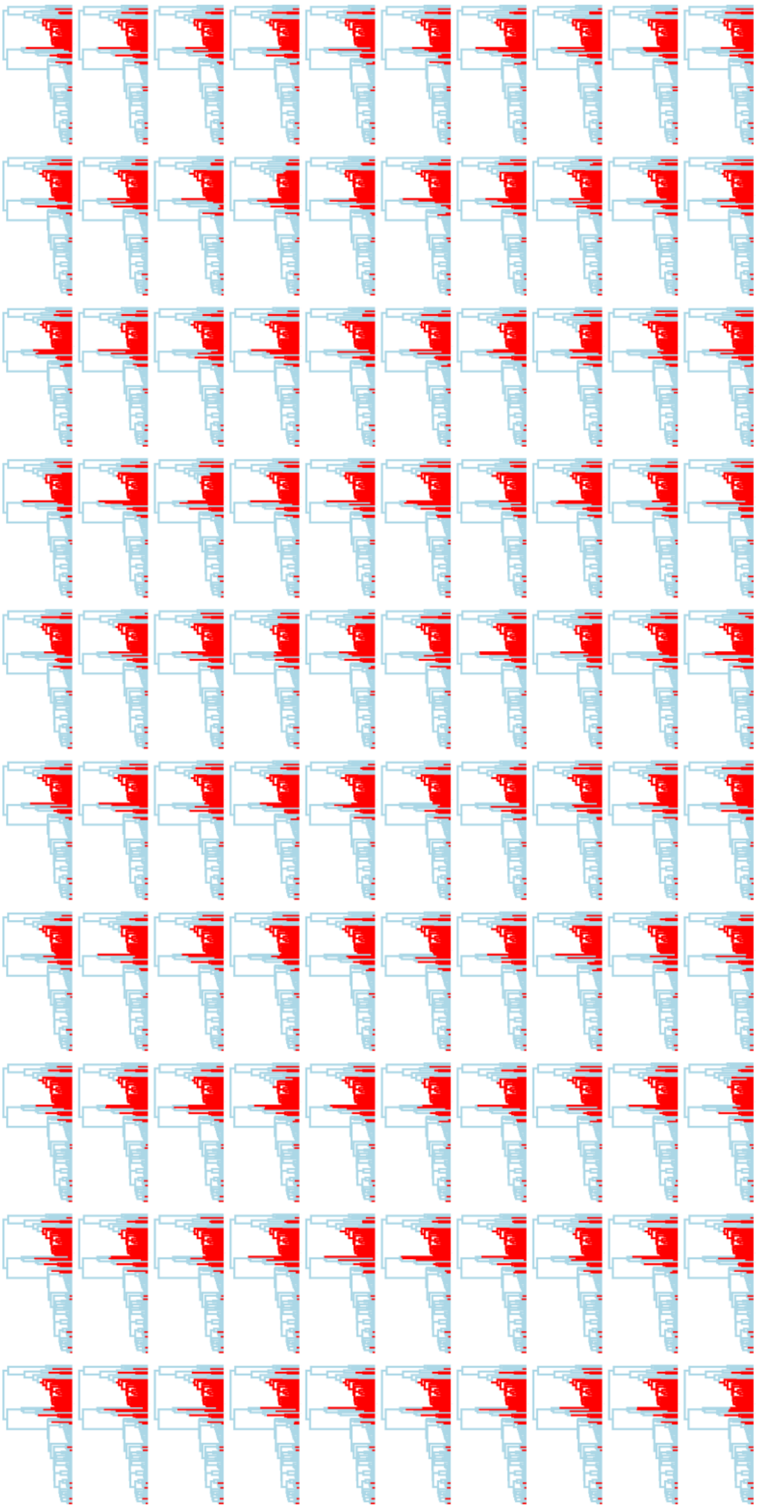

FIGURE S3. All 100 stochastic character simulations of ancestral state reconstruction of semi-aquatic (blue) versus terrestrial (red) lifestyle on the full tree using trait transitional probabilities estimated and averaged from the 30 reduced taxon trees.

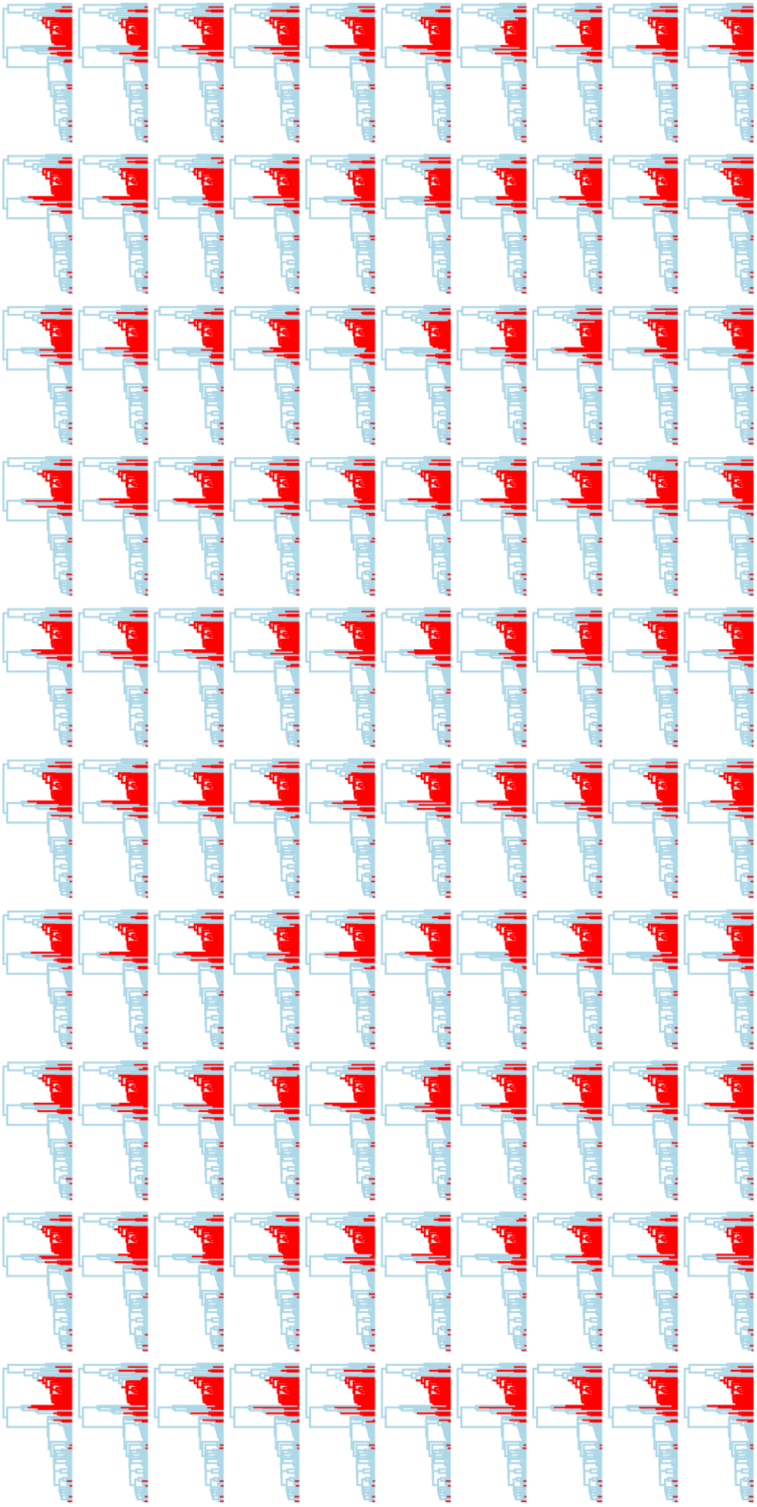

FIGURE S4. All 100 stochastic character simulations of ancestral state reconstruction of semi-aquatic (blue) versus terrestrial (red) lifestyle on the full taxon tree using trait transitional probabilities estimated from the full taxon tree.

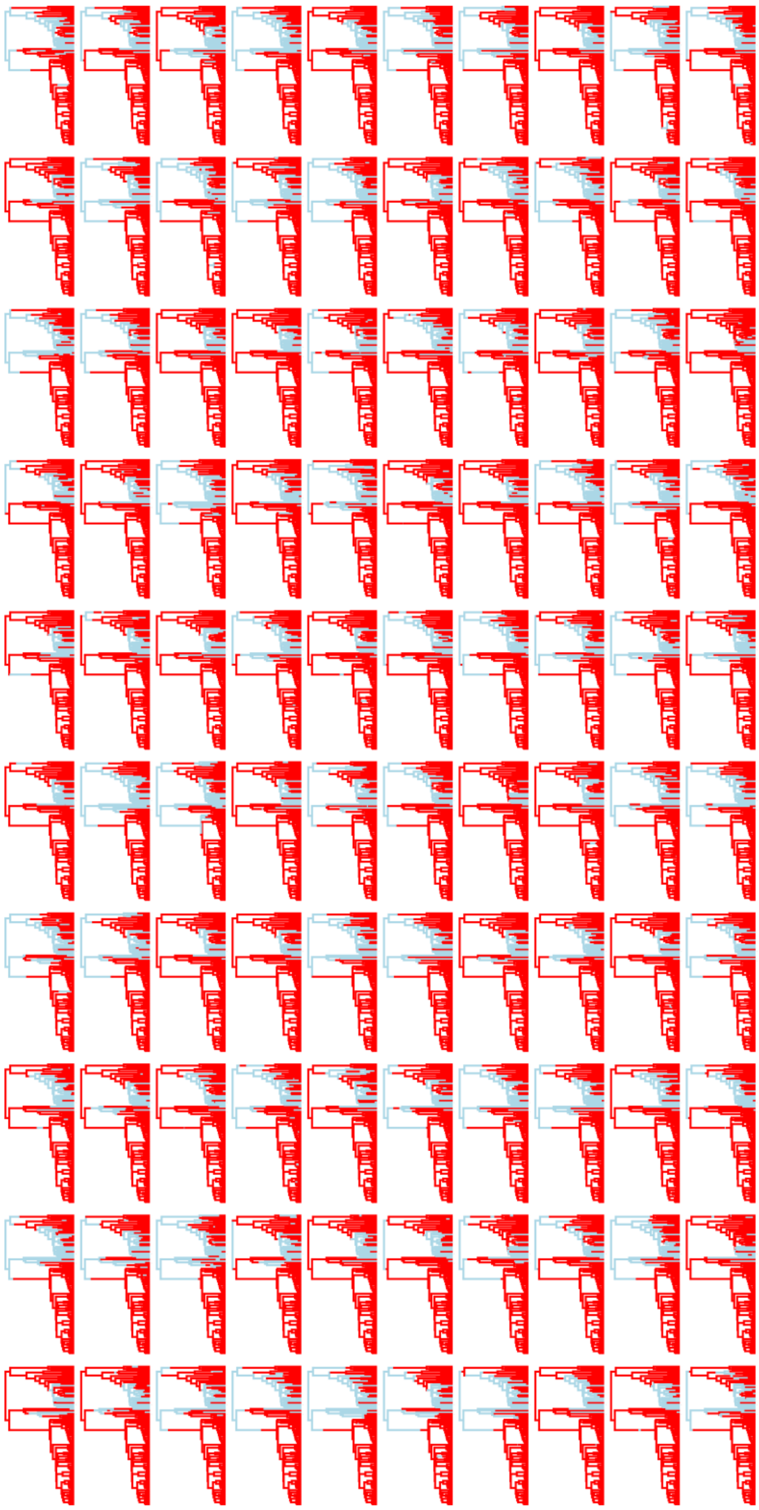

FIGURE S5. All 100 stochastic character simulations of ancestral state reconstruction of capture web presence (blue) or absence (red) on the full tree using trait transitional probabilities estimated and averaged from the 30 reduced taxon trees.

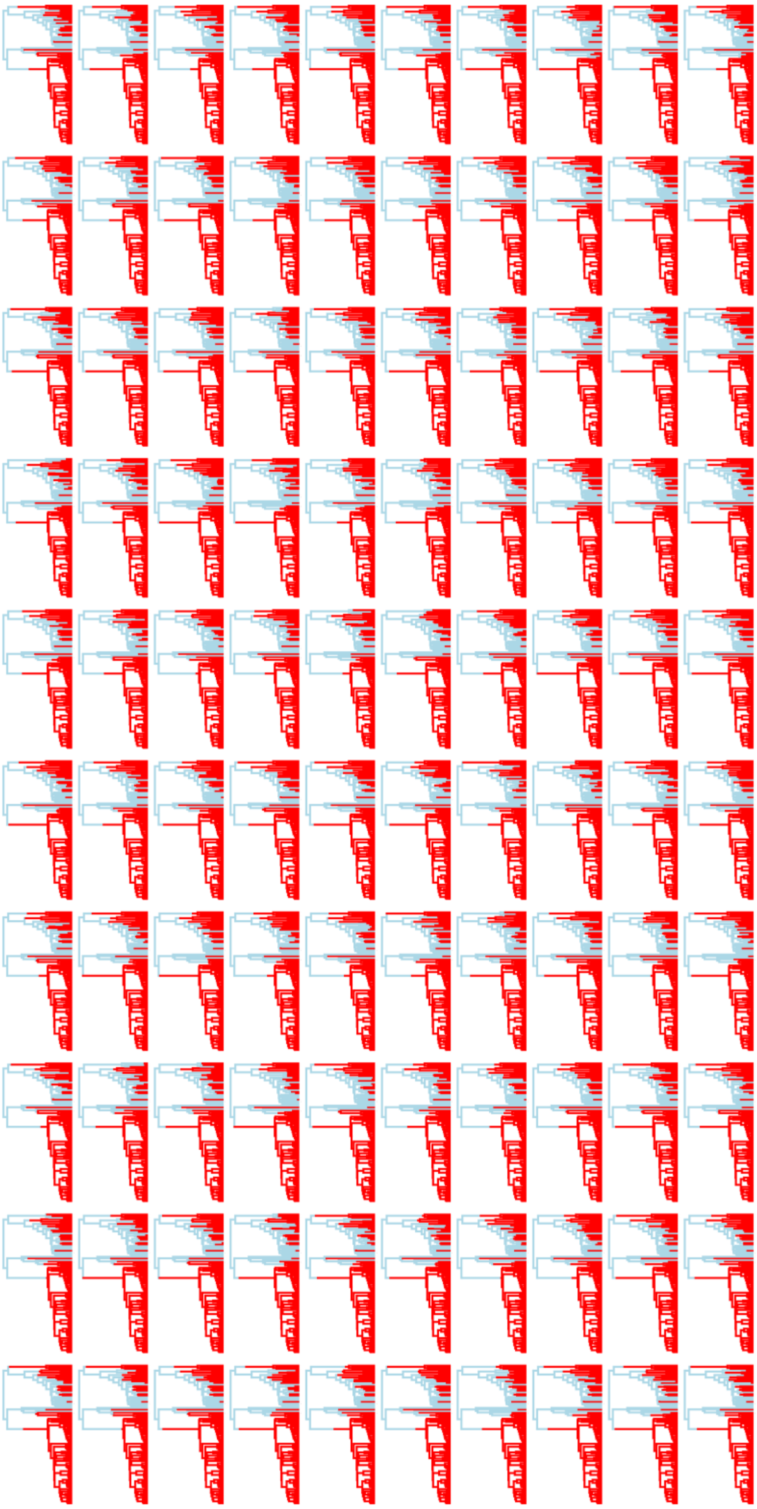

FIGURE S6. All 100 stochastic character simulations of ancestral state reconstruction of capture web presence (blue) or absence (red) on the full tree using trait transitional probabilities estimated from the full taxon tree.

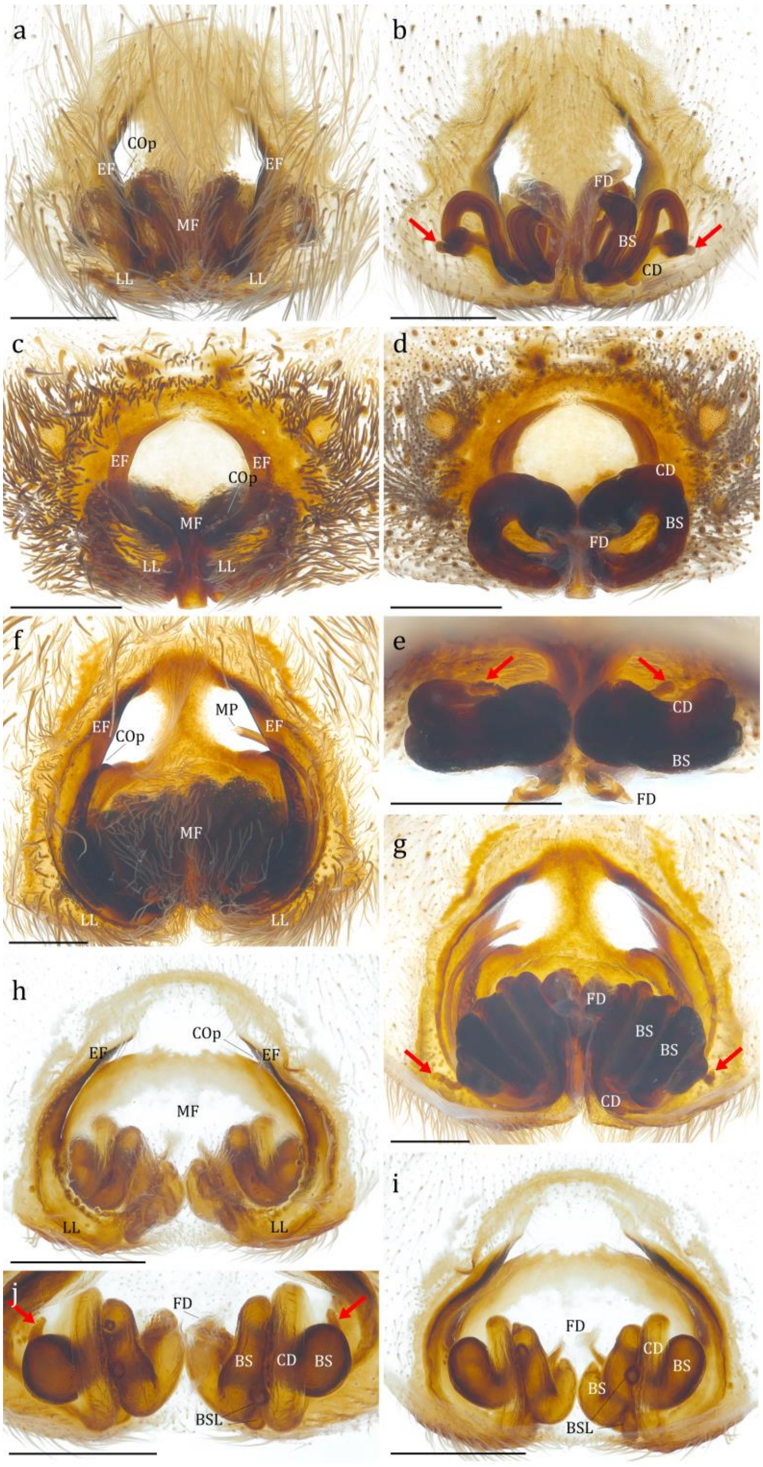

FIGURE S7. Female genitalia of *Dolomedes* Latreille, 1804, *Bradystichus* Simon, 1884, *Megadolomedes* Davies and Raven, 1980, and *Caledomedes* Raven and Hebron, 2018, showing the family diagnostic characteristics of Dolomedidae Simon, 1876, notably the accessory bulb marked in each genus with red arrows; a–b, *D. fimbriatus* (Clerck, 1757) (KPARA_00297): (a) epigyne, ventral view, (b) *idem*, dorsal view; c–e, *B. calligaster* Simon, 1884 (CASENT_9114936): (c) epigyne, ventral view, (d) *idem*, dorsal view, (e) vulva, anterior view; f–g, *Me. trux* (Lamb, 1911) (CASENT_9114937): (f) epigyne, ventral view, (g) *idem*, dorsal view; h–j, *C. flavovittatus* (Simon, 1880) (QM_NewCaledonia8641): (h) epigyne, ventral view, (i) *idem*, dorsal view, (j) vulva, anterior view. BS: base of spermatheca; BSL: lobe of base of spermatheca; CD: copulatory duct; COp: copulatory opening; EF: epigynal fold; FD: fertilization duct; MF: middle field; MP: mating plug of male.

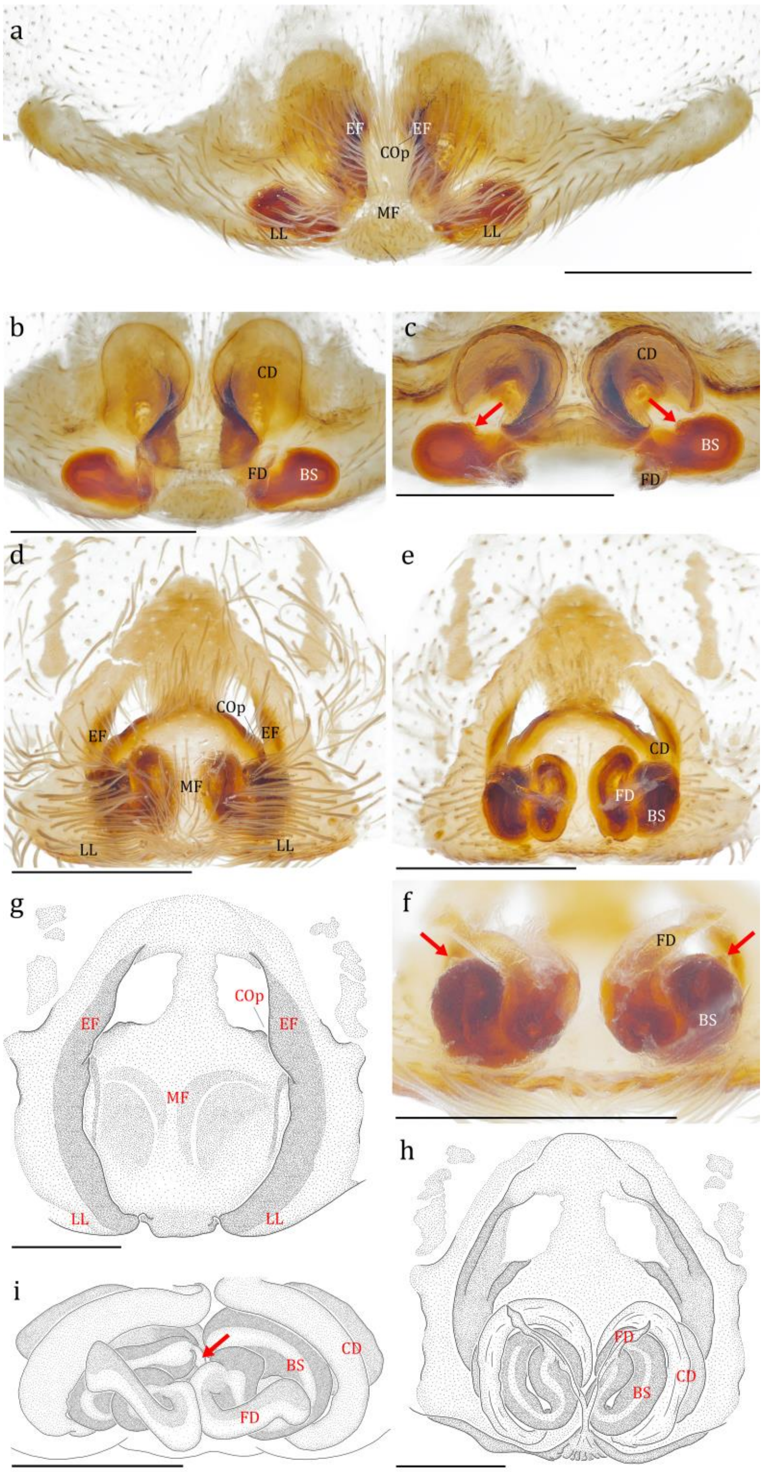


FIGURE S8. Female genitalia of *Mangromedes* Raven and Hebron, 2018, *Ornodolomedes* Raven and Hebron, 2018, and *Tasmomedes* Raven and Hebron, 2018, showing the family diagnostic characteristics of Dolomedidae Simon, 1876, notably the accessory bulb marked in each genus with red arrows. a–c, *Ma. kochi* (Roewer, 1951) (QM_S108067): (a) epigyne, ventral view, (b) vulva, dorsal view, (c) *idem*, anterior view; d–f, *O.* cf. *nicholsoni* Raven and Hebron, 2018 (WAM_T140666): (d) epigyne, ventral view, (e) *idem*, dorsal view, (f) vulva, posterior view; g–i, *T. eberhardarum* (Strand 1913) (g–h, redrawn from Raven and Hebron 2018; i, QM_S67787, credit of the original photo: Robert J. Raven): (g) epigyne, ventral view, (h) *idem*, dorsal view, (i), vulva, anterior view. BS: base of spermatheca; CD: copulatory duct; COp: copulatory opening; EF: epigynal fold; FD: fertilization duct; MF: middle field.

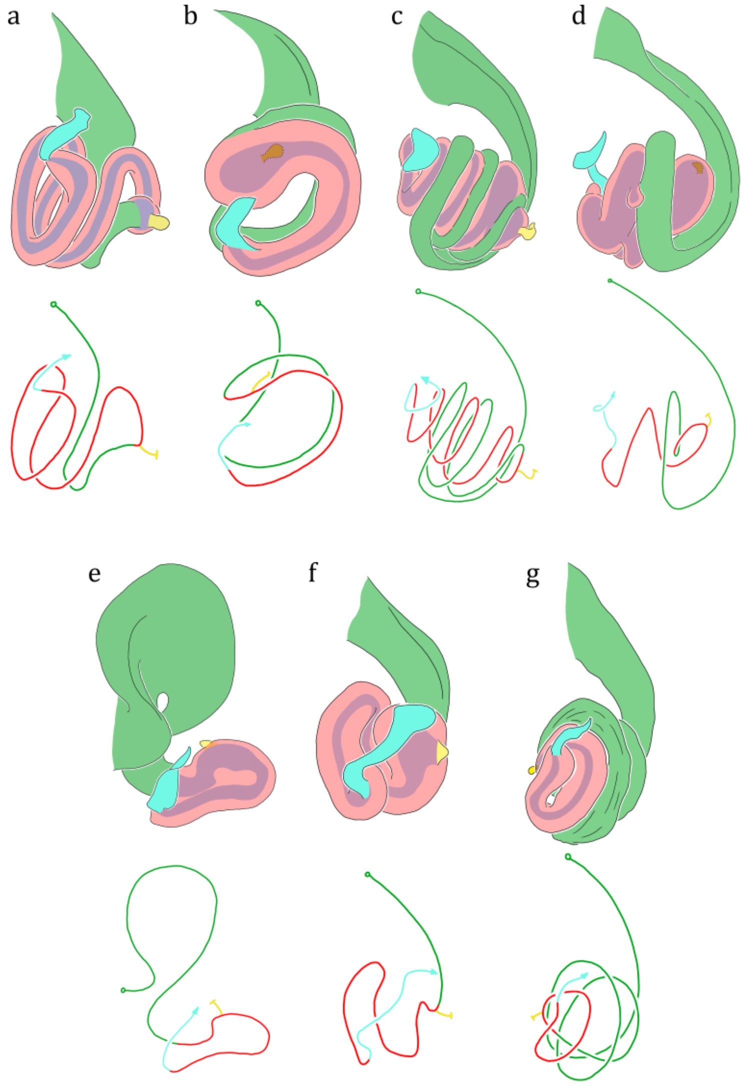


FIGURE S9. Illustration of the left vulva of seven Dolomedidae Simon, 1876 genera in dorsal view with color codon showing connection between structures. (a) *Dolomedes fimbriatus* (Clerck, 1757) (KPARA_00297); (b) *Bradystichus calligaster* Simon, 1884 (CASENT_9114936); (c) *Megadolomedes trux* (Lamb, 1911) (CASENT_9114937); (d) *Caledomedes flavovittatus* (Simon, 1880) (QM_NewCaledonia8641); (e) *Mangromedes kochi* (Roewer, 1951) (QM_S108067); (f) *Ornodolomedes* cf. *nicholsoni* Raven and Hebron, 2018 (WAM_T140666); (g) *Tasmomedes eberhardarum* (Strand, 1913) (redrawn from Raven and Hebron 2018). Green: copulatory duct; red: base of spermatheca; purple: spermatheca lumen;

yellow: accessory bulb; cyan: fertilization duct.

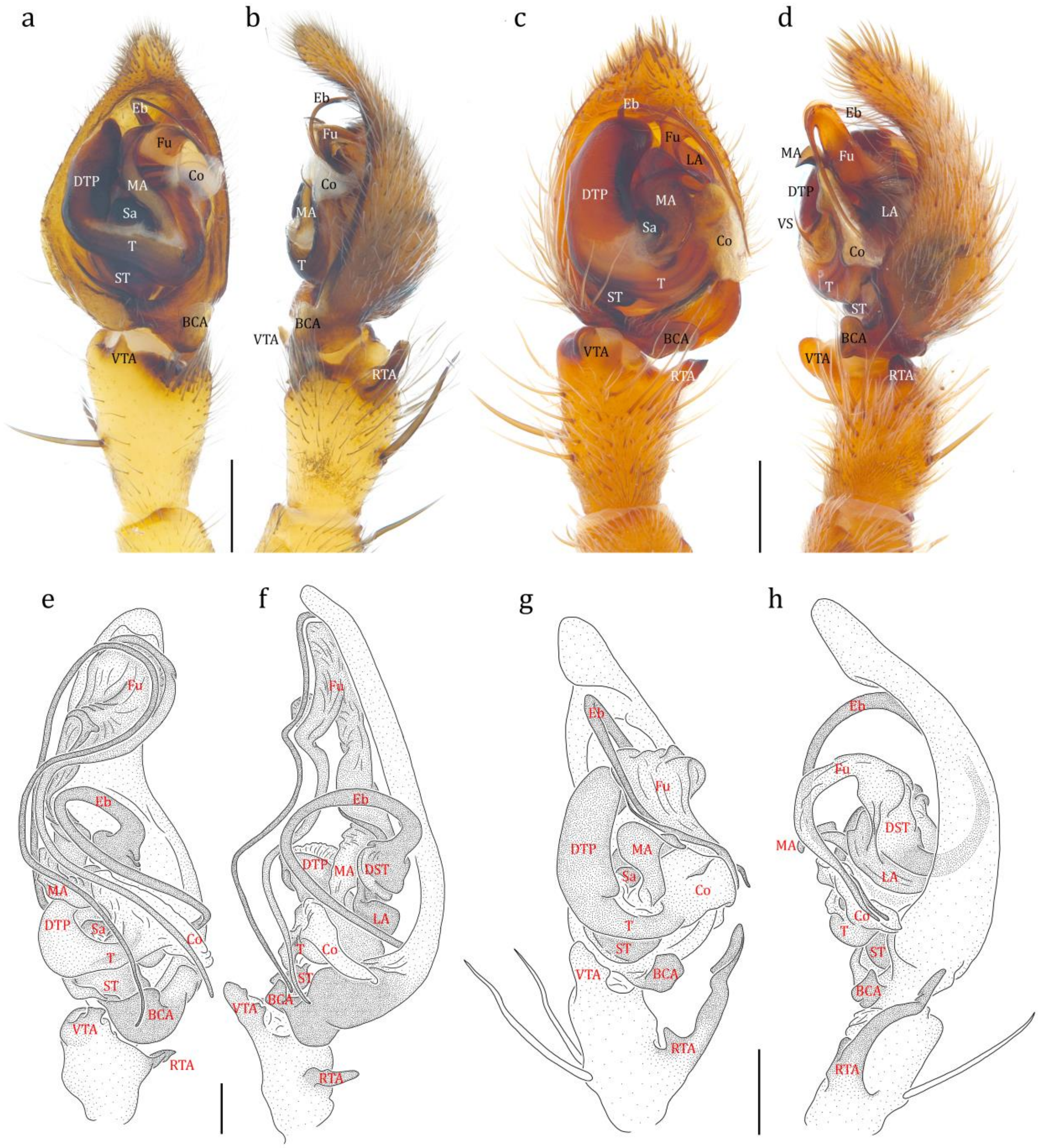


FIGURE S10. Male genitalia of *Dolomedes* Latreille, 1804, *Bradystichus* Simon, 1884, *Megadolomedes* Davies and Raven, 1980, and *Caledomedes* Raven and Hebron, 2018, showing the family diagnostic characteristics of Dolomedidae Simon, 1876. a–b, *D. fimbriatus* (Clerck, 1757) (KPARA_00260): (a) left palp, ventral view, (b) *idem*, retrolateral view; c–d, *B.* cf. *aoupinie* Platnick and Forster, 1993 (QM_S109746): (c) left palp, ventral view, (d) *idem*, retrolateral view; e–f, *Me. australianus* (L. Koch, 1865) (redrawn from Raven and Hebron 2018): (e) left palp, ventral view, (f) *idem*, retrolateral view; g–h, *C. flavovittatus* (Simon, 1880) (redrawn from Raven and Hebron 2018): (g) left palp, ventral view, (h) *idem*, retrolateral view. BCA: basal cymbium apophysis; Co: conductor; DTP: distal tegular projection; Eb: embolus; Fu: fulcrum; LA: lateral subterminal apophysis; MA: median apophysis; RTA: retrolateral tibial apophysis; Sa: saddle; ST: subtegulum; T: tegulum; VS: ventral spine of median apophysis; VTA: ventral tibial apophysis.

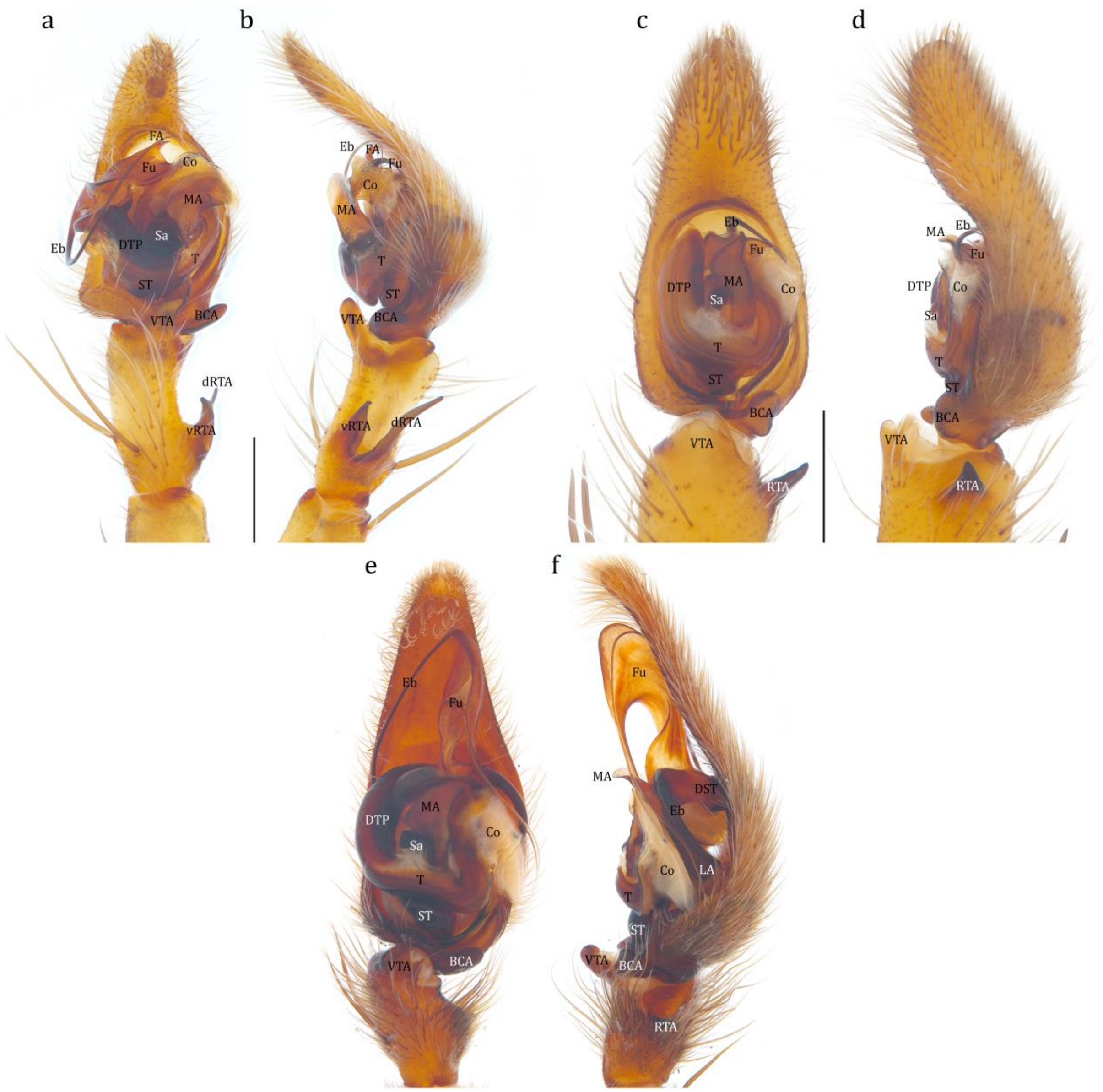


FIGURE S11. Male genitalia of *Mangromedes* Raven and Hebron, 2018, *Ornodolomedes* Raven and Hebron, 2018, and *Tasmomedes* Raven and Hebron, 2018, showing the family diagnostic characteristics of Dolomedidae Simon, 1876. a–b, *Ma. kochi* (Roewer, 1951) (QM_S108067): (a) left palp, ventral view, (b) *idem*, retrolateral view; c–d, *O.* cf. *nicholsoni* Raven and Hebron, 2018 (FMNH_128417): (c) left palp, ventral view, (d) *idem*, retrolateral view; e–f, *T. eberhardarum* (Strand, 1913) (QM_S70376): (e) right palp, mirrored, ventral view, (f) *idem*, mirrored, retrolateral view. BCA: basal cymbium apophysis; Co: conductor;

dRTA: dorsal retrolateral tibial apophysis; DTP: distal tegular projection; Eb: embolus; FA: fulcrum apophysis Fu: fulcrum; LA: lateral subterminal apophysis; MA: median apophysis; RTA: retrolateral tibial apophysis; Sa: saddle; ST: subtegulum; T: tegulum; vRTA: ventral retrolateral tibial apophysis; VTA: ventral tibial apophysis.

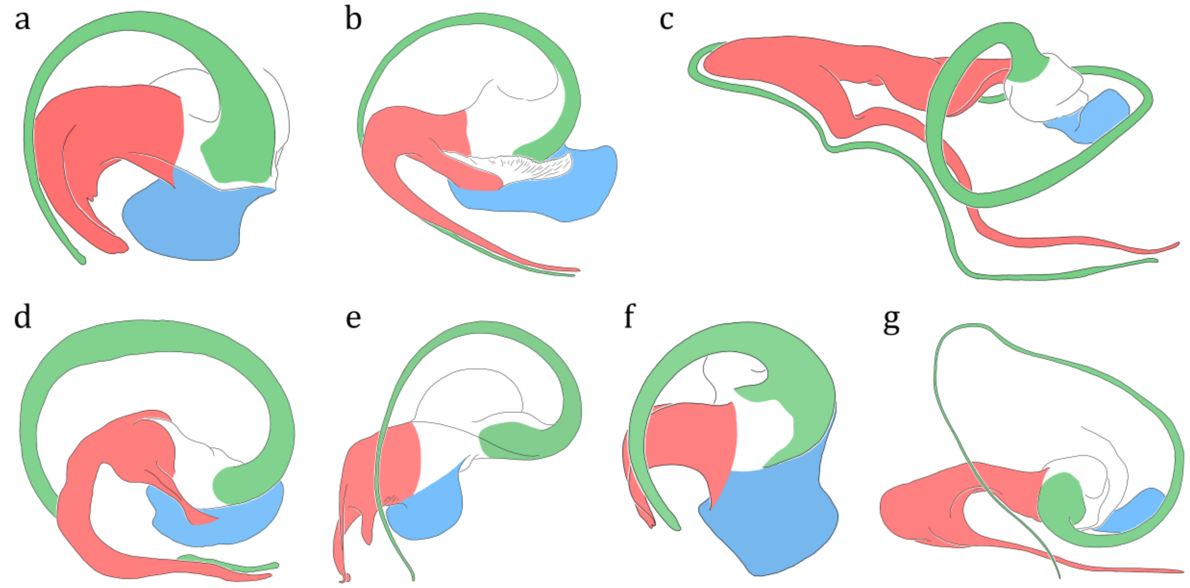


FIGURE S12. Illustration of the distal sclerotized tube of the apical division of the seven Dolomedidae Simon, 1876 genera in retrolateral view with color codon highlighting homologous sclerites. (a) *Dolomedes fimbriatus* (Clerck, 1757) (KPARA_00260), left palp; (b) *Bradystichus* cf. *aoupinie* Platnick and Forster, 1993 (QM_S109746), left palp; (c) *Megadolomedes australianus* (L. Koch, 1865) (redrawn from Raven and Hebron 2018), left palp; (d) *Caledomedes flavovittatus* (Simon, 1880) (redrawn from Raven and Hebron 2018), left palp; (e) *Mangromedes kochi* (Roewer, 1951) (QM_S108067), left palp; (f) *Ornodolomedes* cf. *nicholsoni* Raven and Hebron, 2018 (FMNH_128417), left palp; (g)

*Tasmomedes eberhardarum* (Strand, 1913) (QM_S70376), right palp, mirrored.

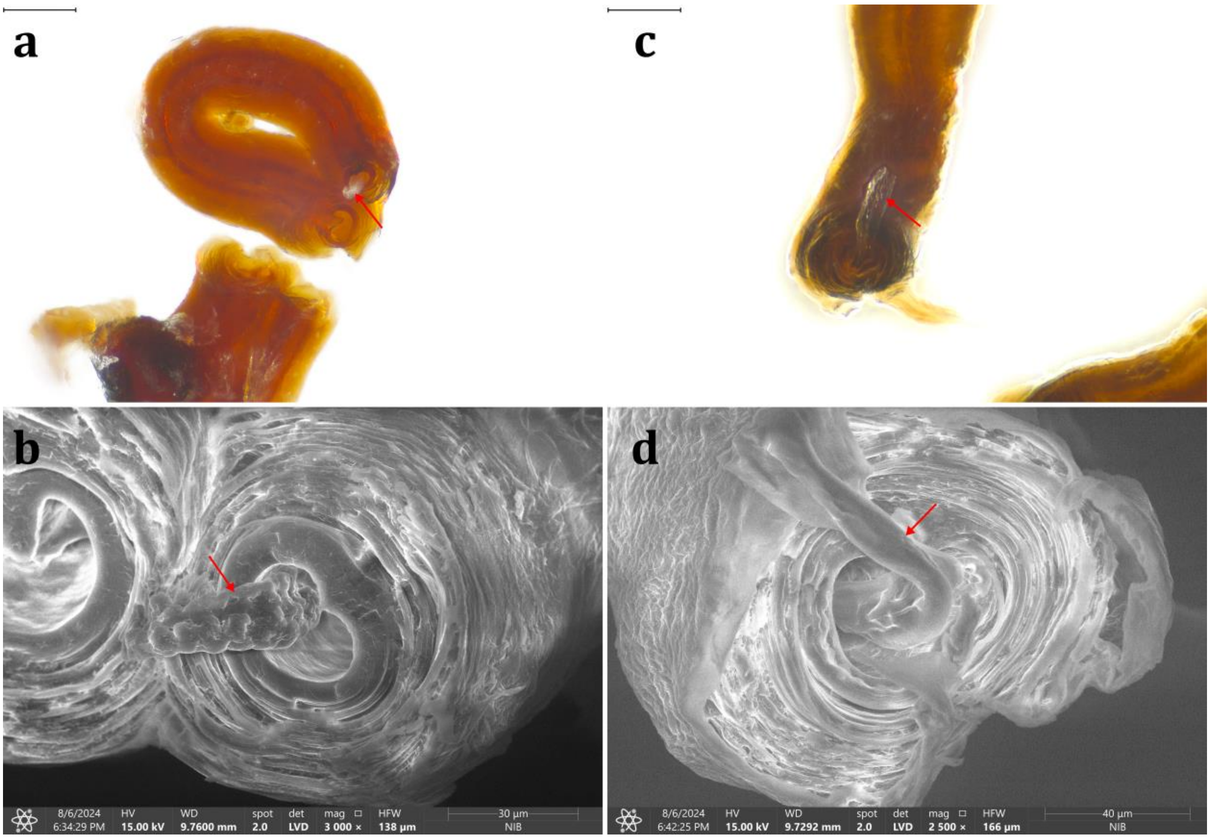


FIGURE S13. Sperm masses (red arrows) found in the "fertilization duct" (*sensu* Sierwald 1989), suggesting these tubular structures should be the elongated base of spermathecae. a–b, *Dolomedes fimbriatus* (Clerck, 1757), KPARA_00014; c–d, *D. scriptus* Hentz, 1845, KPARA_00118.

Hentz N.M. 1845. Descriptions and figures of the araneides of the United States. Boston Journal of Natural History. 5:189–202.

Koch L. 1865. Beschreibungen neuer Arachniden und Myriopoden. Verhandlungen der Kaiserlich-Königlichen Zoologisch-Botanischen Gesellschaft in Wien . 15:857–892.

Lamb J. 1911. Descriptions of some new Queensland Araneidae. Annals of the Queensland Museum. 10:169–174.

Latreille P.A. 1804. Tableau methodique des Insectes. Nouveau Dictionnaire d'Histoire Naturelle Paris. 24:129–295.

Platnick N.I., Forster R.R. 1993. A revision of the New Caledonian spider genus Bradystichus (Araneae, Lycosoidea). American Museum Novitates. 3075:1–14.

Raven R., Hebron W. 2018. A review of the water spider family Pisauridae in Australia and New Caledonia with descriptions of four new genera and 23 new species. Memoirs of the Queensland Museum. Nature 60:233–381.

Roewer C.F. 1951. Neue Namen einiger Araneen-Arten. Abhandlungen des Naturwissenschaftlichen Vereins zu Bremen. 32:437–456.

Sierwald P. 1989. Morphology and ontogeny of female copulatory organs in American Pisauridae, with special reference to homologous features (Arachnida: Araneae). Smithsonian Contributions to Zoology. 484:1–24.

Simon E. 1876. Les arachnides de France. Tome troisième. Paris, France: Roret.

Simon E. 1880. Matériaux pour servir à une faun arachnologique de la Nouvelle Calédonie. Annales de la Société Entomologique de Belgique. 23:164–175.

Simon E. 1884. Description d'une nouvelle famille de l'ordre des Araneae (Bradystichidae). Annales de la Société Entomologique de Belgique. 28:297–301.

Strand E. 1913. Drei neue Spinnen von Victoria in Australien. Jahrbücher des Nassauischen Vereins für Naturkunde. 66:203–209.

Zhang J., Li Z., Lai J., Zhang Z., Zhang F. 2023. A novel probe set for the phylogenomics and evolution of RTA spiders. Cladistics. 39:116–128.
